# Blood host preferences and competitive inter-species dynamics within an African malaria vector species complex inferred from signs of animal activity around aquatic larval habitats

**DOI:** 10.1101/2024.08.04.606513

**Authors:** Katrina A. Walsh, Deogratius R. Kavishe, Lily M. Duggan, Lucia J. Tarimo, Rogath V. Msoffe, Elisa Manase, Nicodem J. Govella, Markus P. Eichhorn, Emmanuel W. Kaindoa, Fidelma Butler, Gerry F. Killeen

**Affiliations:** School of Biological Earth & Environmental Sciences, University College Cork, Cork, Republic of Ireland; Environmental Research Institute, University College Cork, Cork, Republic of Ireland; Department of Environmental Health and Ecological Sciences, Ifakara Health Institute, Off Mlabani Passage, Ifakara, Morogoro, Tanzania; College of Forestry, Wildlife and Tourism, Sokoine University of Agriculture, Morogoro, Tanzania; Nyerere National Park, Tanzania

## Abstract

The dual-specialized behavioural adaptions of *Anopheles arabiensis*, to feed readily upon either people or cattle, enable it to thrive and mediate persistent residual malaria transmission across much of Africa, despite widespread use of long-lasting insecticidal nets (LLINs). Indeed, LLIN scale up has often resulted in it dominating the more efficient but vulnerable malaria vector *Anopheles gambiae*, its sibling species within a complex of the latter name. However, the feeding behaviours and competitive relationships of *An. arabiensis* with other sibling species in well conserved natural ecosystems, where its known preferred hosts are scarce or absent, remain largely unexplored.

Potential aquatic habitats were surveyed for *An. gambiae* complex larvae across a gradient of natural ecosystem integrity in southern Tanzania, encompassing fully domesticated human settlements, a partially encroached Wildlife Management Area (WMA), and well conserved areas of the recently gazetted Nyerere National Park (NNP) before substantive development of tourist access or accommodation. Direct observations, tracks, spoor and other signs of human, livestock or wild animal activity around these water bodies were recorded as indirect indicators of potential blood source availability.

While only *An. arabiensis* was found in fully domesticated ecosystems, its non-vector sibling species *An. quadriannulatus* occurred in conserved areas and dominated the most intact natural ecosystems. Proportions of larvae identified as *An. arabiensis* were positively associated with human and/or cattle activity and negatively associated with distance inside NNP and away from human settlements. Proportions of *An. quadriannulatus* were positively associated with activities of impala and bushpig, implicating both as likely preferred blood hosts. While abundant impala and lack of humans or cattle in intact acacia savannah within NNP apparently allowed it to dominate *An. arabiensis*, presence of bushpig seemed to provide it with a foothold in miombo woodlands of the WMA, despite encroachment by people and livestock. While this antelope and suid are essentially unrelated, both are non- migratory residents of small home ranges with perennial surface water, representing preferred hosts for *An. quadriannulatus* that are widespread across extensive natural ecosystems all year round.

Despite dominance of *An. quadriannulatus* in well-conserved areas, *An. arabiensis* was even found in absolutely intact environments >40km inside NNP, suggesting it can survive on blood from one or more unidentified wild species. While self-sustaining refuge populations of *An. arabiensis* inside conservation areas, supported by wild blood hosts that are fundamentally beyond the reach of insecticidal interventions targeted at humans or their livestock, may confound efforts to eliminate this key malaria vector, they might also enable insecticide resistance management strategies that could restore the effectiveness of pyrethroids in particular.

## Introduction

Historically, *An. gambiae sensu stricto (s.s.)* has been the most notorious vector of malaria across Africa and is the nominate species of the *An. gambiae* complex, often referred to as *An. gambiae sensu lato (s.l.)* comprised of eight other sibling species [1, 2] This species complex is ubiquitous across sub-Saharan Africa and includes three of the four most important vectors across the continent, two of which are widespread across eastern Africa: While *An. gambiae s.s.* dominates moister parts of the region, it’s sibling species *An. arabiensis* does so across extensive arid areas [3–7]. However, the feeding behaviours and competitive relationships of *An. arabiensis* with other sibling species in well conserved natural ecosystems, where its known preferred hosts are scarce or absent, remain largely unexplored. Improved understanding of the major population dynamics drivers for this key malaria vector species, including any competitive interactions with sympatric sibling species that may occur [8] will be essential for devising more effective strategies to control [9,11] or even eliminate [12] them.

Members of the *An. gambiae* complex each exhibit distinctive behavioural preferences for particular mammalian species as blood sources, which in turn affect their vectorial capacity and the effectiveness of vector control interventions targeting those free flying adult mosquitoes. For example, *An. gambiae s.s.* exhibits a very strong, almost singular preference for human blood, so it can mediate exceptionally high rates of malaria transmission [10,13]. However, such human-specialized anthropophagic traits also leaves *An. gambiae s.s.* vulnerable to insecticidal vector control interventions for protecting sleeping humans [12, 14], like indoor residual spraying (IRS) and long-lasting insecticidal nets (LLINs). As a result of widespread uptake of LLINs and IRS measures, both of which specifically target mosquitoes that attack sleeping humans inside their houses, populations of *An. gambiae s.s*. have declined across much of their former natural range and have even been effectively eliminated from large parts of east Africa [12, 15–19].

However, local elimination of *An. gambiae s.s.* across large tracts of Africa coincided with increased relative abundance of its morphologically indistinguishable sibling species *An. arabiensis*, resulting in a decisive shift in species composition within the complex over a remarkably brief period of a few short years [12, 15–17]. The subsequent dominance of *An. arabiensis* arises from a suite of phenotypically plastic behavioural traits that allow this far less exclusively human-dependent vector to minimize its exposure to the active ingredients of LLINs and IRS [20] thereby mediating persistent residual malaria transmission [14, 21–23].

While *An. gambiae s.s.* feeds almost exclusively on human blood, *An. arabiensis* also feeds readily upon cattle [24–26]. Furthermore, as they do not appear to acquire bloodmeals from any other animal in domesticated landscapes, *An. arabiensis* seemingly demonstrate strong specialization upon these two hosts and can flexibly feed on either or both of them, with blood meal choices being highly dependent upon the relative availabilities of cattle and humans across scales as fine as a few metres [27, 28].

These host choice behaviours makes it particularly difficult to control *An. arabiensis*, not only because feeding on cattle makes this species less dependent upon human blood [10, 14, 21, 29], but also evolutionary adaptation to feeding upon animals appears to be associated with outdoor-biting, outdoor resting, early-exiting and crepuscular feeding behaviours [10, 14, 21, 29], that allow it to evade exposure to insecticidal LLIN and IRS even when it does encounter humans [10, 14, 20, 31]. Synergy between all these evasive behaviours make *An. arabiensis* notoriously resilient against attack with LLINs and IRS [20, 31], and therefore a crucially important vector of persistent residual malaria transmission [10, 14, 21].

Having said that, *An. arabiensis* may also be able to escape pesticide pressure entirely by living in places where humans, livestock and agriculture are essentially absent, although the evidence to date remains ambiguous. While the known host preferences [24–26, 28] and associated evasive behaviours exhibited by *An. arabiensis* [10, 20] are consistent with adaptation to domesticated land use patterns [32], particularly in arid areas [3, 5–7, 33, 34] where pastoralism is often the predominant livelihood [35], *An. arabiensis* populations have also been identified inside African conservation areas. For example, adult *An. arabiensis* mosquitoes were caught during a study inside Mana Pools National Park, Zimbabwe [36], although these experiments were completed at a permanent research camp with a small resident population of cattle. Similarly, while an *An. arabiensis* population has been documented at the Malahlapanga springs inside Kruger National Park in South Africa [37–39], and it has been suggested that this malaria vector may be surviving on blood meals from wild animals rather than their known preferred human and cattle hosts [37] , this survey site was <20 km from the park boundary and adjacent villages. It therefore remains unclear whether *An. arabiensis* can exploit alternative hosts in wild areas, potentially creating a population-stabilizing portfolio effect [40] that sustains refuge populations far away from vector control interventions and other anthropogenic pesticide pressures.

If indeed such wild refugia populations of *An. arabiensis* do exist inside conserved wilderness areas, far away from insecticide selection pressure, they might well retain original wild type traits for susceptibility to public health insecticides, particularly pyrethroids. Physiological resistance to the pyrethroids, arising from near-exclusive reliance upon this exceptionally useful insecticide class for LLINs and IRS, has clearly undermined progress to date and now threatens a catastrophic resurgence of malaria-related morbidity and mortality [41–49]. The persistence of such genetically diverse source populations of *An. arabiensis* inside conserved wilderness areas could therefore represent an invaluable form of natural capital [50–52], which could potentially be exploited by astute, long-overdue pre-emptive insecticide resistance management strategies [49,53].

The goal of this study was therefore to investigate aquatic larval habitat occupancy by *An. arabiensis* across an environmentally heterogenous study area in southern Tanzania with highly variable availabilities of diverse mammalians hosts (Figure 1 and S5 Appendix), in order to determine whether this key vector of residual malaria transmission [10, 14, 20, 21, 23] does indeed survive in such wild refugia by feeding upon one or more species of wild mammal. Fortuitously, this study also revealed the presence of *An. quadriannulatus,* another zoophagic, arid-adapted member of the *An. gambiae* complex of negligible relevance to malaria transmission [6, 7, 24, 25, 34, 54, 55] co-existing with *An. arabiensis* inside conserved natural ecosystems, in a finely-balanced competitive relationship that was heavily influenced by fine-scale variations in the comparative availabilities of humans, livestock and particular wildlife species.

**Figure 1.**
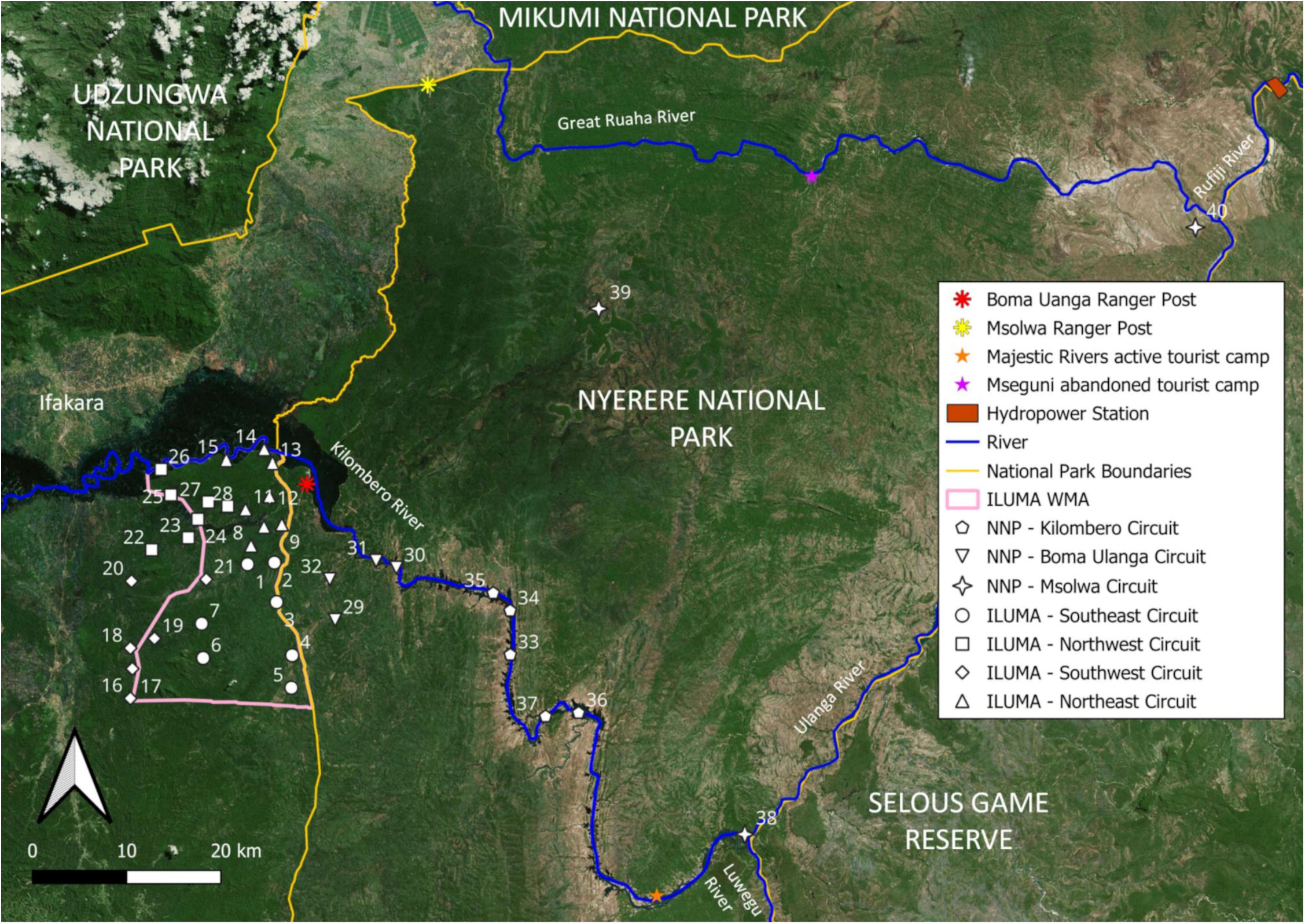
Map of the environmentally heterogeneous study area, ranging from fully domesticated human settlements just outside the western border of the Ifakara-Lupiro- Mangula Wildlife Management Area (ILUMA WMA. [56–58]**) through to well conserved natural ecosystems within Nyerere National Park (NNP) to the east of it, displaying the distribution of suitable camping locations used as the sampling frame for all the surveys described in this report (S5 Appendix, table S5.3).**

Each of the 32 camps detailed in table S5.3 are illustrated in the geographic context of the boundaries of the Ifakara-Lupiro- Mangula Wildlife Management Area (ILUMA WMA) and Nyerere National Park (NNP), with camp number 1 being the central main camp Msakamba (S5 Appendix).

## Results

### Aquatic habitat characteristics and occupancy by An. gambiae complex larvae

For a comprehensive, unabridged analysis and interpretation of all the data collected on aquatic habitat characteristics and occupancy by mosquito larvae, see S1 Appendix. In summary, a total of 8,266 dips across 1,944 potential larval habitats were completed over the sampling period from January 2022 to July 2023 (Table S1.1). Larvae from the genus *Anopheles* were present in a total of 1,058 habitats, out of which 72.3% were found to be occupied by *An. gambiae* complex larvae, representing an overall occupancy rate of 39.4% across all habitats (Table S1.1).

Ecosystem integrity, survey round, habitat type, habitat perimeter, short vegetation and the number of dips were all significant predictors of *An. gambiae* complex larval occupancy (Table S1.2). Ecosystem integrity was the third most important predictor of larval occupancy, indicating a significant decrease in occupancy rates between aquatic habitats in fully domesticated ecosystems and aquatic habitats intact natural ecosystems (Table S1.2), suggesting that larval occupancy was somewhat lower for aquatic habitats in well conserved areas further away from human settlements and domesticated land use (Figure 2). Despite this, *An. gambiae* complex larvae, were nevertheless present with mean occupancy rates exceeding 25% in even absolutely intact environments (Figure 2), confirming that *An. gambiae* complex adults were ovipositing all year-round in aquatic habitats >40km away from the nearest resident human or livestock hosts.

**Figure 2.**
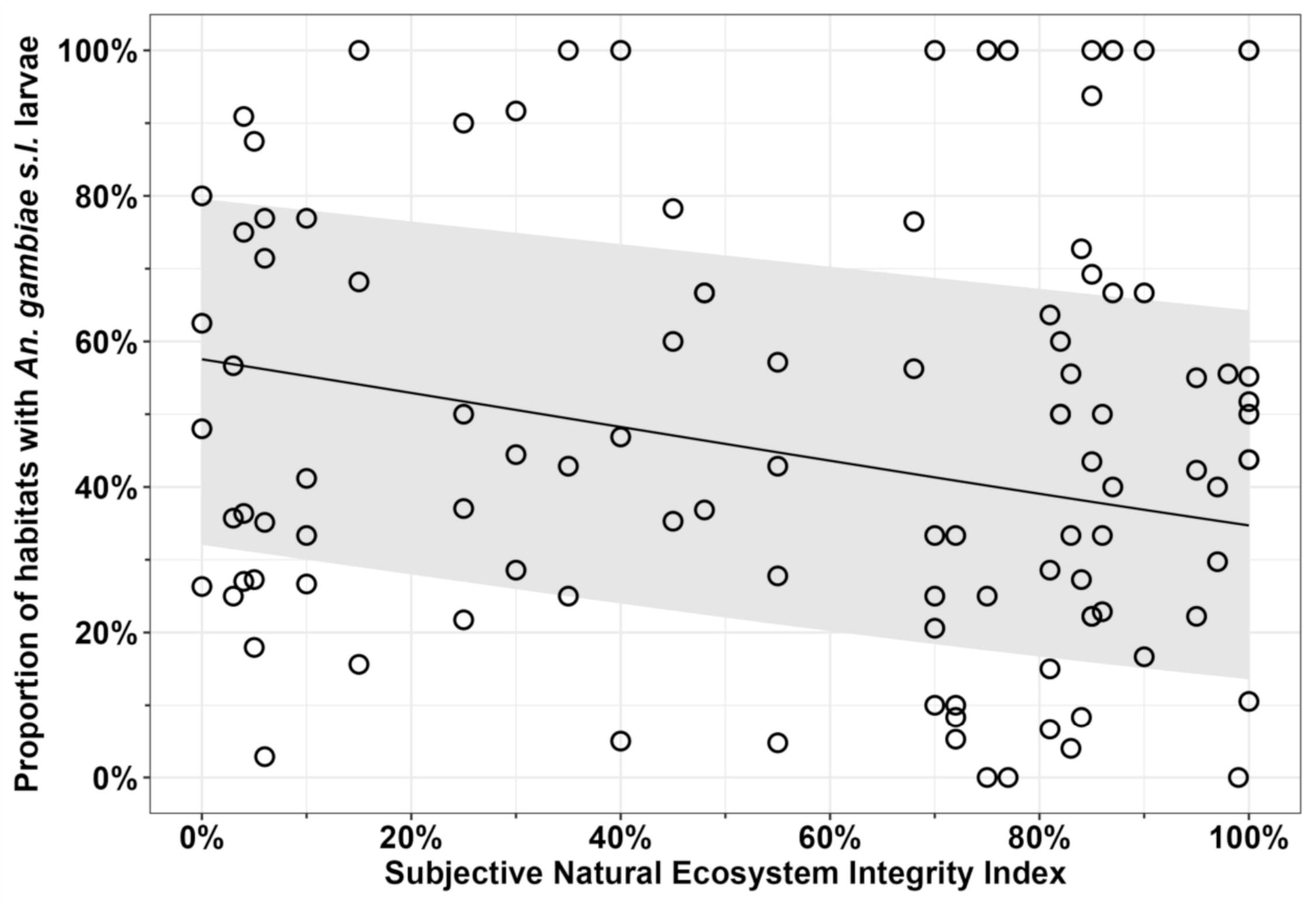
The proportion of habitats occupied by *An. gambiae* complex larvae for each survey conducted over four rounds plotted as a function of ecosystem integrity.

The estimated mean trend with 95% confidence intervals reflects the predicted values for each surveyed camp location based on a fitted GLMM identical to the multivariate model detailed in Appendix 1 (Table S1.2), except that all fixed effects other than the subjective natural ecosystem integrity index (SNEII) were instead treated as random effects, so that the predictions reflect the absolute means rather than the means for the reference values of each variable.

### Competitive co-existence of An. gambiae complex sibling species Anopheles arabiensis and Anopheles quadriannulatus

Polymerase chain reaction (PCR) results from the successfully amplified *An. gambiae* complex F0 adults that were raised from collected larvae during surveys in the first three survey rounds in 2022, demonstrated that these *An. gambiae* complex populations were not comprised of only *An. arabiensis*, but also included large numbers *An. quadriannulatus*. These two sibling species often were found co-existing alongside each other in the same habitats on the same day (Figure 3).

**Figure 3.**
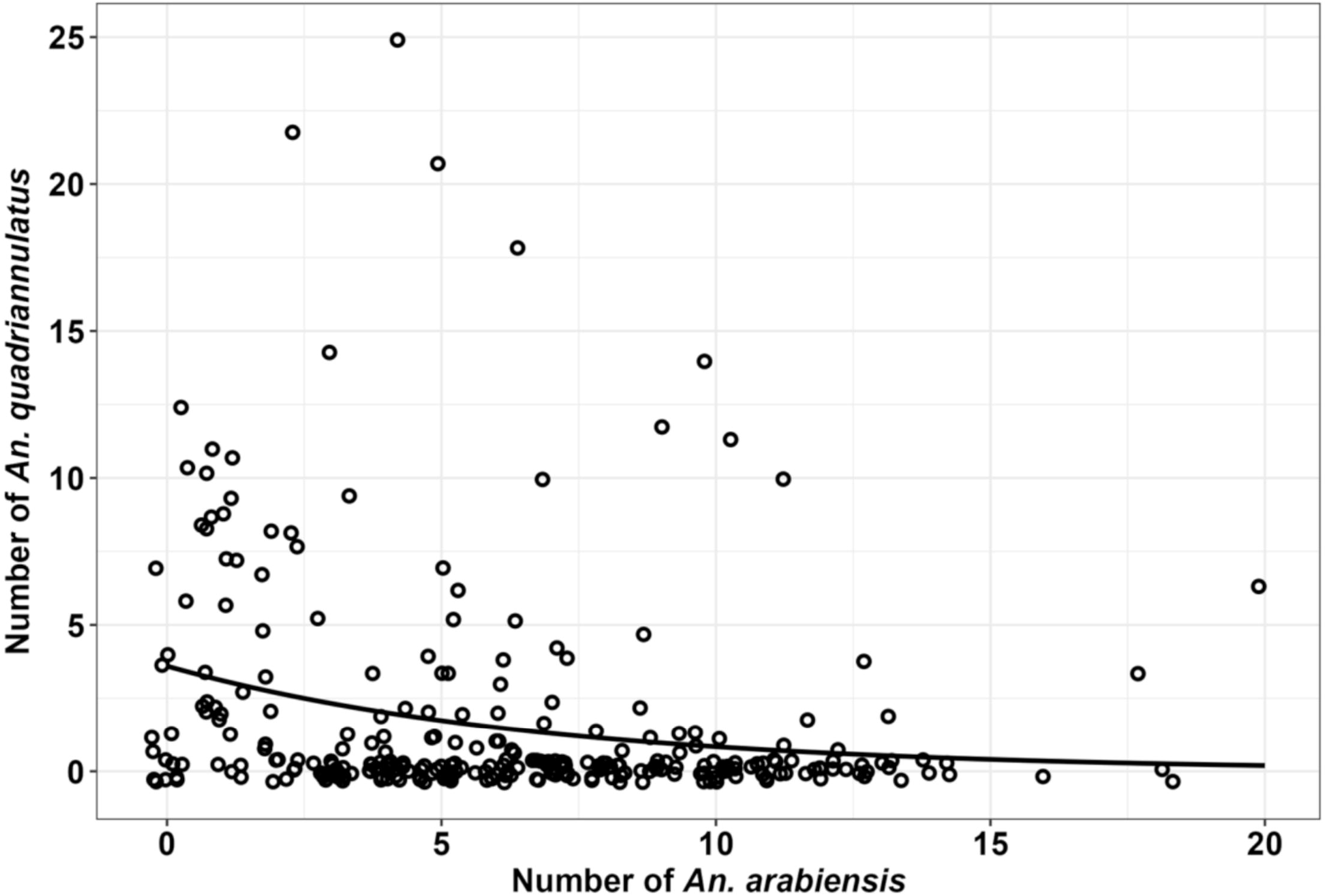
The number of *An. arabiensis* larvae per habitat plotted against the number of *An. quadriannulatus* larvae per habitat, as determined during the full fourth round of surveys.

Although variable species composition within the *An. gambiae* complex could have been explained by independently varying densities of the two sibling species found within the study area, this did not appear to be the case: As illustrated in Figure 3, the absolute numbers of *An. arabiensis* identified at each habitat were negatively correlated (ρ = -0.31, p = <0.0001) with the absolute numbers of *An. quadriannulatus* found. In simple terms, wherever there were more *An. arabiensis,* there tended to be less *An. quadriannulatus*, and *vice versa*, confirming that these two species appear to compete with each other in the strict sense (Box 1) to at least some extent.

#### Box 1. Defining competition, competitive displacement and competitive co-existence in strict ecological terms

*Competition* occurs when two species require the same limiting resources [59, 60]. The superior competitor reduces the availability of resources, mainly through exploitative or interference mechanisms [61,62], thus weakening the survival and reproductive success of the latter competitor. Therefore, in a competitive relationship between two species, the relative abundance of one species is affected by the abundance of its competitor [63,64], just as illustrated for *An. arabiensis* and *An. quadriannulatus* in Figure 2. The most extreme outcome of competition is known as *competitive displacement* [65], when one species exhibits such a clear competitive advantage that its dominance leads to the elimination of the other. Hence, this concept is based on the principle of competitive exclusion [66] wherein it is assumed, sometimes too simplistically, that two species can only co-exist if they compete for different limiting resources [66,67].

However, complete competitive displacement is rarely observed between naturally co-occurring species that seemingly require the same resources. A famous example is that of the ‘paradox of the plankton’, where Hutchinson [68] raises the question as to why there is such diversity of co-existing plankton, despite apparently requiring the same limiting resources. Indeed, although such *competitive co-existence* sounds counter-intuitive, it can be explained by stochastic dynamics driven by external environmental fluctuations [68,69] as well as intrinsic chaos [70,71] and meso-scale spatial heterogeneity [72], so that ecosystem equilibrium is never reached and no single species can dominate [68].

Similarly, *Anopheles* populations are often found in sympatry [25], with aquatic stages sharing the same habitats [25, 73–75], where larval stages need to compete for limiting resources such as nutrients and space [76–80]. While *Anopheles* larval habitats certainly exhibit the kind of stochastic dynamics considered to enable competitive co-existence of two or more species [68, 81, 82], it is also worth noting that the competitive balance between species within larval populations is clearly influenced by the local abundance [73,83] and safe accessibility [16, 84–87] of suitable blood hosts for the relevant adult populations to feed upon. The influence of such factors, particularly the safe accessibility of blood meals, on the competitive dynamics and population composition of co-existing mosquito species are considered further in the discussion and are emphasized by Lounibos (2007) [8]. Indeed, Lounibos (2007) [8] concludes that the dominant competitor reduces the population abundance of the weaker competitor mainly through larval resource or interference competition mechanisms, but also that complete displacement [65] between competing mosquito species does not tend to occur. Empirical field observations of sympatric populations of more than one mosquito species in most locales, even though they clearly share habitats and resources therein, therefore, also deviates from the classical but simplistic principle of competitive exclusion [66,67], and instead resembles competitive co-existence like that observed between plankton species [68].

### Effects of distance, ecosystem integrity and landcover on An. gambiae complex species composition

The surprising PCR results also demonstrated that while only *An. arabiensis* were found in fully domesticated and highly encroached locations, *An. quadriannulatus* were found in camps with Subjective Natural Ecosystem Integrity Index (SNEII; see [56,58, S6 Appendix]) scores of 50% or higher and were most common in the best conserved locations (S2 Figure), suggesting that this non-vector utilizes wild animal hosts for blood feeding. The relative abundance of *An. arabiensis* declined as SNEII increased and the numbers of cattle herds detected became fewer (S2 Figure), suggesting that the outcome of its competitive interactions with *An. quadriannulatus* may be influenced by the availability of this known preferred host for the former species [25,73–74]. However, further investigation was required to address concerns about the potential for transportation of larvae to the central insectary and subsequent procedures for rearing them to adults distorting sample species composition, and to find out whether *An. quadriannulatus* completely replaces *An. arabiensis* deeper inside NNP, even further away from humans and livestock.

The protocol for collecting field identified mosquitoes was therefore revised (S3 Appendix) and validated (S4 Appendix), so that additional samples of larvae were preserved *in situ* and nine extra camps much further inside NNP were added for the fourth and final survey round in 2023 (Figure 1, S5 Appendix). Using these samples of larvae from round four that were identified and preserved at the point of collection in the field, distance from the NNP boundary or nearest human settlement inside the park, SNEII, and historical landcover (S6 Appendix) were all found to be determinants of the sibling species composition of the *An. gambiae* complex by univariate analysis (Table 1). However, historical landcover type was excluded from the final multivariate model (Table 1), possibly because it correlated with the distance variable. For example, acacia savanna was exclusively found inside NNP and was the dominant landcover type in the most isolated parts the park along the Kilombero river. Despite the visually obvious correlation between distance inside the park and ecosystem integrity (Figure 4), the SNEII still accounts for additional variance that is not already captured by distance. The best fit GLMM of these spatial and environmental covariates indicate that larval habitats located closer to human settlement and at locations with low ecosystem integrity scores, have higher proportions of *An. arabiensis* rather than *An. quadriannulatus* (Table 1, Figure 4). As larval habitats are located further away from the boundary and in areas of improved ecosystem integrity, the inverse effect occurs so that the odds of *An. gambiae* complex larvae amplifying as *An. arabiensis* decline by 63% for every 10km further past the boundary and by 5% for every percentage point increase in the SNEII (Table 1, Figure 4), suggesting that *An. quadriannulatus* may have a competitive advantage in fully intact natural ecosystems far away from human settlements and their associated cattle herds.

**Figure 4.**
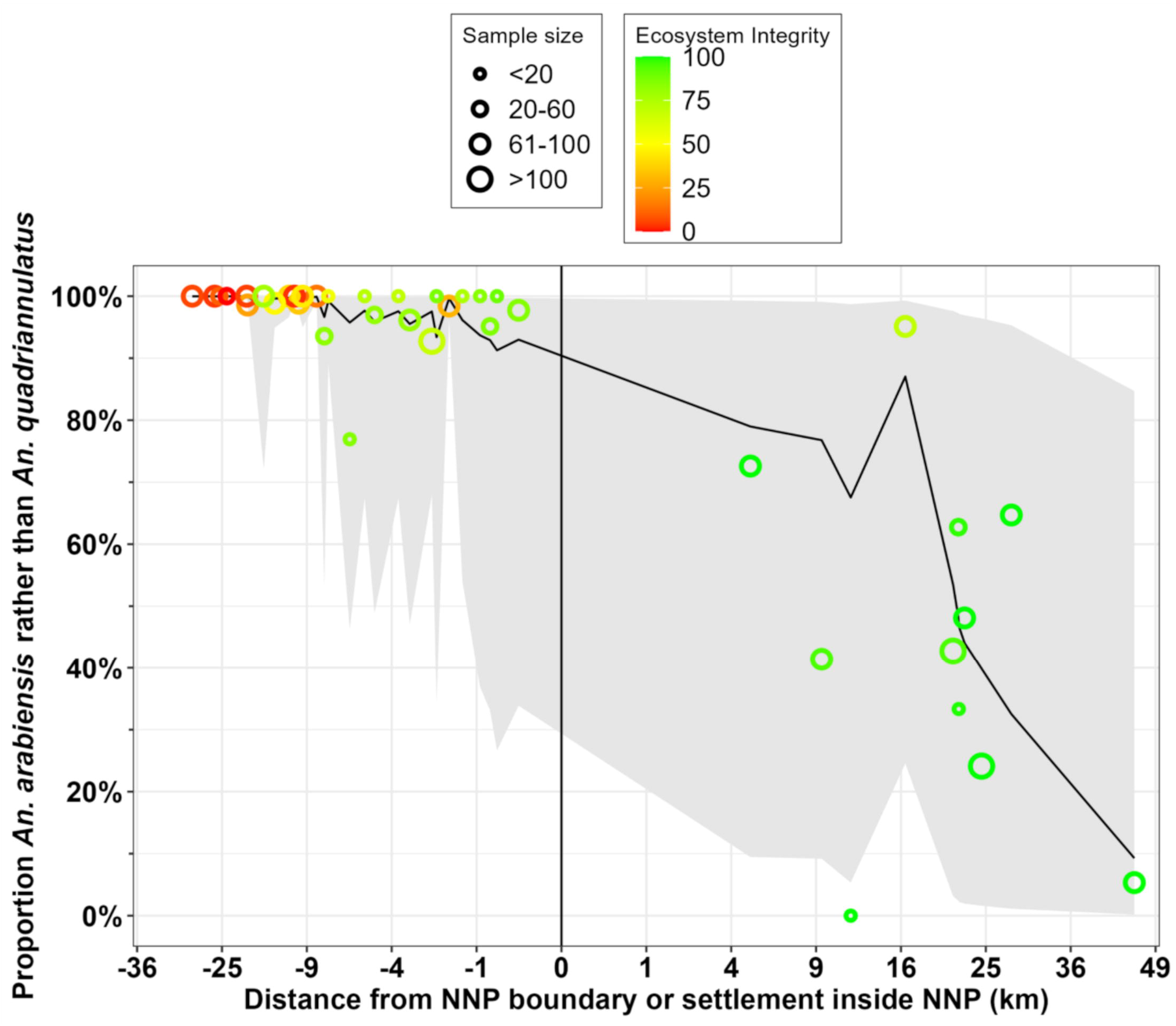
The proportion of field-identified *An. gambiae* complex collected and preserved *in situ* in the fourth round with the revised protocol (S3 Appendix and S5 Appendix) that were identified as *An. arabiensis* rather than *An. quadriannulatus* by PCR (88) plotted against distance to the nearest boundary of Nyerere National Park (NNP), with locations outside the park indicated by negative values. Sample size indicates the number of specimens identified, collected and successfully amplified at that location, while colour indicates scores for the Subjective Natural Ecosystem Integrity Index

**Table 1.**
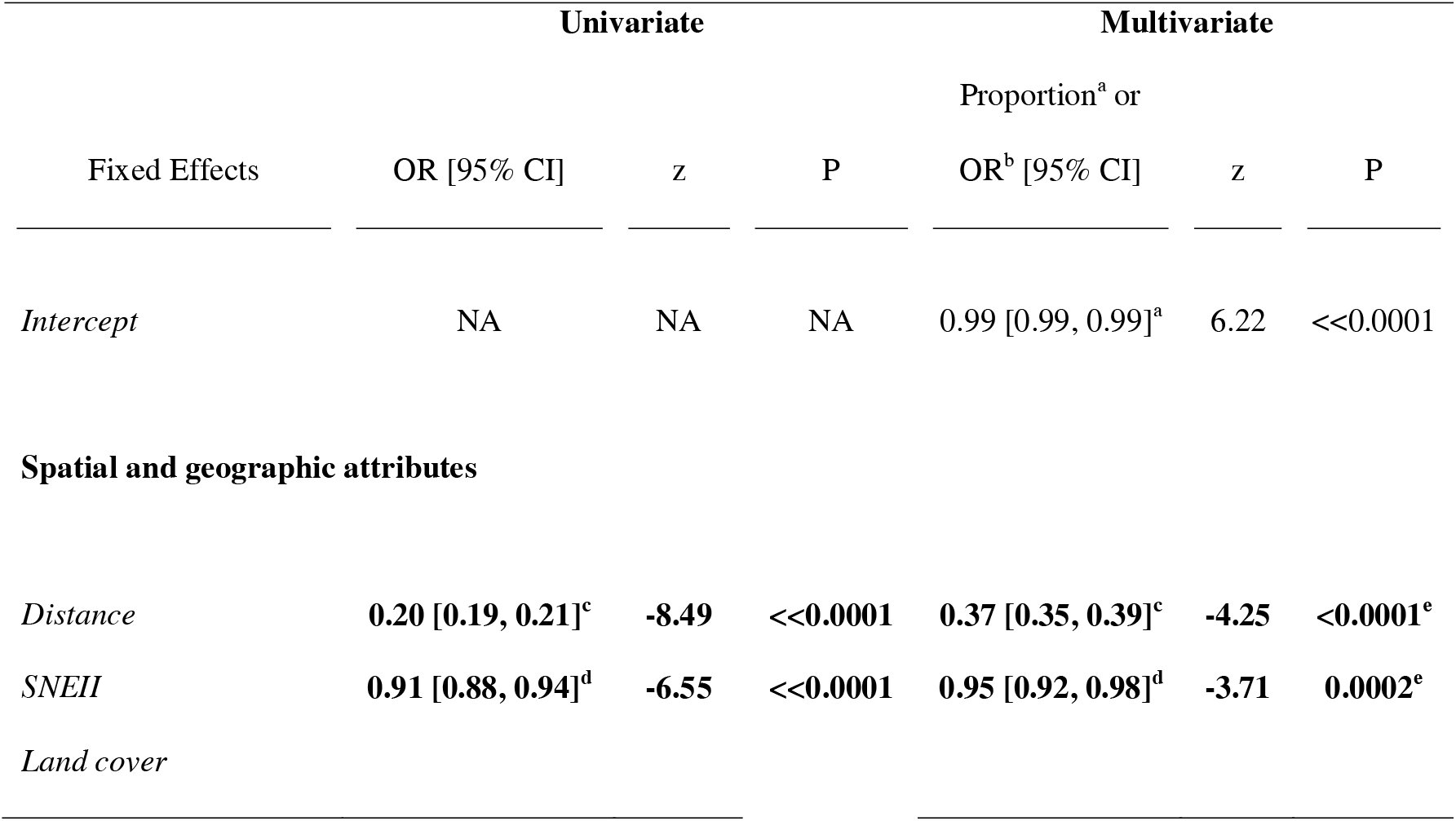
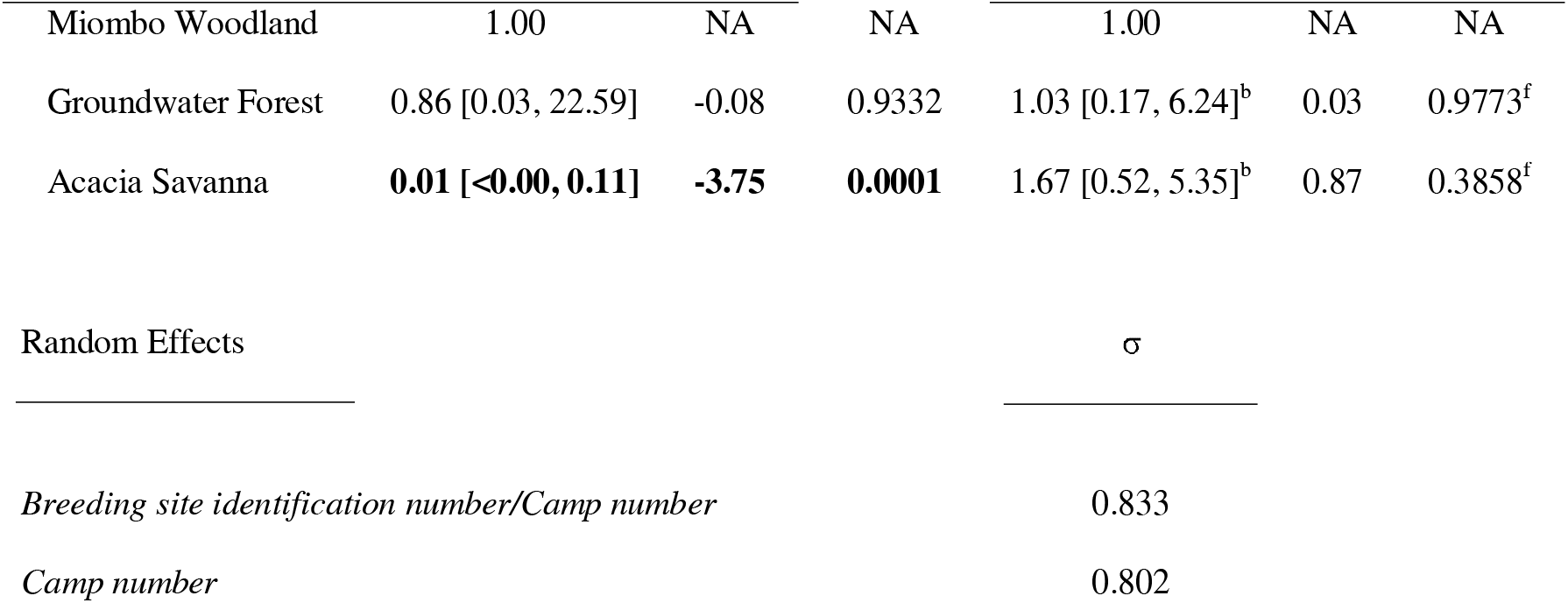
Univariate and multivariate outputs of generalized linear mixed models (GLMMs) for the effects of spatial and geographic attributes on the proportion of *An. arabiensis* rather than *An. quadriannulatus.* All model outputs were fitted to a binomial distribution with logit link function. A random effect that nested breeding site identification number within camp was included to account for variation between and covariance within larval populations. Statistically significant effects are highlighted in bold.

(SNEII [S6 Appendix]). The predicted mean values and their 95% confidence intervals for each surveyed camp location are respectively plotted as a black line and grey ribbon, and were calculated based on the estimated intercept and the fixed effects of distance and SNEII from the final multivariate model detailed in Table 1.

The map in figure 5 clearly illustrates how *An. arabiensis* dominated fully domesticated habitats around villages and all throughout ILUMA WMA, even though very low proportions (<7%) of *An. quadriannulatus* occurred in its best conserved areas, generally along the NNP boundary. A conspicuous change in species composition was observed at locations inside NNP (Figures 4 and 5), all of which were at least 5km from the nearest boundary of NNP itself or that of an adjoining part of the Selous Game Reserve or Mikumi National Park. The SNEII reaches its maximum value at most camps inside NNP (Table in S6 Appendix) and most of these had much higher proportions of *An. quadriannulatus* than any other camp in ILUMA and of the neighbouring villages to the west (Figures 4 and 5). Nevertheless, although the proportions of *An. arabiensis* generally declined with increasing distance inside NNP and away from substantial permanent human settlements (Figures 4 and 5), that geographic trend was remarkably erratic with lots of variability and wide confidence intervals (Figure 4), indicating considerable uncertainty regarding predictions of the fitted model from Table 1.

**Figure 5.**
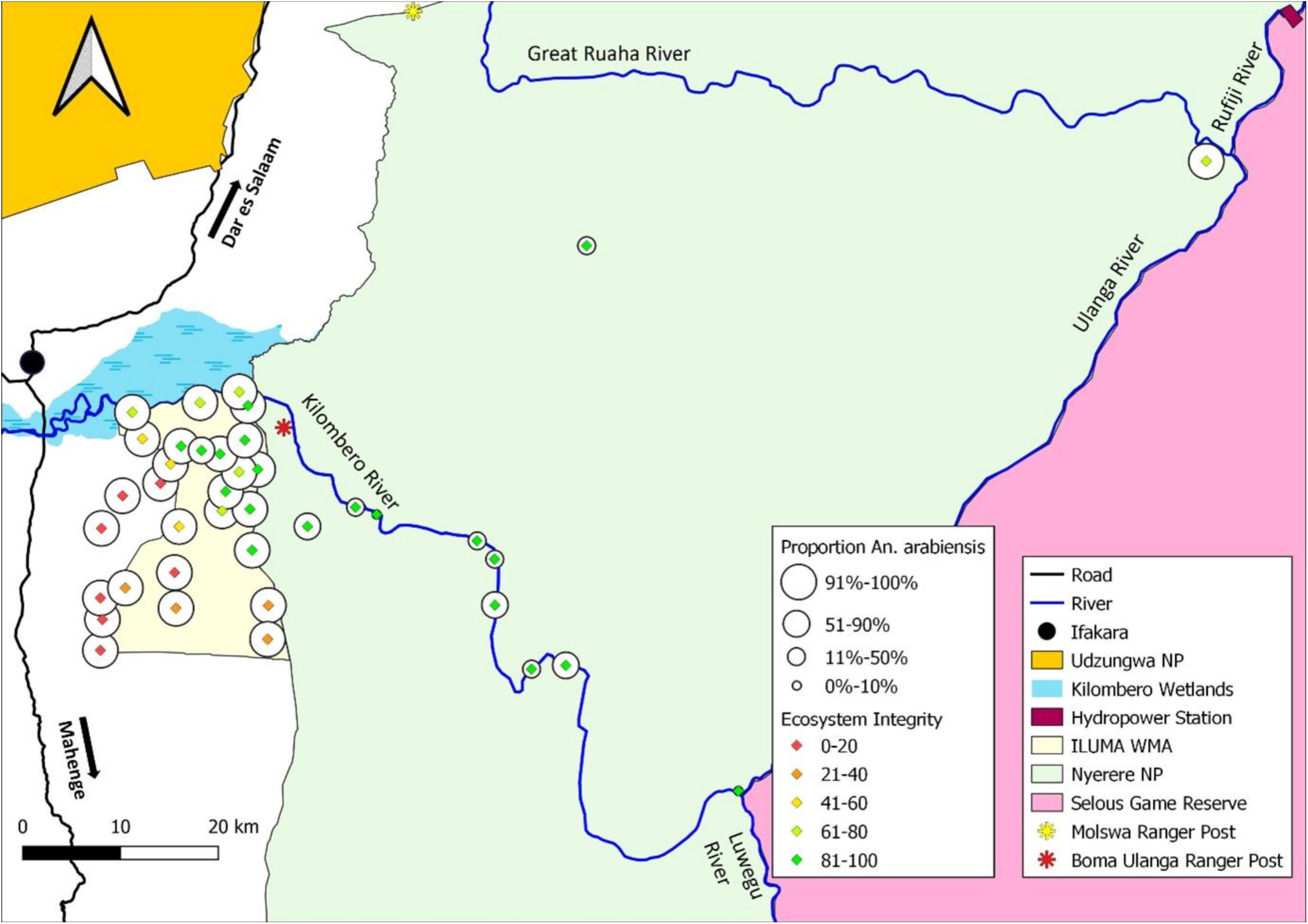
Map of ILUMA WMA and NNP illustrating how the proportion of *An. arabiensis* versus *An. quadriannulatus* varies geographically with respect to distance inside the park and away from human settlements, as well as the subjective natural ecosystem integrity (SNEII) at each camp location.

A clear outlier deep within NNP, *Bwawa la Umeme* (Camp 40), is a useful exception that proves the rule because it had the highest proportions of *An. arabiensis* (95%) and the lowest SNEII score inside the national park (Figure 5, Table in S6 Appendix). This camp was located approximately 16km from the hydropower station and a large permanent human settlement within the national park, and had also been exposed to intensive human activity such as deforestation over the previous two years, suggesting that this population may be able to readily access human blood meals.

In contrast, *Shughuli Kubwa* (Camp 38), was located 47km from the nearest boundary of NNP other than its border with the adjoining Selous Game Reserve (Figure 1), representing the furthest surveyed camp away from major human settlements. At this exceptionally well conserved location, only 5 of 76 (6%) of identified specimens were *An. arabiensis* (Figures 4 and 5), indicating a clear competitive advantage of *An. quadriannulatus* in this wild area. Nevertheless, it is notable that even in this remarkably remote, beautifully conserved location, *An. arabiensis* persisted in small numbers. Indeed, camp number 30, *Bwawa la Kiboko Zanzibar*, which had a very small sample size (n=3), was the only camp where were no *An. arabiensis* were identified (Figures 4 and 5), so it seems that this species was ubiquitous across all surveyed parts of this very well protected park.

While these data in themselves suggest that *An. arabiensis* may be able to exploit at least some wild animal species as sources of blood, and the NNP rangers could not recall ever having seen any signs of poaching or other unauthorized human activities as far downstream as *Shuguli Kubwa* (Camp 38), it is noteworthy that several small but permanently occupied ranger posts and tourist camps were scattered along the banks of the Kilombero and Ulanga Rivers, including one only 11km upstream from *Shuguli Kubwa*. While these minor outposts were all very small (Between 2 and 30 occupants at any given time), they nevertheless represented perennial sources of human blood that are within the regular flight ranges of *Anopheles* mosquitoes, relative to the sampling frame depicted in Figures 1 and 5. It was therefore not possible to rule out the possibility that the persistence of *An. arabiensis* in even the best conserved parts of the park, arises from the year-round availability of human hosts at fixed locations, albeit in a remarkably sparse and scattered population. As explained in the discussion, however, a subsequent complementary study confirmed the presence of *An. arabiensis* in the even more remote location of *Mseguni* (Figure 1), with no such small outposts of human existence nearby (Kavishe DR, personal communication). It therefore seems likely that they can survive in complete wilderness by availing of blood from species other than humans and cattle.

### Associations between An. gambiae complex species composition and potential host availability

Note, however, that the predicted means calculated from the GLMM detailed in Table 1 have wide confidence intervals (Figure 4), suggesting that the notable variability of *An. gambiae* complex species composition within the park may arise from heterogeneities in more specific factors other than distance and ecosystem integrity, such as blood resource availability and distribution. The following analysis was therefore undertaken to determine whether such variability in the competitive balance between *An. arabiensis* and *An. quadriannulatus* could be attributed to surveyable indicators of the activities of all the various mammalian species present in the study area, including humans and cattle.

Cattle herds and humans were initially examined as two distinct variables, but convergence failures during the multivariate analysis indicated that these variables were highly correlated (S7 Figure). Fortunately, both humans and cattle are known to be strongly preferred hosts of *An. arabiensis* [24,25,28], so a new variable was calculated for the combined total number of detections of people and cattle herds. This variable yielded a lower AIC value and was therefore treated as a fixed effect variable in the following univariate and multivariate analyses of the influences of the diverse potential host species upon sibling species composition within the *An. gambiae* complex.

The best fit model from multivariate analysis indicates that cattle, humans, impala and bushpig are all mammalian species that appear to influence the competitive balance between *An. arabiensis* and *An. quadriannulatus* (Table 2). Hippos and waterbuck appeared to be significant covariates in the univariate analysis but were later excluded from the multivariate models once impala was accounted for, indicating covariance between these three species, all of which were detected most frequently around waterbodies inside NNP. Similarly, common duiker was also excluded from the multivariate model once bushpig had been added, again suggesting covariance between these two species, probably arising from their common preference for moister woodlands and forests. Slender mongoose, baboons and warthog were also dropped from the multivariate analysis for similar statistical reasons, suggesting that their geographical distribution covaried with that of the species retained within the model. Of all the possible wild animal species (S8 Appendix, Table S8.8), only red duiker, wildebeest, Sharpe’s grysbok, klipspringer, African wild dog, side-striped jackal, banded mongoose, white-tailed mongoose, honey badger, African clawless otter and porcupine were not detected during radial surveys in round 4 and are therefore not represented in Table 2. Lions, hyena, and leopard were all considered to be closely associated with wild herbivore populations, albeit in complex ways, and to contribute negligibly to overall mammalian biomass (S9 Appendix), so they were not considered for inclusion in the multivariate models even though they were associated with *An. gambiae* complex population composition in the univariate analysis (Table 2).

**Table 2.**
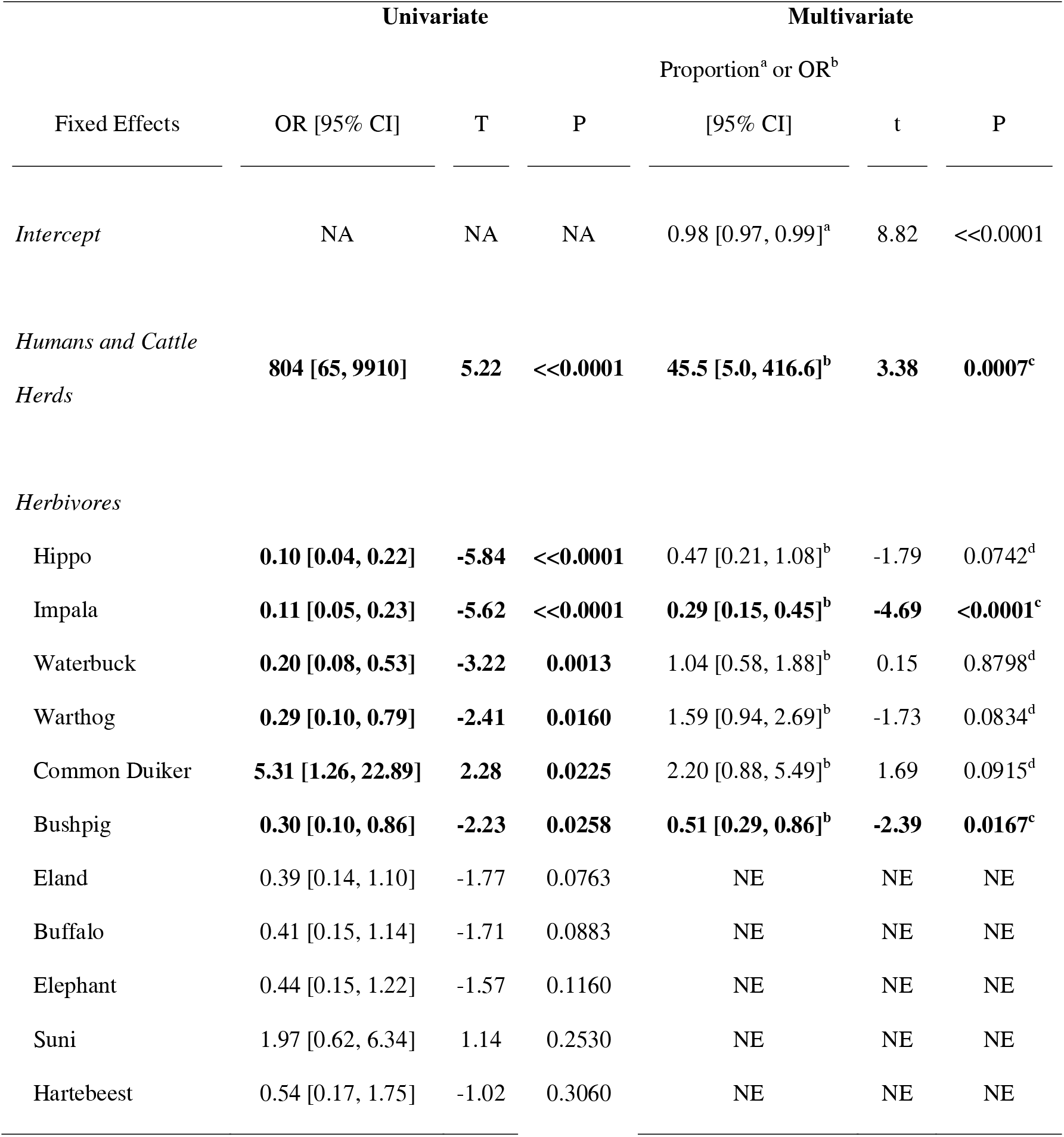
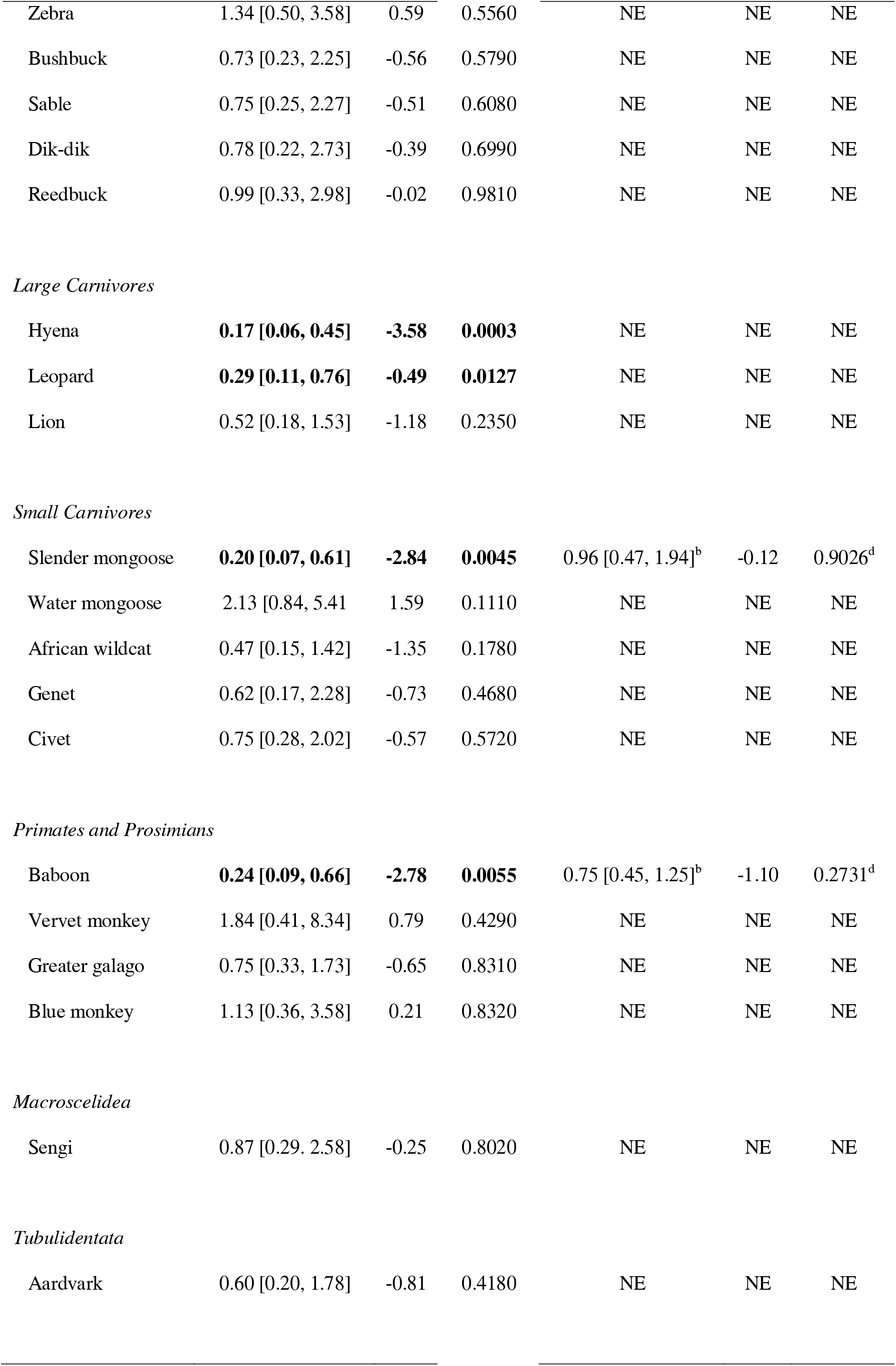
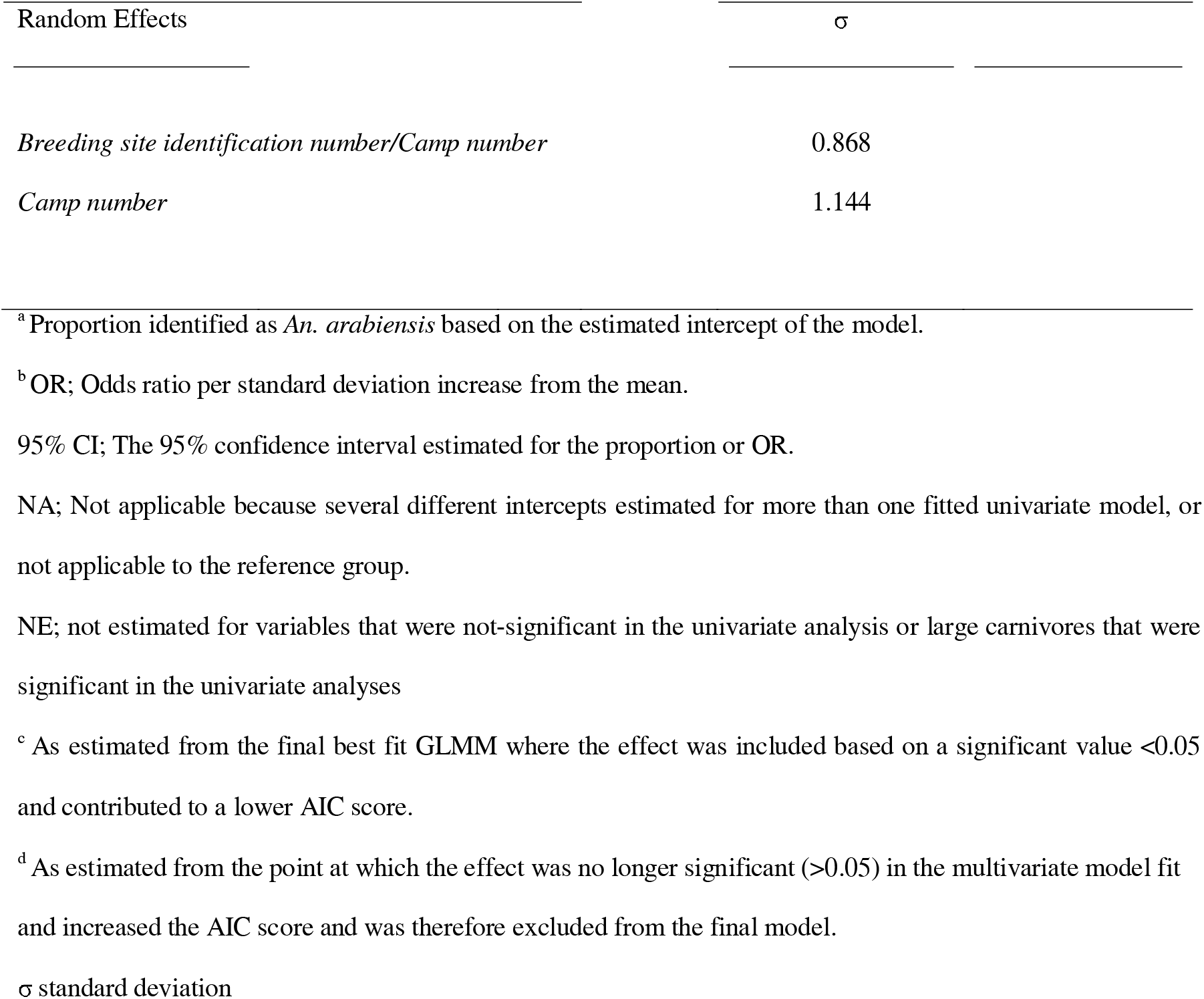
Univariate and multivariate outputs of generalised linear mixed models (GLMMs) of the effects of recorded detections of individual animal species on the proportion of *An. arabiensis* rather than *An. quadriannulatus*. All models were fitted to a binomial distribution with logit link function to the dependent variable. A random effect comprised of breeding site identification number nested within camp number was included to account for variation between and covariance within larval populations at individual waterbodies. Statistically significant effects are highlighted in bold.

Humans and cattle had the greatest effect on the relative abundances of *An. arabiensis* and *An. quadriannulatus* collected from larval habitats (Table 2). Consistent with the initial descriptive observations (S2 Figure), such a high odds ratio indicates that where cattle or humans are present, populations of *An. gambiae* complex are overwhelmingly dominated by *An. arabiensis*. When their preferred blood sources become scarce or completely absent, however, proportions of *An. arabiensis* decline, with this species seemingly losing its competitive advantage. Detections of humans or cattle were most frequently recorded in fully domesticated habitats, where the proportions of *An. arabiensis* consistently exceeded 91% (Figure 4). The highest number of detections of humans and cattle were in the villages outside the ILUMA western boundary, and these two host species were less frequently detected around larval habitats inside ILUMA. Nevertheless, humans and cattle were present, even if only in small numbers, across most of the WMA, including places that were only a kilometre from the NNP boundary. Only two camps inside ILUMA WMA, namely *Bwawa la Chamvi* and *Bwawa la Mrope* (camp numbers 9 and 13 in figure 1 and S5 Appendix), appeared devoid of these hosts (Figure 6). However, given the extent of human and livestock encroachment at the surrounding camps, the high proportions of *An. arabiensis* found at these locations were likely within flying distance of the nearest human or cow. Only one camp inside NNP (*Kambi ya Mamba*, camp number 33 in figure 1 and S5 Appendix), exhibited any signs of the presence of people (a few tracks of one or two fish poachers) and it is notable that it had higher proportions of *An. arabiensis* than most other camps inside NNP (Figure 6). This indicates that even a small number of humans may be enough to give *An. arabiensis* an advantage in a competitive relationship that they appear to dominate when allowed to. No detections of humans or cattle were found at *Bwawa la Umeme* (camp number 40 in figure 1 and S5 Appendix), which was located 16km away from a large permanent human settlement, suggesting that the high proportions of *An. arabiensis* at this location (Figure 4) may be dispersing over long distances to acquire human blood.

**Figure 6.**
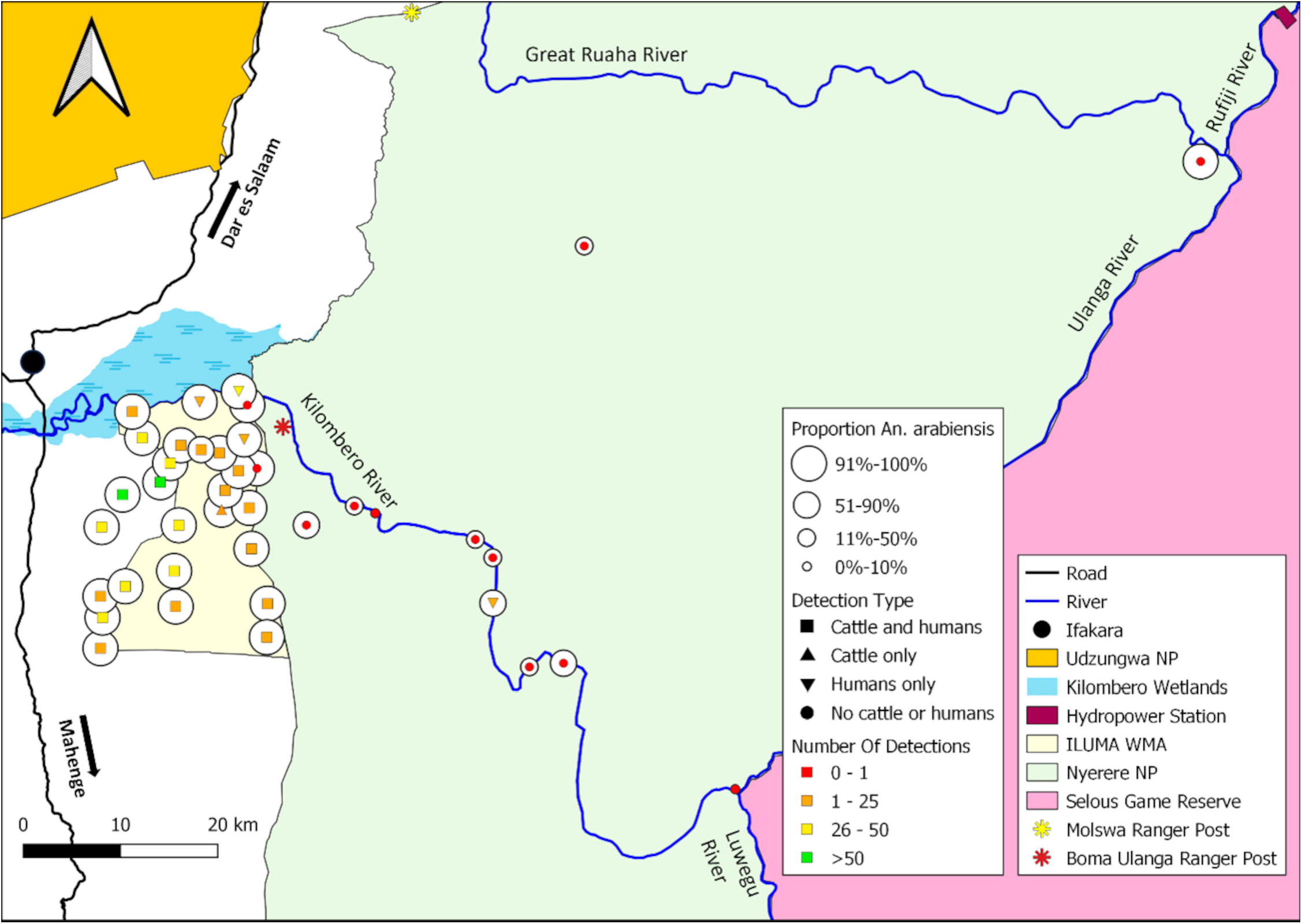
Map of ILUMA WMA and NNP illustrating how the proportion of *An. arabiensis* versus *An. quadriannulatus* varies geographically with respect to the number of times that activities of cattle and/or humans were detected at each camp.

Interestingly, the relative abundance of *An. arabiensis* was strongly and negatively associated with impala activity (Table 2). Impala were the predominant and most abundant antelope recorded inside NNP, particularly at camps along the Kilombero river where *An. quadriannulatus* were, on average, just as common as or more abundant than *An. arabiensis* (Figure 7), suggesting that this common antelope of dry savanna mosaic habitats may well be a preferred blood source for *An. quadriannulatus*.

**Figure 7.**
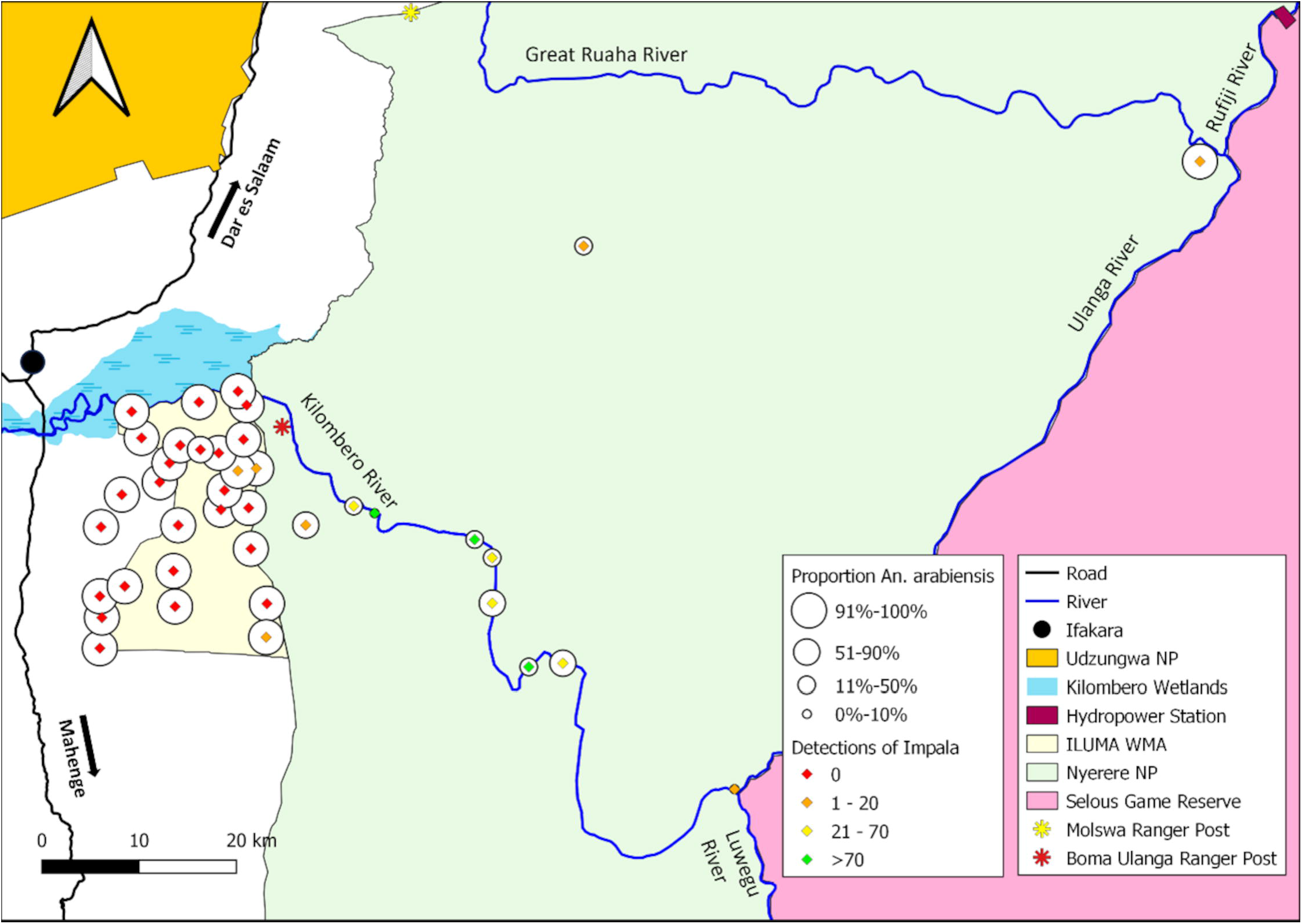
Map of ILUMA WMA and NNP illustrating how the proportion of *An. arabiensis* versus *An. quadriannulatus* varies geographically with respect to the number of times that activities of impala were detected at each camp.

The abundance of bushpig also seemed to be negatively associated with the relative abundance of *An. arabiensis* (Table 1), despite being detected far less frequently than either impala, humans or cattle, suggesting that bushpigs may provide a second preferred blood source for *An. quadriannulatus*. Bushpigs were most commonly detected in the groundwater forests of the north-eastern parts of ILUMA WMA and in the encroached areas at the interfaces between humans and wildlife (Figure 8), with the three highest number of detections occurring at *Mskamba*, *Bwawa la Nandete* and *Bwawa la Semka* (camps 1, 4 and 28 in figure 1 and S5 Appendix), all of which have moderate to high SNEII scores (S6 Appendix), and were experiencing or recovering from encroachment at the time (Table in S5 Appendix[(56, 58)]). Although only small proportions (<10%) of *An. quadriannulatus* were observed at these locations (Figure 8 and 9), they were nevertheless present despite these areas also exhibiting plenty of evidence for human and/or cattle activity (Figures 4 and 5). This suggests that bushpigs may sustain small populations of *An. quadriannulatus* that might not otherwise persist in fringe areas where humans, livestock and wildlife coexist.

**Figure 8.**
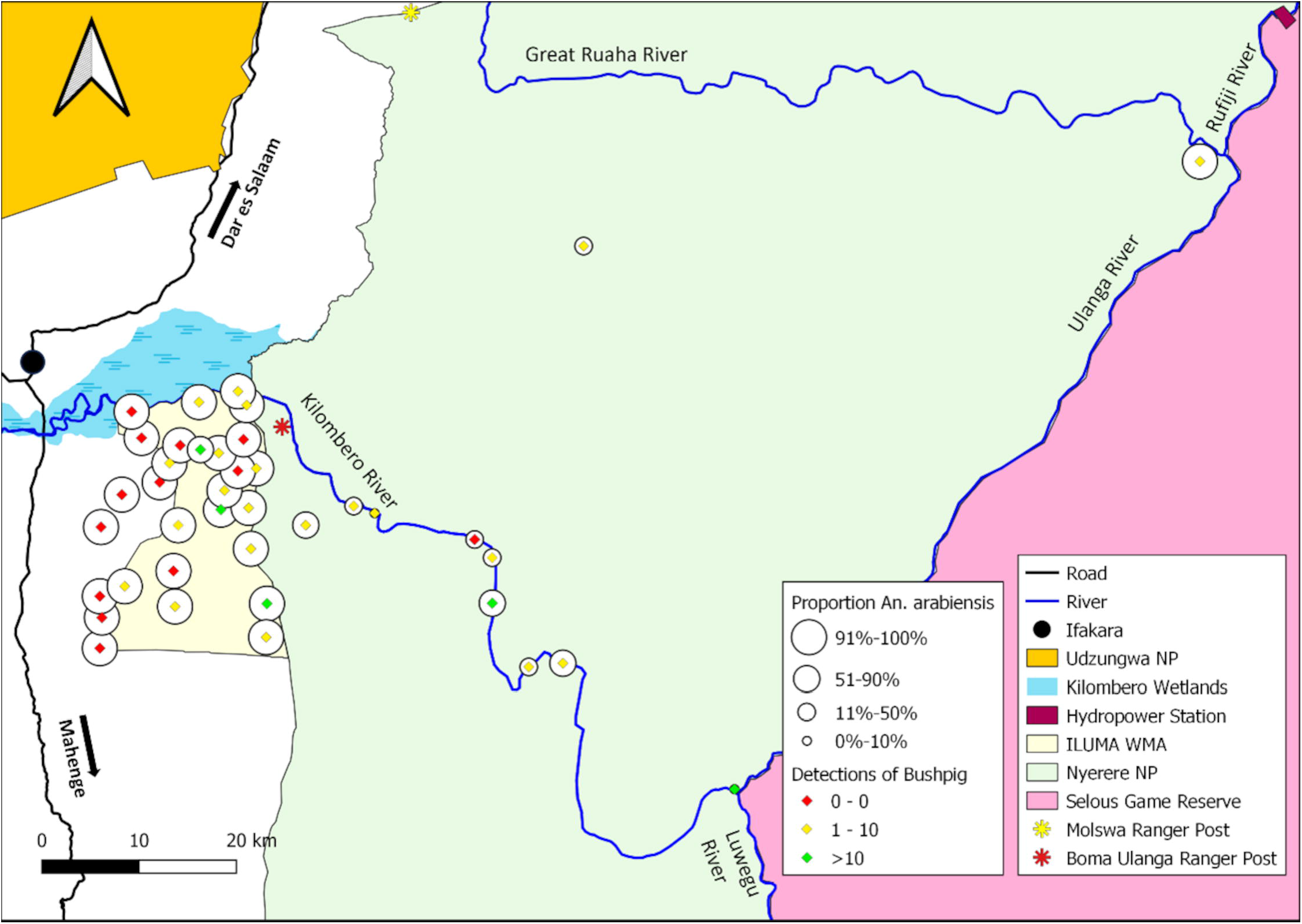
Map of ILUMA WMA and NNP illustrating how the proportion of *An. arabiensis* versus *An. quadriannulatus* varied geographically with respect to the number of times activities of bushpig were detected at each camp.

**Figure 9.**
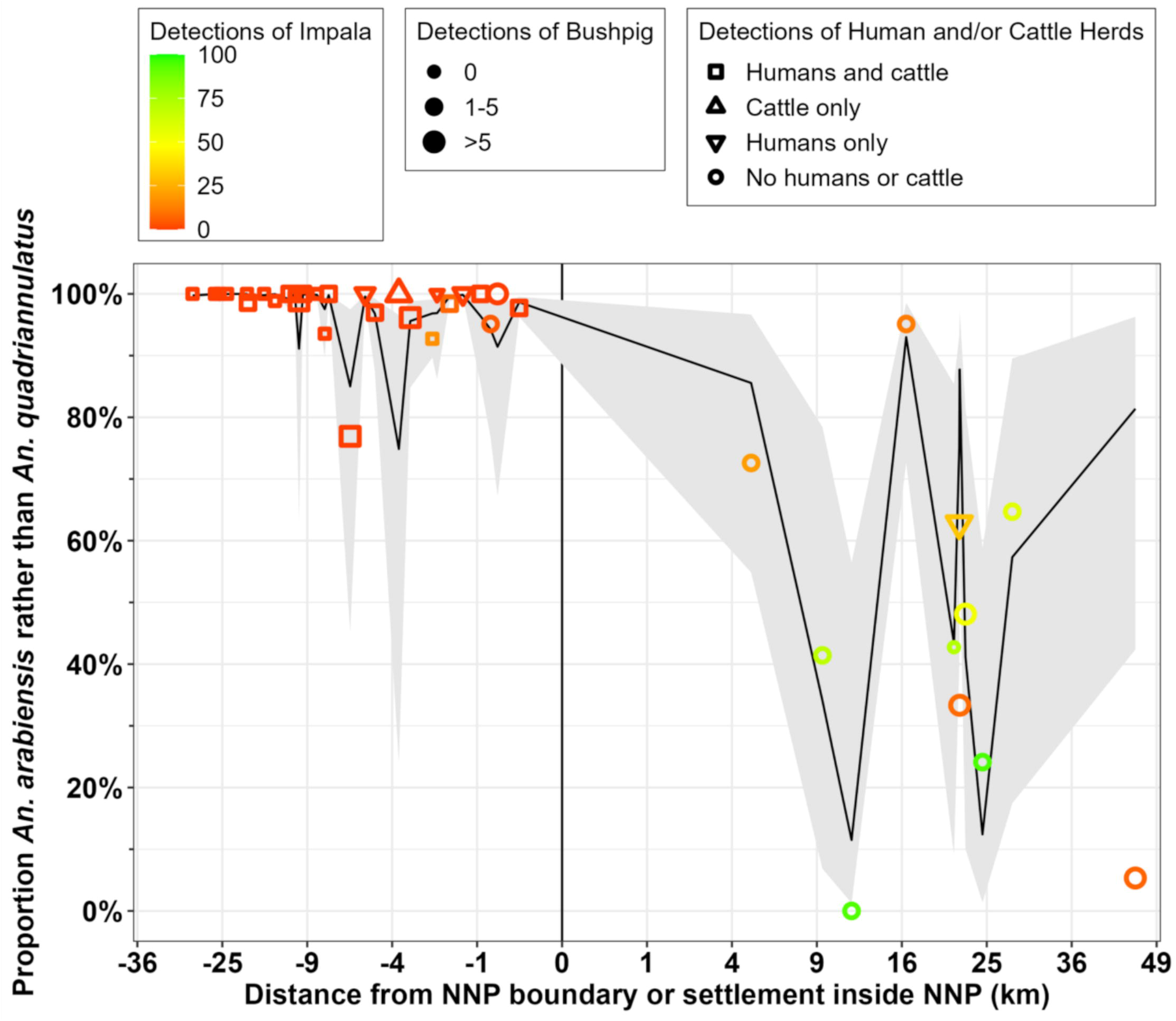
The proportion of field-identified *An. gambiae* complex collected and preserved *in situ* in the fourth round with the revised protocol (S3 Appendix and S5 Appendix) that were identified as *An. arabiensis* rather than *An. quadriannulatus* by PCR (88) plotted against distance to the nearest boundary of Nyerere National Park (NNP), with locations outside the park indicated by negative values. The size of each point indicates the number of times activities of bushpig were detected at that location, while colour indicates the number of times impala were detected. The predicted mean values and their 95% confidence intervals for each surveyed camp location are respectively plotted as a black line and grey ribbon, and were calculated based on the total number of detections for cattle and humans, impala and bushpig and the generalized linear mixed model (GLMM) presented in Table 2.

The estimated effect size for combined detections of cattle and humans on the relative abundance of *An. arabiensis* is much greater than the effect sizes estimated for impala or bushpig (Table 2), suggesting that *An. arabiensis* have a very strong competitive advantage over *An. quadriannulatus* when their preferred blood sources (cattle or humans) are present at even low densities, only allowing *An. quadriannulatus* to compete with it in wild areas far away from human settlement and livestock.

Taken at face value in mathematical terms, such a considerable effect size for humans and cattle (Table 2) would suggest that *An. arabiensis* might be expected to disappear in the complete absence of their two known preferred hosts. Instead, however, they persist at relative abundances that are generally readily detectable and usually exceed 20% deep inside NNP (Figure 9). Even at the furthest possible distance from any human settlement (Figure 9), five *An. arabiensis* larvae were identified and each were collected from a different aquatic habitat. It is highly unlikely that these five specimens collected from five separate habitats could have hatched from eggs laid by a single common mother that had fed on a person or cow outside the park and then flew almost 50km before ovipositing. It therefore seems far more likely that these specimens represent a self-sustaining wild population that feeds on one or more alternative wild hosts that have not been captured by the best fit GLMM in Table 2.

Reassuringly, the fluctuations of relative abundance between these two mosquito species across a gradient of fully domesticated to essentially intact natural ecosystems, is far better explained by data from surveys of mammalian activity (Figure 9), than by broader geographic parameters, namely distance and the SNEII (Figure 4). The calculated predicted means from the final multivariate host species model (Table 2) captures most of the considerable variability between camps within NNP and yields far narrower confidence intervals (Figure 9) than that based on geographic and ecological variables alone (Table 1 and Figure 4). The one major outlier is also informative: the model prediction also failed to capture the low proportions of *An. arabiensis* and high proportions of *An. quadriannulatus* recorded deep inside NNP at *Shughuli Kubwa* (camp number 38 in figure 1 and S5 Appendix), where there appeared to be few impala and bushpig (Figures 7 and 8), so *An. arabiensis* might be expected to predominate this location. This suggests that populations of *An. quadriannulatus* are feeding on other wild animals and that impala and bushpig are not the only hosts available to *An. quadriannulatus* in protected natural ecosystems.

## Discussion

Although no specific mammal species could be identified as a potential alternative blood source to cattle or humans that *An. arabiensis* are known to prefer, the results presented herein clearly demonstrate a competitive relationship between *An. arabiensis* and *An. quadriannulatus* where these two sibling species coexist within well conserved natural ecosystems. While *An. arabiensis* is a stereotypical vector of residual malaria transmission [10, 14, 20, 31], *An. quadriannulatus* is generally considered to be a non-vector [24, 25, 54]. This study implicates both impala and bushpig as likely preferred blood sources for the latter, relatively harmless species, allowing it to dominate the former, more dangerous, sibling species in areas where these mammals occur. Having said that, the results presented also indicates that self-sustaining refuge populations of *An. arabiensis* nevertheless exist in even the most remote, well protected conservation areas where humans and livestock are essentially absent, where they presumably survive by feeding on one or more wildlife species that remain to be identified.

### Distribution of An. quadriannulatus

At the outset of the study, *An. arabiensis* was thought to be the only persisting member of the *An. gambiae* complex in the Kilombero valley since the elimination of *An. gambiae s.s.* in this area by LLINs circa 2010 [24, 25, 54]. It was therefore assumed *a priori* that it would be the sole sibling species present, even in the best conserved corners of ILUMA WMA, by extending its ability to feed on cattle to also exploit wild bovids, African buffalo in particular. Instead, the data revealed the presence of the much more zoophagic and relatively harmless species, *An. quadriannulatus* [24, 25, 54, 55] , in conserved wilderness areas, an observation that was completely unanticipated given that this species had only once been identified over the long history of mosquito research in the Kilombero valley [89]. Nevertheless, the sympatric populations of the two sibling species reported herein are consistent with the historical literature across Africa, which indicated that *An. quadriannulatus* occur in the same geographic regions as *An. arabiensis* [24], although the distribution of the former is much patchier than the latter [24,54].

Originally identified as a separate species by Davidson (1964) [90], populations of *An. quadriannulatus* were initially reported in areas of southeast Africa including Zimbabwe, Zambia, Eswatini and South Africa [24, 25, 90]. Soon after, populations exhibiting more endophilic tendencies were discovered in Ethiopia [24] that have since been recognised as a separate species, named *An. quadriannulatus* species B at the time [91] and since renamed *An. amharcus* [1]. The patchy distribution of these two allopatric species are therefore thought to represent relics of a zoophagic, exophilic, ancestral species that was found throughout the ancient rainforests of Africa [32]. In Tanzania, records of *An. quadriannulatus* populations are almost entirely lacking. Documented originally on the island of Zanzibar [92], reports of this species seems to be lacking almost entirely. However, data from sentinel sites in 2022 and 2023 showed that *An. quadriannulatus* were present (NJ Govella, personal communication). On the mainland, records are apparently limited to a report from Mbozi in southwestern Tanzania, which is referred to by Kweka *et al.* (2020) [93] and a recent study that revealed the presence of an *An. quadriannulatus* population at Lake Manyara, an arid region of the Rift Valley in northern Tanzania [93].

In general, studies of *An. gambiae* complex are conducted to understand the ecology of the most important malaria vectors and are therefore most frequently carried out in inhabited areas, where human-biting mosquitoes of epidemiological importance thrive by exploiting abundant human populations. Although behaviourally plastic malaria vectors like *An. arabiensis* feature strongly, highly zoophagic species like *An. quadriannulatus* may be underrepresented in such studies around towns, villages and farms. As demonstrated herein, finding self-sustaining populations of *An. quadriannulatus* may require sampling of conserved wilderness areas at least 10km from the nearest human settlement. This species may therefore have a far wider distribution than previously appreciated because their apparent inability to compete with *An. arabiensis* wherever humans and cattle abound restricts them to remote, conserved wilderness areas that are rarely surveyed.

Furthermore, these observations perhaps shed light on the apparent scarcity of *An. quadriannulatus* across Africa over recent years when compared with historical records. When much of the classical literature on African *Anopheles* was written half a century ago [24,25] rural settlements across the continent were, generally speaking, fewer, smaller and frequently surrounded by natural ecosystems with healthy wildlife populations. Today, the opposite is true: Increasingly, intact wilderness is found only inside protected areas that are surrounded by growing populations of humans and livestock. It is therefore plausible that the already relict range of *An. quadriannulatus* contracted even more over recent decades.

### Competitive co-existence of An. arabiensis and An. quadriannulatus populations

The presence of *An. quadriannulatus* in varying proportions also gave insights into the potential drivers of their ability to co-exist with *An. arabiensis* in fully conserved wilderness, despite their apparently competitive relationship (See Box 1 for discussion of the general principles of competitive co-existence and a brief overview of how it relates to the larvae of sympatric mosquito populations). According to Gillies and Coetzee (1987) [25], the range of aquatic habitats used by all the various freshwater species of the *An. gambiae* complex appear to be very similar, without any obvious differences between sibling species. While some distinctions in habitat use have been documented more recently between other members of the same complex, namely *An. gambiae s.s.* and *An. coluzzii*, these are nevertheless relatively subtle [94,95]. Interestingly, the initial discovery of *An. quadriannulatus* describes wild animal footprints and shallow pools as the specific habitat types they were typically found in [96], and this is remarkably consistent with the observed characteristics of occupied habitat types found in the most remote and best conserved parts of the study area described herein (Figure S3.1F, for example).

The co-occurrence of both *An. arabiensis* and *An. quadriannulatus* in the same habitats is not surprising, as sympatric species of the *An. gambiae* complex are often found sharing the same habitat [73–75, 97]. The relative abundance of one species compared to another in shared habitats depends on their ability to compete with each other under local conditions, which in turn depends upon a variety of factors that affect the development or survival of various life stages, such as the presence of predators, food availability and larval densities [98–100]. Furthermore, environmental variability can also influence larval competitive interactions, which may lead to composition shifts over time and space [39, 98, 101].

Although the habitats surveyed during this study may appear similar, *An. arabiensis* and *An. quadriannulatus* may have subtly distinctive environmental preferences within an aquatic habitat, determined by factors like water quality and temperature that may change considerably over temporal scales ranging from hours and even minutes up to weeks and months. For example, a study in Kruger National Park, South Africa yielded evidence of a seasonal species composition shift, from *An. arabiensis* predominating in the wet season to a sudden proliferation of *An. quadriannulatus* in the dry season, and suggested that the latter could withstand the higher saline concentrations found in stagnant dry-season water bodies [39]. Furthermore this view was supported by the surprizing appearance of *An. merus*, a saltwater species from the same complex normally associated with coastal environments, at the same time [39].

Additionally, while both *An. arabiensis* and *An. quadriannulatus* are both clearly adapted to arid conditions [3, 5–7, 33, 34], the latter is considered to exhibit greater tolerance of cooler, more temperate conditions [24]. This suggests that *An. quadriannulatus* may enjoy temporary competitive advantages when weather conditions in NNP generally become cooler at the end of the rainy season and start of the dry season, from May through to July. Previous studies have shown that water temperature has considerable effects on mixed populations of *An. arabiensis* and *An. gambiae s.s.* larvae, with the forming exhibiting higher survival rates in high water temperatures exceeding 30°C [101] but outcompeted below that threshold [98,101]. This suggests that higher water temperatures and the hotter, drier environmental conditions of the acacia savannah inside NNP may also provide *An. arabiensis* with a competitive edge against *An. quadriannulatus* in these intact natural ecosystems.

Although interspecific larval competition may be a contributing factor to the fluctuating species composition observed in this study, it is also important to consider the full mosquito life cycle because each life stage is interdependent. For example, competitive interactions at larval stages influence adult population density and fitness [84, 102–106]. Conversely, however, larvae cannot occupy habitats without a female first successfully acquiring a blood meal and then finding a suitable aquatic habitat for oviposition somewhere reasonably nearby. The ability of adult females to acquire such limiting blood resources should also thereby influence larval population abundance and distribution, so it is perhaps unsurprising that the competitive balance between *An. arabiensis* and *An. quadriannulatus* larvae (Figure 3), and between other pairs of species within the *An. gambiae* complex, are so clearly influenced by the availability of preferred blood hosts nearby [73,83], and by vector control measures that protect some of those hosts against attack [4, 12, 15–17, 84–86, 107–109].

The close association of *An. quadriannulatus* with two wild animal species (Table 2, Figures 7, 8 and 9) suggests that the availability of preferred blood resources may either be a limiting or enabling factor for individual species that strongly influences mosquito population composition. And so, although *An. arabiensis* displaces *An. quadriannulatus* wherever they can access its known clearly preferred hosts (cattle and humans) [24,28], *An. quadriannulatus* does not completely displace *An. arabiensis* where these hosts are absent and impala and bushpig are present, but merely supresses their abundance. The influence of host availability on species composition ratios have previously been documented within the *An. gambiae s.l.* complex, between *An. arabiensis* and the highly human-specialised species *An. gambiae s.s.*: Charlwood and Edoh [73] found that the relative abundance of *An. arabiensis* larvae was higher in aquatic habitats that were close to cattle, whereas Minakawa *et al*., [83] demonstrated that higher densities *An. gambiae s.s*. were found in aquatic habitats close to houses. In view of this, it is suggested that the availability of blood meals influences adult population dynamics, in turn, influencing the outcomes of interspecies competition in larval habitats.

Deliberate interference with mosquito access to human blood sources through LLIN and IRS interventions previously caused a species shift in the *An. gambiae* complex, so that the nominate species was competitively reduced [8] and eliminated across much of its range [4], notably across large tracts of east Africa, by *An. arabiensis* [4, 12, 15–17, 86, 107–109]. Such distinct competitive reductions in response to a vector intervention has also been previously documented for other malaria vectors [8, 12, 14, 84, 85, 87]. In Kenya, *An. funestus* were historically replaced by zoophagic and exophilic species from the same species group, namely *An. rivulorum* and *An. parensis*, following IRS that presumably affected the competitive balance in larval habitats [84, 85, 87, 110]. Furthermore, one of the most important vectors of malaria in South and Central America, namely *An. darlingi*, was previously eliminated in Guyana (formerly British Guiana), in response to an IRS event that subsequently created a species shift towards the more zoophilic vector, *An. aquasalis* [111].

These classic examples of shifting species composition in response to vector control interventions were therefore directly facilitated by selectively protecting humans from attack. Once this preferred host of the originally dominant mosquito species was no longer readily accessible, more zoophilic species were released from competition pressures [112], thus increasing in abundance. Here, we document similar species composition shifts across a spatial rather than a temporal scale, clearly underpinned by geographic variations in the natural availability and distribution of different blood resources, two of which (humans and cattle) are closely associated with anthropogenic insecticide coverage.

### Mammalian blood host availability and inferred host preferences of mosquitoes

Blood source utilisation patterns of *An. arabiensis* have been repeatedly characterized in diverse settings all across Africa [10] but the blood feeding habits of *An. quadriannulatus* are far less well understood. The clear association of *An. arabiensis* with detections of cattle or human blood sources reported herein is consistent with the known host preference behaviours of this malaria vector [24–26, 28]. The remarkably high intercept and strong effect size for detected activities of humans and cattle displayed in Table 2, clearly captures the specialised feeding behaviours demonstrated by *An. arabiensis,* for which no evidence of any other significant host could be detected with process explicit models fitted to data from pastoralist communities in northern Tanzania [28]. The consistency of these quantitatively dominant effects of humans and cattle (Table 2) with the known host preferences of *An. arabiensis* is reassuring with respect to the reliability of this methodological approach used.

As *An. quadriannulatus* are generally thought to be of little medical importance [24,25,54] their blood feeding behaviours have rarely been studied and remain poorly understood. Where this vector occurs in domesticated areas, it predominantly feeds on cattle [24] but will also feed on humans in some cases [113–115]. According to the classical literature, it is thought that *An. quadriannulatus* primarily survive on wild animals [24,24]. However, conclusive bloodmeal analysis for this species is almost lacking entirely, presumably because they are of such little medical interest and because it is notoriously difficult to collect resting blood fed specimens of such exophagic mosquito species [77, 116–118] A study by Prior and Torr (2002) [36] inside Mana Pools National Park, Zimbabwe, identified mixed blood-meals in a few specimens that included DNA from cattle and other unidentified hosts that were presumed to be wild ungulates. However, it seems that evidence for a definite link between *An. quadriannulatus* and a specific wild host species has not yet been reported.

While the *a priori* intention of fitting the host association model detailed in table 2 was to investigate the blood sources of the vector *An. arabiensis*, it also provided insight into the feeding behaviours of this highly zoophagic and seemingly non-vector in some of the most remote and best protected parts of the largest national park in Africa. Impala were identified as the most plausible and common blood source available to these *An. quadriannulatus* populations, an adaptation that may be attributed to several factors associated with the ecology of this antelope. Impala were the most frequently detected antelope in parts of NNP where high populations of *An. quadriannulatus* persisted and breeding herds of impala can comprise as many as 120 individuals [119]. This presents a large quantity of mammalian biomass, which is an important factor determining the host choice of blood-seeking adult mosquitoes [120–124].

Furthermore, impala have a flexible foraging ecology that encourages small home ranges and a sedentary existence. Impala are predominantly grazers in the wet season when grasses are plentiful but then readily switch to browsing in the dry season, allowing them to access reliable food supplies all year round This allows impala to form large resident herds that remain within much smaller home ranges (often less than 1km^2^) than larger more specialised antelopes, regardless of season [119]. Furthermore, these compact home ranges are specifically established close to perennial surface water sources, so they can drink daily and browse on the trees, shrubs and bushes that grow around such waterholes and streambeds throughout the dry season [119]. Impala are also least active after dusk and spend most of this time lying down [119], which coincides with the peak crepuscular biting activity peaks that are generally associated with zoophagy in mosquitoes [10,14,30]. Impala may therefore represent an especially reliable, abundant and historically widespread blood resource that is often concentrated around waterbodies where emergent blood-seeking mosquitoes may breed, and therefore an astute choice for *An. quadriannulatus* to have evolved to specialise upon.

Host-specialised blood utilisation behaviours are very common among anopheline mosquitoes, occurring in 82% of all studied species [124,125]. Such specialisation to feed on only one or two vertebrate species is thought to have evolved under the selection of several factors, notably the rates at which various potential host species are encountered [125]. For example, it has been shown that the speciation of the highly human-specialised *An. gambiae s.s.* arose with the arrival of Bantu agriculturists in the forest belt of central Africa [32].

Similarly, the unusually high historical abundance of impala across all the mixed acacia savannah habitat of eastern and southern Africa [119], may well have underpinned the evolution of *An. quadriannulatus* to specialise on this antelope that remain widespread across protected areas today.

Having said all that, it is also important to consider the limitations of associative observational studies when inferring preferred blood resources. For example, because other mammalian species were not detected as frequently inside NNP, the models detailed in Table 2 had less statistical power for evaluating the influence of these other potential blood host species. Correspondingly, a lack of other attributable influences may not necessarily imply that these numerous alternative mammals are not utilised as blood sources just as readily. It may well be the case that *An. quadriannulatus* simply feed flexibly and opportunistically upon whatever the most readily available hosts happen to be in any given locale, with impala being by far the most abundant in this case.

That being said, the statistical identification of bushpig as a second potential preferred blood host provides encouraging evidence of the validity and power of the approach taken, because even the small number of detections of this species, when compared to many other wild animals, proved sufficient to give plausible evidence that *An. quadriannulatus* may well utilise them as blood sources. Like impala, bushpigs are also social animals, living in groups of up to 15 individuals [119]. Their preferred habitats include woodland and forest that provide dense vegetation [119], consistent with their observed distribution across this study area, with smaller pockets of *An. quadriannulatus* found around such locations within ILUMA WMA. Bushpig forage at night, and often thrive in swampy areas at the interface between humans and wild animals [126]. Like impala, they are also highly water dependent with a sedentary existence, and permanently stay within remarkably small home ranges where they enjoy ready access to surface water all year round [119], and may therefore also offer quite reliable feeding opportunities for blood-seeking mosquitoes.

Of course, it is reasonable to question the reliability with which the activities of humans, livestock and such an impressive diversity of wild mammals were surveyed, especially given the dense vegetation cover across much of the study area and the cryptic or fluctuating habits of many of these mammalian species. However, practical experience during this study and detailed analysis of the animal activity survey data [56,57] both confirm that indirect observations of tracks, spoor and other signs were quite sensitive and reliable indicators, revealing the presence of almost every species envisaged at the outset, including 15 that were never directly sighted over the course of the two years. Although the activities of humans, livestock and wild animals were detected in several different ways, footprints were by far the most frequent indicator found around waterbodies. Indeed, the moist soil around the perimeter of many water bodies, particularly during the wet season, facilitated clear track indentations that could be confidently identified by the investigators and VGS carrying out the radial activity surveys. Although the age of the tracks was recorded during the survey, these distinctions were not considered for the analysis because the most recent prints were always more likely to be recorded, as they usually covered the tracks of older prints in such wet areas around waterbodies.

While the survey methodology proved useful for inferring host species, the greatest overall limitation of this study is simply that it was purely observational in nature and can therefore only demonstrate plausible indirect associations rather than probable direct causality. Unfortunately, although over 4,000 live adult mosquitoes were caught over the two years of this study, only a handful of these were blood-fed [127], despite the use of an interception screen specifically chosen to capture at least some blood-fed mosquitoes [117]. Consequently, such direct and conclusive evidence of feeding upon particular host species remains elusive.

Nevertheless, this novel methodology for associating the species composition in larval surveys with such indirect indicators of host availability has proven a simple, yet effective and surprisingly powerful approach that yielded clear evidence for four different preferred hosts of the *An. gambiae* complex within study area, two for *An. arabiensis* and two for *An. quadriannulatus.* However, beyond this malaria-related example, this new approach to studying host preferences of adult mosquitoes, based on their larval populations and the various mammals that frequent them, may have much broader applications for studying a wide range of vector borne zoonoses and emerging pathogens. Specifically, it may be deployed to fill a strategic methodological gap by allowing exploration of the adult feeding habits of zoophagic species, the vast majority of which are very poorly understood because they predominantly feed on wild hosts in expansive outdoor habitats and are, therefore, very difficult to capture or observe [116,118, 128]. Fortunately, larvae trapped within the water bodies they grow in are much easier to collect and identify than free-flying adults, so the new survey tactics describe herein could be applied to explore the full diversity of mosquitoes found in such aquatic habitats and the broad spectrum of blood sources they are thought to specialize upon to establish their distinctive ecological niches [125,129], even for cryptic wild vertebrate hosts that are difficult to survey by direct observation. By mapping out overlaps between mosquito species in terms of their preferred hosts in this way, it may well be possible to indirectly identify bridge vector species that could transfer pathogens from various wildlife species into livestock and/or humans.

Having said all that, the existence of *An. arabiensis* deep inside NNP could not be explained by any of the variables retained in the final multivariate model in table 2, so it was not possible to address the original objective of the study, which was to associate wild populations of *An. arabiensis* with one or more hosts other than humans and cattle. Prior to data collection, it was hypothesised that perhaps African buffalo might provide a plausible blood source for wild type mosquitoes because of their close phylogenetic relationship with cattle [130], combined with their size and propensity to often form large herds [119] that offer considerable biomass for host-seeking mosquitoes to exploit. This common wild bovine species have previously been suggested as a potential blood source for a wild population of *An. arabiensis* identified in Kruger National Park, South Africa, based on the observation that they frequented aquatic habitats that were occupied by larvae [37]. Furthermore, bloodmeal analysis from a study in Uganda identified African buffalo DNA in a blood-fed *An. gambiae* complex mosquito but unfortunately the specimen was not identified to species level [131]. As the host association model in Table 2 did not provide any evidence that the abundance of African buffalo or any wild herbivore favoured *An. arabiensis*, questions remain regarding the ecological niche of this vector species outside of landscapes that are at least partially domesticated, with humans and/or cattle available to support this mosquito species.

### Plausibility of self-sustaining An. arabiensis refuge populations in remote, fully intact wilderness areas

Given that *An. arabiensis* appears so well adapted to dry environments [3, 5–7, 33, 34], their presence deep inside NNP might possibly be explained by dispersal from surrounding domesticated lands. While, host-seeking adults from the *Anopheles gambiae* complex usually exhibit modest flight range of a few hundred metres to a few kilometres [25, 132, 133], they also sometimes disperse much further by exploiting high altitude winds [134–136]. Indeed, as no wild alternative hosts for *An. arabiensis* could be identified for this vector species inside fully intact, well protected wilderness areas (Table 2), it could be argued that these are *sink* populations [137,138] that may not be independently viable without sustained influx of already blood-fed adults from surrounding *source* populations [137,138] in domesticated villages. However, it is difficult to envisage how such long-range dispersal alone could sustain such readily detected densities of *An. arabiensis* so deep inside NNP for several reasons.

Firstly, the reciprocal of this spatial distribution pattern for *An. arabiensis* was not observed: Despite the presence of abundant *An. quadriannulatus* inside NNP, no specimens of this species were found in any of the nearby villages or even across most of the ILUMA conservation area. Notably, almost all (9/11) sampled location within the buffer zone represented by the WMA where *An. quadriannulatus* larvae were found also had detectable numbers of impala, bushpig or both. It therefore seems likely that local population viability, for *An. quadriannulatus* at least, is determined primarily by local conditions, particularly the availability of preferred hosts, at quite fine spatial scales.

Secondly, each individual *An. arabiensis* larva collected at *Shughuli Kubwa* (camp number 40), located 47km inside NNP from the park boundary (Figure 1) was found in a separate aquatic habitat, strongly suggesting that they originated from more than one mother. Also, several unfed, apparently host-seeking adult female *An. arabiensis* were also captured at the same remote location [127]. Furthermore, since the study was completed, another 52 *An. arabiensis* larvae were collected from a new location called *Mseguni* inside NNP (Figure 1), which is located even further away from humans and cattle than *Shughuli Kubwa*. This strongly suggests the existence of a well-established population that acquire bloodmeals locally, rather than an otherwise non-viable *sink* population sustained by external *source* populations [137,138] outside the park.

### Implications of wild An. arabiensis refuge populations living inside conservation areas

It was clear that unauthorised human and livestock encroachment was extensive throughout most of the ILUMA WMA and had a substantial negative impact on the availability and distribution of wild animals [56,58]. In particular, illegal cattle herding represents an influx of preferred blood sources for *An. arabiensis* and all major forms of encroachment reduce the availability of wild mammalian blood sources, thus displacing the non-vector *An. quadriannulatus* in favour of the malaria vector it competes with. Through improved management and conservation efforts of community-based conservation areas, to effectively control unauthorised, illegal human activities and facilitate the return of wild animals to the WMA, a natural reduction of *An. arabiensis* in favour of *An. quadriannulatus* may well follow. Such competitive displacement by closely-related species may well help explain the remarkably swift and decisive elimination of highly human-specialized malaria vectors by LLINs or IRS on several occasions, in a rapid and decisive manner that cannot be explained with simple single-population models [8,12]. Therefore, natural competitive suppression of this important malaria vector might represent a previously undocumented form of natural capital that arises from supporting and protecting conservation areas.

However, while it may be possible for *An. quadriannulatus* to competitively suppress *An. arabiensis* in conservation areas to some extent, the latter vector species was clearly not completely displaced from fully intact natural habitats lacking cattle or humans, even in the wildest areas with plenty of the wild herbivores the former species appears to prefer. This highlights how the behavioural plasticity of *An. arabiensis* allows it to extend its ecological niche deep inside wild areas, where they can even compete for resources against this much more zoophagic member of the same complex. Furthermore, the apparent existence of such refugia populations of *An. arabiensis* in these remote, well protected ecosystems confirms the initial hypothesis that alternative blood resources create an additional population-stabilising portfolio effect [40] for this species, further emphasizing the major vector control challenges posed by its notoriously flexible feeding behaviours [10, 14, 20, 31].

Given that *An. arabiensis* were clearly and robustly the dominant competitive species wherever any humans or cattle whatsoever were found across the study area, and a refuge population of this species even exists deep inside NNP where there are none, the prospects for eliminating this key vector of residual malaria transmission with conventional human- centred vector control measures appear remote. Not only are wild refugia populations feeding upon wild animals largely out of the reach of current first-choice LLIN and IRS interventions that protect humans where and when they sleep, usually in reasonably well-established houses and shelters, they may also limit the population suppression effects [10] of new complementary approaches like spatial repellents and veterinary endectocides [139] that respectively extend insecticidal coverage to humans and livestock outdoors [11]. Furthermore, in parts of neighbouring Zambia where sympatric populations of *An. quadriannulatus* and *An. arabiensis* occur in domesticated settings, it seems that the former is much more vulnerable to insecticide attack than the latter, as evidenced by an abrupt species composition shift towards the latter vector species immediately after effective scale up of IRS [140]. It might therefore be difficult to target adult *An. arabiensis* with insecticide-based interventions without also affecting *An. quadriannulatus* populations at least as much.

On the other hand, two notable historical examples of successful elimination of this species from Brazil [141–143] (Retrospectively confirmed as *An. arabiensis* by molecular techniques [144]), and Egypt [145] (By far the most likely culprit based on ecological niche mapping [4,7]), primarily through comprehensive larvicide application, suggest it can nevertheless be tackled decisively at source as larvae, before they can express the evasive behaviours that make the adults so difficult to control. In both cases, however, *An. arabiensis* was an invasive species that had established itself within a limited, albeit quite extensive new geographic range, within which it proved possible to contain, suppress and eventually eliminate. It is very difficult to envisage such “scorched earth” larval source management campaigns being feasible or affordable across the vast natural range of this species in Africa, especially if that includes refuge populations in the many large tracts of largely uninhabited wilderness scattered across the continent.

On a more positive note, although wild refugia populations deep inside NNP pose clear challenges for vector elimination, they also provide a potential opportunity for reversing the deleterious effects of pyrethroid resistance on the progress of malaria control to date [41–49]. Self-sustaining populations that do not come into contact with humans, livestock or agriculture, and are therefore free from the selective pressures of pesticides, may represent invaluable reservoirs of relatively unmodified mosquito genomes, complete with original wild-type insecticide susceptibility traits that would otherwise have been lost from the natural gene pool. The existence of such large reservoirs of wild-type genomes that have never been bottlenecked by insecticide pressure, could enable insecticide resistance management strategies that exploit astute insecticide combinations to select back these traits [146] in neighbouring populations living alongside people, thereby restoring the effectiveness of current [147] and future [148–152] interventions that remain dependent on this exceptionally useful insecticide class.

## Conclusions

While only *An. arabiensis* was found in fully domesticated ecosystems, its non-vector sibling species *An. quadriannulatus* occurred in conserved areas and dominated the best conserved natural ecosystems. As its two known preferred hosts [27,28] the importance of humans and cattle to *An. arabiensis*, was confirmed by its clear positive association with signs of activity by these two mammalian host species. The relative abundance of *An. quadriannulatus* increased with distance inside NNP and away from human settlements and was positively associated with activities of impala and bushpig, specifically, implicating both as likely preferred blood hosts. While abundant impala and lack of humans or cattle in fully intact acacia savannah within NNP apparently allowed it to dominate *An. arabiensis*, presence of bushpig seemed to provide it with a foothold in miombo woodlands of the ILUMA WMA, despite considerable encroachment by people and livestock. While this antelope and suid are essentially unrelated, both are non-migratory residents of small home ranges with perennial surface water, representing likely preferred hosts for *An. quadriannulatus* that are widespread across extensive natural ecosystems all year round. Despite dominance of *An. quadriannulatus* in well-conserved areas, *An. arabiensis* was even found in absolutely intact natural environments >40km inside NNP, suggesting it can survive on blood from one or more unidentified wild species. Self-sustaining refuge populations of *An. arabiensis* inside conservation areas, supported by wild blood hosts that are fundamentally beyond the reach of insecticidal interventions targeted at humans or their livestock, may confound efforts to eliminate this key malaria vector. This new approach to indirectly identifying the most commonly utilized blood sources may well be applicable to an unprecedented diversity of zoophagic mosquitoes in particular, enabling the incrimination of possible bridge vector species potentially capable of mediating pathogen spillover from wildlife reservoirs into livestock and/or human populations.

## Methods

As explained in S5 Appendix, this study was conducted in the Kilombero valley in the Morogoro region of southern Tanzania, where the local malaria transmission systems and vector populations have been exceptionally well-characterised.

At the outset, the study design was centred around the ILUMA WMA but was then later extended deep into NNP (S5 Appendix). This large study area was fundamental for addressing the objectives of this specific study and the broader goals of the overall project, as it represents a geographical gradient of natural ecosystem integrity ranging from fully domesticated land uses in the west to completely intact, well conserved natural ecosystems to the east.

This comprehensive range of de facto land use practices and ecosystem characteristics were also considered ideal for investigating the influence of a wide range of the potential blood host species upon the population dynamics of *An. arabiensis*, potentially leading to heterogeneities in host availability that could create portfolio effects that buffer populations of this important vector against control measures [40].

This study was carried out using a rolling cross-sectional design with four rounds of surveys encompassing a total of 40 defined locations, referred to herein as *camps,* over the course of two years (Figure 1 and S5 Appendix), which were carefully selected to encompass a range of land uses and ecosystem integrity states with a wide range of mammalian species abundance and diversity (S5 Appendix and S8 Appendix). Under the original protocol, a total of 28 camp locations were identified in or immediately to the west of the ILUMA, most of which were sampled over 3 distinct survey rounds completed in 2022. However, no camp within ILUMA lacked any signs of human disturbance and only a few remained relatively intact and undisturbed (Table in S5 Appendix). Furthermore, none of the F2 adults initially reared from wild F_0_ specimens collected within the WMA exhibited pyrethroid susceptibility phenotypes, so 12 new camps inside NNP were added for a fourth and final round of surveys between February and July 2023, spanning the whole wet season and the beginning of the dry season for that calendar year.

Quantitative larval surveys were carried out within a 2km radius of each camp (S3 Appendix) and complemented by a survey for any signs of activity by wild or domesticated mammals (Figure 10) and humans, as well as land use and vegetation cover, around each habitat and *en route* between them (S8 Appendix [56,57])

**Figure 10.**
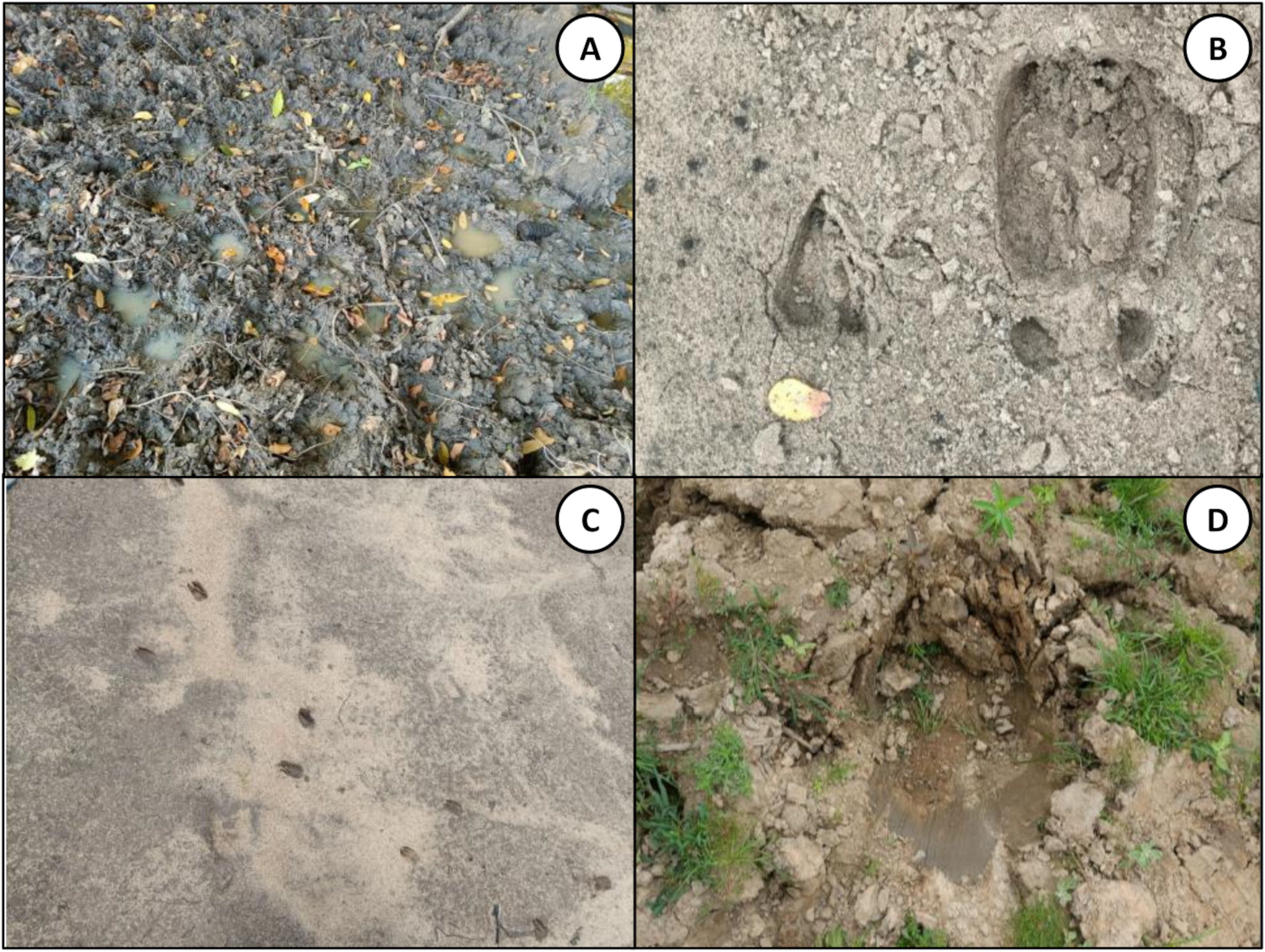
Examples of animal prints that were recorded as detections during the radial surveys of human, livestock, wildlife activity and land use. A: collection of cattle hoofprints surrounding a pool, B; hartebeest and a buffalo print in sand, C; common duiker prints in sand, D; a hippo print on the riverbank.

As the only PCR identified larval samples that were collected and preserved *in situ* under the revised protocol (S3 Appendix) during the fourth round were used for the final analyses of sibling species composition of the *An. gambiae* complex, only the fourth round of complimentary radial surveys, completed at exactly the same times and places, were used to test for associations between the species composition and the availability of various potential hosts. Hence, any detection from the radial activity surveys that complemented the larval survey data from rounds one to three were omitted from the analyses for that particular purpose. Therefore, any larvae from an aquatic habitat could be spatially and temporally associated with the frequency and types of human, livestock or wildlife detections that were recorded around the aquatic habitats that the larvae were collected from at the time they were collected. Recorded detections from every survey over all four rounds were, however, used to calculate a formal and objective *natural ecosystem integrity index (ONEII)* that was used to evaluate a subjective *natural ecosystem integrity index* (SNEII), a powerful and less laborious indicator of ecosystem integrity based on consensus perceptions of the investigators (S6 Appendix (56, 58)). All statistical analyses of An. gambiae complex species occupancy and sibling species composition within larval habitats and their associations with distance from the nearest NNP boundary or human settlement, ecosystem integrity, historical landcover and activities of humans, livestock and wildlife were conducted using generalized linear mixed models as detailed in S10 Appendix.

## Acknowledgements

The authors wish to thank the Village Game Scouts of ILUMA WMA, for their hard work and participation in field activities. We also thank all the governance, management, and stakeholder communities of the ILUMA WMA for all their collaboration and kind assistance. Furthermore, we thank Mr Frederic Masanja and Mr Fadhili Songo for all the essential institutional support provided by the Ifakara Health Institute over the course of the study. We also extend our appreciation to Ms. Elaine Kelly, Ms. Christine Dennehy, Ms Marie Riordan, Dr. Ronan Hennessy and Mr. Allen Whittaker at UCC, for their help with many administrative and technical issues. We also thank Dr. Ramiro Crego and Dr. Tom Reed at the University College Cork, Prof. Felister Mombo at the Sokoine University of Agriculture, Mr. Japhet Kihonda at the Ifakara Health Institute, Dr. Diego Ayala at the Institute of Research for Development, and Dr. William Hawley at the Centres of Disease Control and Prevention, for their scientific advice and input. A very special word of thanks is due to our recently deceased friend and colleague, Mr Octavian Malopola, without whom this work would never have even begun.

## Availability of data and materials

Mosquito larvae occupancy and habitat attribute data collected from the field is available as S11 Appendix. The humans, livestock and wildlife data collected from the field and mosquito larvae species composition data from PCR are available as S12 Appendix.

### Supporting Files

S1 Appendix. Habitat characteristics and occupancy by mosquito larvae, particularly those from the *Anopheles gambiae* complex

S2 Figure. Preliminary *An. gambiae* complex species composition results from F0 adults raised from field collected larvae.

S3 Appendix. Detailed larval survey protocol.

S4 Appendix. Validation of field identification methodology for larvae from the Anopheles gambiae complex

S5 Appendix. Study Area and Sampling Frame.

S6 Appendix. Assessment of natural ecosystem integrity, historical natural land cover and distance from nearest national park boundary or human settlement.

S7 Figure. The detection frequency of humans plotted against the detection frequency of cattle herds at each camp number demonstrating a strong linear correlation, as tested using a Pearson’s linear correlation test.

S8 Appendix. Procedures for surveying activities of humans, livestock and wildlife, as well as land use and vegetation cover, around mosquito larval habitats within a 2km radius of each camp location.

S9 Appendix. Conceptual framework for regression analysis of the association between Anopheles gambiae sibling species composition and indicators of activity by diverse mammalian species that could act as potential sources of blood for mosquitoes.

S10 Data entry, cleaning, preparation and analysis

S11 Data. All the data from the larval habitat occupancy and habitat attribute surveys.

S12 Data. The species composition data from the larvae that were collected in the field, and all the data from the complimentary humans, livestock and wildlife surveys.

