## Supplementary material for "Blood host preferences and competitive inter-species dynamics within an African malaria vector species complex inferred from signs of animal activity around aquatic larval habitats": S1_Appendix

**S1 Appendix: Habitat characteristics and occupancy by mosquito larvae, particularly those from the *Anopheles gambiae* complex**

**Results**

A total of 8,266 dips across 1,944 potential larval habitats were completed over the sampling period from January 2022 to July 2023. The relative frequencies of each attribute for the total number of habitats are summarised in Table S1.1. Larvae from the genus *Anopheles* were present in a total of 1,058 habitats. Out of these positive habitats for *Anopheles* larvae, 72.3% were also identified as having *An. gambiae* complex present, demonstrating an overall occupancy rate of 39.4% for this species complex (Table S1.1).

*Culex* larvae were present in 763 habitats indicating lower overall occupancy (39.2%) than *An.* larvae (Table S1.1). The combined presence of both early and late instar larvae was the most frequent observation for both *Anopheles* and *Culex* mosquitoes (32.3%, 23.6%), followed by late stage only (15.1%, 10.0%), and early stage only (6.9%, 5.6%), respectively (Table S1.1). Pupal stages were identified in only 122 habitats (6.3%).

**Table S1.1:** Frequencies and proportions of the recorded biological, physical, spatial, and temporal attributes of all the habitats surveyed. Total habitat observations are equal to the sum of the number of habitats that possessed the given attribute at the time it was surveyed, totalled across all four rounds from January 2022 to July 2023.

| Attribute | Total Habitat Observations | Proportion of Total Habitats (%) |
|---|---|---|
| *Anopheles larvae* | | |
| Absent | 886 | 45.6 |
| Present | 1058 | 54.4 |
| Early instar only | 134 | 6.9 |
| Late instar only | 293 | 15.1 |
| Early and late instar | 631 | 32.3 |
| *An. gambiae* complex present | 765 | 39.4 |
| *Culex larvae* | | |
| Absent | 1181 | 60.8 |
| Present | 763 | 39.2 |
| Early instar | 110 | 5.6 |
| Late instar | 194 | 10.0 |
| Early and late instar | 459 | 23.6 |
| *Pupal stage* | | |
| Absent | 1822 | 93.7 |
| Present | 122 | 6.3 |
| *Habitat Type and Description* | | |
| Pools, puddles tracks & depressions | 1343 | 69.1 |
| Springs and swampy areas | 125 | 6.4 |
| Flowing streams | 97 | 5.0 |
| Flooded valleys | 7 | 0.4 |
| Waterholes | 173 | 8.9 |
| Artificial drains or ditches | 4 | 0.2 |

| | | |
|---|---|---|
| Human dug wells | 105 | 5.4 |
| Artificial containers | 2 | 0.1 |
| Rice paddies | 69 | 3.5 |
| Ridge and furrow agriculture | 11 | 0.6 |
| Other agriculture | 5 | 0.3 |
| Other | 3 | 0.1 |
| *Habitat Perimeter* | | |
| <2m | 270 | 13.9 |
| 2m-20m | 1092 | 56.2 |
| 21-200m | 402 | 20.7 |
| >200m | 180 | 9.2 |
| *Number of Dips* | | |
| 1 | 174 | 8.9 |
| 2 | 182 | 9.4 |
| 3 | 395 | 20.3 |
| 4 | 337 | 17.3 |
| 5 | 585 | 30.1 |
| 6 | 50 | 2.6 |
| 7 | 52 | 2.7 |
| 8 | 34 | 1.8 |
| 9 | 16 | 0.8 |
| 10 | 119 | 6.1 |
| *Water Depth* | | |
| <0.1 | 1256 | 64.6 |
| 0.1 – 0.5 | 650 | 33.4 |
| >0.5 | 38 | 2.0 |
| *Short Vegetation* | | |
| Absent | 1301 | 66.9 |
| Present | 643 | 33.1 |
| *Tall Vegetation* | | |
| Absent | 1580 | 81.3 |
| Present | 356 | 18.7 |
| *Floating Vegetation* | | |
| Absent | 1780 | 91.6 |
| Present | 164 | 8.4 |
| *Location* | | |
| Inside ILUMA WMA | 1072 | 55.2 |
| Inside NNP | 294 | 15.1 |
| Outside ILUMA WMA | 577 | 29.7 |
| *Historical Landcover* | | |
| Miombo woodland | 1671 | 86.0 |

| | | |
|---|---|---|
| Groundwater forest | 123 | 6.3 |
| Acacia savanna | 150 | 7.7 |
| *Season* | | |
| Wet | 1365 | 70.2 |
| Dry | 579 | 29.8 |
| *Round* | | |
| 1 | 234 | 12.0 |
| 2 | 591 | 30.4 |
| 3 | 559 | 28.8 |
| 4 | 560 | 28.8 |

Pools, puddles, tracks, and depressions were the most frequent habitat type accounting for more than two thirds of all habitats surveyed, followed distantly by waterholes, and springs and swampy areas, which only accounted for 6% of all habitats (Table S1.1). Human-dug wells and rice paddies were the most frequently surveyed man-made habitats, representing approximately 5% and 4% of all habitats surveyed, respectively (Table S1.1). Other artificial or man-made habitats combined only accounted for just over 1% of all surveyed habitats (Table S1.1). The frequency distribution of habitat type is also reflected by distribution patterns of the habitat's other physical attributes. More than half of all habitats had a perimeter between 2m and 20m (Table S1.1), which naturally included many from the pools, puddles, tracks, and depressions category. Whereas aquatic habitats exceeding 20m in perimeter, accounted for less than one third of all those surveyed (Table S1.1), and would have been mostly associated with less frequently sampled waterbodies such as rice paddies, springs and swampy areas, waterholes, and flooded valleys (Table S1.1). As the number of dips taken was dependent on habitat size (S3 Appendix), and more than 85% of the habitats were dipped five times or less, all habitats with a perimeter less than 20m fitted into this modest size category, as did many that were greater than 20m but were not fully accessible. Almost two-thirds of habitats were shallower than 10cm (Table S1.1) and were absent in most habitats especially the pools, puddles, tracks, and depressions category.

About 70% of habitats were surveyed during the wet season (Table S1.1) when surface water and mosquitoes were most abundant. The first round of the study had the lowest proportion of habitats sampled (Table S1.1) because one circuit was omitted, so only 20 out of a possible 28 camps were visited. Furthermore, data from the first six camps surveyed were omitted for analysis due to missing data in the *number of dips* column, which was only introduced to the data collection form (S3 Appendix, table S3.5) two weeks into the study. Although the number of camps slightly differed per round, the number of habitats surveyed during rounds 2, 3 and 4 remained roughly consistent (Table S1.1). The habitats surveyed in round two were conducted in all the original 28 camps located inside ILUMA WMA and outside it in the villages to the west (S5 Appendix, table S5.3). By the time the third round was conducted in August 2022, water sources in four camps in the north-east circuit had fully dried out and therefore were not surveyed. Round three also included four new camps inside NNP, collectively known as the NNP-Boma Ulanga circuit (S5 Appendix, table S5.3). For round 4, potential larval habitats were surveyed across a total of 40 camps, including the initial 28 camps, the additional four camps that were added at the end of round 3, and a further eight camps located deeper into NNP that were visited during two separate circuits, namely the NNP-Kilombero River and NNP-Msolwa circuits (S5 Appendix). The number of habitats surveyed inside the national park accounted for only 15% of the total over the 4 rounds (Table S1.1), as they were added at the last round in the final stage of the study. A total of 22 out of 40 camps were located inside ILUMA WMA and all of them were surveyed at least twice (S5 Appendix, table S5.3), so they correspondingly accounted for more than half of all aquatic habitats inside miombo woodland. Only 6% of habitats were surveyed in the groundwater forest (Table S1.1), because this landcover type was only found in the north of ILUMA WMA, where the availability of surface water was mainly limited to waterholes and small puddles. Dry acacia savanna landcover was exclusively found in a small number of camps well inside NNP that were surveyed only once or twice (S5 Appendix, table S5.3).

A forward-step selection process was used to build the multivariate generalized linear mixed model (GLMM) described in Table S1.2, allowing any confounding effects of habitat attributes on occupancy by *An. gambiae* complex larvae to be accounted for. The effect of season on occupancy was no longer significant once round and habitat type were accounted for, probably because each round was designed to represent a period in the wet or dry season and therefore captures most of the variance with season. The covariate habitat type may also have had confounding effects as occupancy was higher in waterholes and rice paddies, which were mostly seasonal, and were therefore more common after rainfall during the wet season. Floating vegetation and tall vegetation also proved to have no effect on the occupancy of *An. gambiae* complex once other covariates were accounted for. Habitat type may account for any variance associated with these covariates, as these vegetation types were often closely associated with specific types of waterbodies. For example, tall vegetation would be characteristic of late-stage rice paddies and floating vegetation was commonly observed in waterholes.

**Table S1.2 Generalised additive mixed model univariate and multivariate outputs for the proportion of habitats occupied by *An. gambiae* complex larvae based on spatial, temporal, abiotic and biotic habitat attributes fitted to a binomial distribution and a logit link function.** Variation between camps across rounds was accounted for as a random effect. First order temporal autoregression was accounted for by nesting the number of weeks since the first survey in camp number. Statistically significant effects are highlighted in bold.

| | **Univariate** | | | **Multivariate** | | |
|---|---|---|---|---|---|---|
| Fixed Effects | OR [95% CI] | t | P | OR [95% CI] | t | P |
| *Intercept*[a] | NA | NA | NA | 1.35 [0.83, 2.21] | 1.21 | 0.2268 |
| *Survey Implementor* | | | | | | |
| Individual 1 | 1.00 | NA | NA | NE | NE | NE |
| Individual 2 | **3.91 [2.91, 5.24]** | **9.09** | **<<0.0001** | NE | NE | NE |
| **Temporal and spatial characteristics** | | | | | | |
| *Round number* | | | | | | |
| 1 | **0.36 [0.23, 0.56]** | **-4.50** | **<<0.0001** | **0.27 [0.17, 0.43]** | **-5.60** | **<<0.0001**[d] |
| 2 | **0.29 [0.21, 0.41]** | **-7.19** | **<<0.0001** | **0.21 [0.15, 0.31]** | **-8.41** | **<<0.0001**[d] |
| 3 | **0.19 [0.13, 0.27]** | **-9.20** | **<<0.0001** | **0.18 [0.12, 0.26]** | **-9.04** | **<<0.0001**[d] |
| 4 | 1.00 | NA | NA | 1.00 | NA | NA |
| *Season* | | | | | | |
| Wet | 1.00 | NA | NA | 1.00 | NA | NA |
| Dry | **0.41 [0.30-0.55]** | **-5.93** | **<<0.0001** | 0.37 [0.09-1.44] | -1.44 | 0.1513[e] |
| *Land cover* | | | | | | |
| Miombo woodland | 1.00 | NA | NA | 1.00 | NA | NA |
| Groundwater forest | 0.82 [0.43, 1.56] | -0.60 | 0.5467 | 0.76 [0.37, 1.60] | -0.71 | 0.4746[e] |
| Acacia savannah | **1.93 [1.08, 3.44]** | **2.23** | **0.0262** | 0.89 [0.45, 1.76] | -0.33 | 0.7400[e] |

| | | | | | | |
|---|---|---|---|---|---|---|
| *Distance* | 1.01[0.99, 1.03][b] | 1.59 | 0.1120 | 0.97 [0.96, 0.99][b] | -1.83 | 0.0678[e] |
| *Location* | | | | | | |
| Inside ILUMA | 1.00 | NA | NA | 1.00 | NA | NA |
| Outside ILUMA | 1.09 [0.68, 1.77] | 0.36 | 0.7165 | 0.60 [0.32, 1.13] | -1.59 | 0.1132[e] |
| Inside Nyerere NP | 1.34 [0.83, 2.18] | 1.20 | 0.2313 | 0.96 [0.54, 1.73] | -0.12 | 0.9042[e] |
| *SNEII* | 0.91 [0.91, 0.92][c] | -0.33 | 0.7410 | **0.38 [0.37, 0.38][c]** | **-3.80** | **<0.0001[d]** |
| **Abiotic habitat characteristics** | | | | | | |
| *Habitat type* | | | | | | |
| All other habitats | 1.00 | NA | NA | 1.00 | NA | NA |
| Waterholes & rice paddies | **2.58 [1.95, 3.42]** | **6.61** | **<<0.0001** | **1.78 [1.22, 2.58]** | **3.01** | **0.0028[d]** |
| Springs, streams & swamps | **1.72 [1.30, 2.26]** | **3.85** | **<0.0001** | 1.02 [0.71, 1.46] | 0.08 | 0.9332[e] |
| *Perimeter* | | | | | | |
| <2m | 1.10 [0.84, 1.45] | 0.69 | 0.4900 | **1.61 [1.15, 2.26]** | **2.74** | **0.0058[d]** |
| 2m-20m | 1.00 | NA | NA | 1.0 | NA | NA |
| >20m | **1.84 [1.51, 2.24]** | **4.80** | **<<0.0001** | 1.19 [0.90, 1.58] | 1.29 | 0.2194[e] |
| *Number of dips* | **1.14 [1.09, 1.18]** | **6.03** | **<<0.0001** | **1.15 [1.08, 1.22]** | **4.34** | **<<0.0001[d]** |
| *Water depth* | | | | | | |
| <0.1 | 1.00 | NA | NA | 1.00 | NA | NA |
| *0.1-0.5* | 0.98 [0.80, 1.18] | -0.25 | 0.8029 | 0.82 [0.65, 1.03] | -1.71 | 0.0871[e] |
| *>0.5* | 0.54 [0.28, 1.05] | -1.81 | 0.0711 | 0.59 [0.28,1.26] | -1.36 | 0.1732[e] |
| **Biotic habitat characteristics** | | | | | | |
| *Short vegetation* | | | | | | |
| Absent | 1.00 | NA | NA | 1.00 | NA | NA |
| Present | **1.68 [1.37, 2.05]** | **5.02** | **<<0.0001** | **1.30 [1.03, 1.63]** | **2.22** | **0.0261[d]** |
| *Floating vegetation* | | | | | | |
| Absent | 1.00 | NA | NA | 1.00 | NA | NA |
| Present | **1.62 [1.17, 2.225]** | **2.89** | **0.004** | 0.95 [0.64, 1.43] | -0.23 | 0.8187[e] |
| *Tall vegetation* | | | | | | |
| Absent | 1.00 | NA | NA | 1.00 | NA | NA |
| Present | 1.28 [1.01, 1.64] | 2.02 | 0.0433 | 0.76 [0.56, 1.02] | -1.86 | 0.0632[e] |

| Random Effects | σ |
|---|---|
| *Camp number* | 0.351 |
| First order continuous autoregression | Phi |
| *Weeks since start of study/Camp number* | 0.232 |

OR; Odds ratio.
95% CI; The 95% confidence interval
[a] Larval occupancy under reference conditions.
NA; Not applicable because several different intercepts were estimated for more than one fitted univariate model, or not applicable to the reference group.
NE; Not estimated due to confounding effects.

[b] Odds ratio for every kilometre increase from the NNP boundary.
[c] Odds ratio between fully domesticated and fully intact natural habitats
[d] As estimated from the final best fit GLMM where the effect was included based on a significant value <0.05.
[f] As estimated from the point at which the effect was no longer significant (>0.05) in the multivariate model fit and was therefore no excluded from the final model.
σ; standard deviation

Ecosystem integrity, survey round, habitat type, habitat perimeter, short vegetation and the number of dips were all significant predictors of *An. gambiae* complex larval occupancy in aquatic habitats. Ecosystem integrity was the third most important predictor of larval occupancy, indicating a significant decrease in occupancy rates between aquatic habitats in fully domesticated ecosystems and aquatic habitats in fully intact natural ecosystems (Table S1.2, Figure 2), suggesting that larval occupancy was somewhat lower for aquatic habitats in well conserved areas further away from human settlements and domesticated land use. Despite this, *An. gambiae* complex larvae, were nevertheless present with mean occupancy rates exceeding 25%, even in the best conserved environments far away from any signs of humans or livestock (Figure 2). This indicates that adult mosquitoes from the *An. gambiae* complex were ovipositing year-round in aquatic habitats even >40km away from the nearest resident human or livestock hosts.

The odds of occupancy by *An. gambiae* complex larvae were 73% (P <<0.0001), 79% (P <0.0001) and 82% (P <<0.0001) lower, respectively for each of the consecutive rounds completed in 2022 when compared to all the habitats surveyed in round 4, which was completed the following year in 2023. This significant difference may probably be attributable to the two individuals that implemented the surveys, with the new person who took over for round 4 exhibiting an exceptionally meticulous approach. As round and surveyors were obviously closely associated, the latter variable was dropped from the multivariate model because round captured all the variance attributed to the former plus any additional variance that occurred in between replicates.

Abiotic and biotic predictors of *An. gambiae* complex occupancy included habitat type, the number of dips taken, habitat perimeter and the presence of short vegetation. Although the springs, swamps and streams category, was no longer significant in the multivariate analysis, habitats that were classified as waterholes or rice paddies had higher occupancy rates compared to the reference category of other habitats. The apparent effect of habitat perimeter differed between univariate and multivariate analysis, probably because it was highly correlated with the number of dips made. The number of dips was the second most important predictor of habitat occupancy and the effect size remained consistence across univariate and multivariate analysis. In the final model fit, each additional dip increased the odds of a habitat being occupied with *An. gambiae* complex larvae by 15%. Therefore, the effect size estimated for larger habitats that had perimeters greater than 20m, was reduced and was no longer statistically significant, probably because it was confounded by the higher number of dips taken from them. The model also revealed that the odds of occupancy was higher by almost two thirds in the smallest habitats (<2m in perimeter), which were dipped or sampled with a turkey baster only once or twice, compared to those of intermediate size (perimeters between 2m and 20m). The effect of short vegetation was less pronounced in the final model than in the univariate analysis.

### Discussion

The remarkably large differences in reported occupancy rates between the investigator and the trained VGS member may have been influenced by several factors including subtle variations in sampling methodology and technique. Mosquito larvae are extremely sensitive to predation threats and so if disturbed, they will rapidly swim downwards from the water surface or sidewards along it. Simple technique adjustments, like avoiding casting shadows across the habitat before sampling it, and improving the angle and motion of dipper deployment (1, 2) may have increased the odds of *An. gambiae* complex larvae capture. More specifically, it was notable that the VGS member also often meticulously investigated the habitat perimeter prior to dipping, thus increasing detection sensitivity by dipping at points where larvae were already observed. While this could of course introduce biases of its own, including the additional year of surveys completed by the VGS volunteer improved the statistical power of the model fit, indicating that the patterns exhibited were consistent across these two years with two different surveyors. It should be noted however, that such large differences between different individual surveyors are quite normal (3, 4) as increases in larval habitat detection among *community-owned resource persons* has previously been clearly documented in an operational municipal larval source management programme in Dar es Salaam, Tanzania. Interestingly, volunteers that were familiar with the area and were recruited through local community leaders were better at detecting larval habitats than those who were recruited by programme leaders and administrative staff  (4).

Pools, puddles, tracks, and depressions were the most frequently surveyed habitat category encompassing a wide range of different waterbodies with similar appearances. Although this broad habitat category was available year-round, it was observed that specific waterbodies that were available within this category were influenced both seasonally and geographically. For example, during the wet season, this category included any surface-water pools of accumulated rainwater and any human, animal, vehicle tracks or other indentations formed in sodden soil. During the dry season, surface-water under this category was limited to stagnant isolated pools formed by retreating streambeds, which were observed to be the predominant habitat type during the driest months when waterholes, swamps, flooded valleys, and rice paddies dried up.

The habitat attributes found to favour *An. gambiae* complex larvae in this study were consistent with those reported by classic literature, which includes small, open sunlit habitats (5, 6). Although habitat occupancy was higher in habitats with vegetation, it was observed to be often sparsely distributed. Rice paddies positive for *An. gambiae* complex in this study were typically in the early stages of cultivation when the crop height was low, an observation that was also consistent with those reported in Gillies and De Mellion, 1968). Furthermore, as per the protocol and to avoid trampling the crop, dips were taken along the margins of the field that would have been more exposed to the sun and provided optimal conditions for developing larvae (5, 6).

The results showed that rice paddies, waterholes and small habitats (<2m), were all characteristic attributes of higher *An. gambiae* complex occupancy rates compared to any other habitat type (Table S1.2). As there was no difference in occupancy rates between fully domesticated settlements outside ILUMA WMA and inside NNP (Table S1.2) and no rice paddies exist in NNP because the area is under strict conservation, this indicates that waterholes and the surrounding prints of wild animals may well be predominant breeding sites for wild populations of *An. gambiae* complex mosquitoes. Therefore, to conduct further research on the aquatic stages or, when attempting to catch adult mosquitoes in wild areas far from human activity, it is recommended to set up camp near waterholes rather than alongside other habitats such as flowing streambeds, which proved to be occupied less (Table S1.2). However, although not detected in the model, it was also observed by both investigators that when water levels were low during the dry season, small pools found on the surfaces of exposed rocky outcrops in riverbeds were often occupied with *An. gambiae* larvae, particularly along the Kilombero River deep inside NNP.

The overall occupancy rate estimated from this study was 54% for all *Anopheles* larvae and 39% for *An. gambiae* complex larvae, which was reasonably consistent with some other reported occupancy rates, in rural towns or settlements over the last two decades, which ranged from about 35% to 70% (7-16). For example, Fillinger *et al.* (2004) identified 67% of surveyed aquatic habitats as being occupied with *An. gambiae* complex larvae, while a second study in Kenya also demonstrated a similar occupancy rate of 51% for all *Anopheles* larvae (10). More recently, a study by Epopa *et al.* (2020) (16) demonstrated anopheline occupancy rates of 46% which was conducted in an area of low human density between two villages. Indeed, because the presence of anopheline larvae in aquatic habitats is highly ubiquitous (5), and the larval ecology varies between anopheline species (6, 17), even within the *An. gambiae* complex (18-21), one can't totally compare or over interpret differences in the reported occupancy rates of anopheline larvae between these studies. Additionally, as larval surveys are ordinarily conducted in areas of human settlement, the distribution of sampled breeding sites is biased towards man-made habitats that represent productive breeding sites, so natural habitats far away from people may not be fully represented.

Although the overall occupancy rates were relatively consistent with past and present reports, this study demonstrated for the first time, a modest but steady decrease in occupancy rates over a gradient of ecosystem integrity (Figure 2) that extended almost 80km inside a newly designated and as of now, undeveloped national park with negligible human intrusion. The unanticipated extension of the sampling frame into NNP at a later stage of the study, meant that larval surveys at these camps were only replicated once or twice compared to the camp locations inside ILUMA WMA and in the villages along its western boundary, that were part of the original protocol and almost all surveyed three or four times (S5 Appendix, table S5.3). Correspondingly, less occupancy data was available for these additional camps that generally represented the furthest distances from human settlement and had the highest SNEII scores overall. However, the statistical power of the final fitted model in Table 1, as indicated by the low p-values and narrow confidence intervals, suggests that although the number of replicates was somewhat lower for the best conserved camps, the number of habitats surveyed at each camp were sufficient to give a robust model fit.

The SNEII is a variable that captures the overall general condition of the ecological state of an area and therefore reflects a myriad of other more specific factors that could act as hidden drivers of lower occupancy rates in well conserved locations. In particular, it is closely associated with the distance from human settlement, with the lowest SNEII scores assigned to those in fully domesticated settlements and higher SNEII scores generally assigned much further away from permanent human settlements, with the highest being in NNP. In other settings, aquatic habitats that are located closer to human settlements have been found to have higher occupancy and larval density than those located further away (16, 22, 23), indicating that proximity to known preferred blood sources influences the rates at which *An. gambiae* complex use suitable habitats. However, these studies were conducted across scales of hundreds of metres, while the study reported herein assesses occupancy trends spanning up to a total of 70km of distances inside or outside of NNP. Reduced occupancy observed in areas furthest away from humans and their cattle is plausible for *An. arabiensis* given these are their only two known preferred blood sources (17, 24, 25). However, although occupancy of *An. gambiae* complex decline overall, aquatic habitats in camps with higher SNEII scores were also occupied by a second species of the same complex, *An. quadriannulatus,* with increasing abundance relative to *An. arabiensis*.
