## Supplementary material for "Blood host preferences and competitive inter-species dynamics within an African malaria vector species complex inferred from signs of animal activity around aquatic larval habitats": S2_Figure

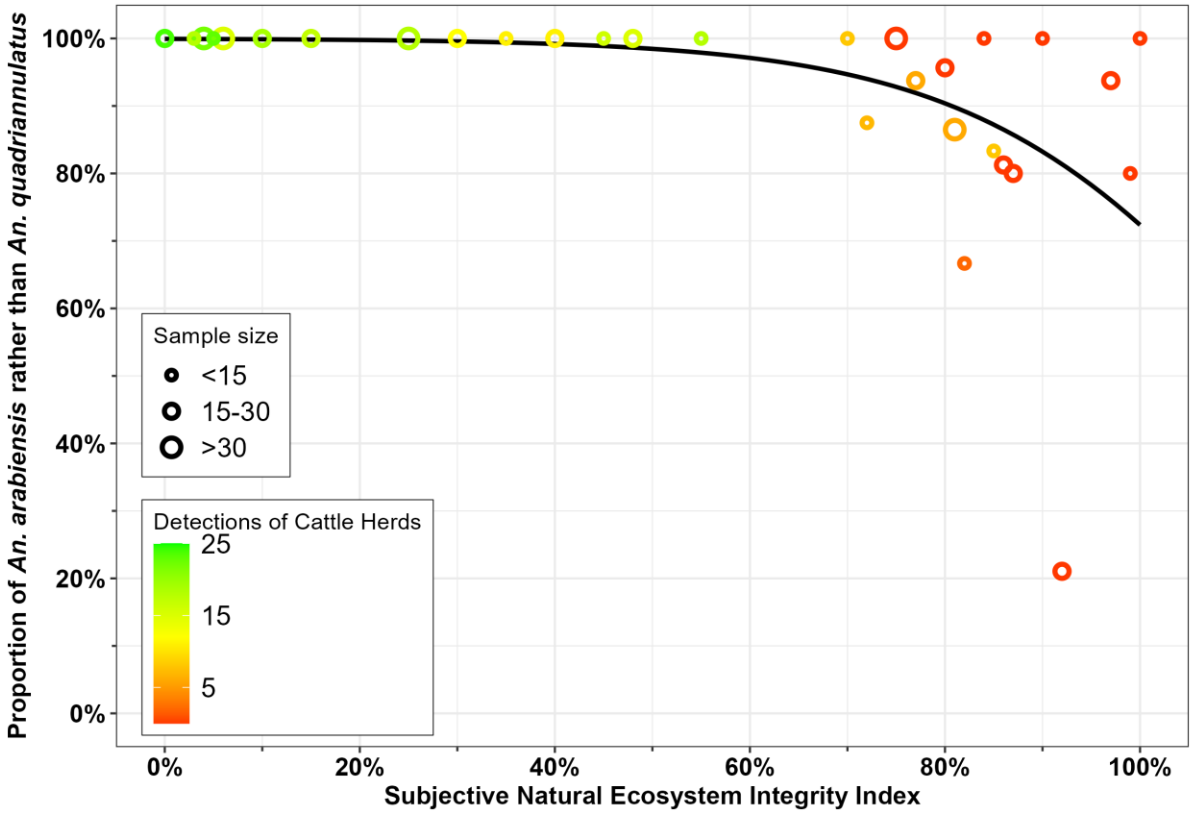


**Figure S2:** A scatterplot graph of the proportion of *An. gambiae* complex $F_0$ adults raised from larvae collected over survey rounds 1 to 3 that were identified as *An. arabiensis* rather than *An. quadriannulatus* by PCR amplification (1). Crude proportions of *An. arabiensis* for each camp are plotted against the Subjective Natural Ecosystem Integrity Index (SNEII; [(2, 3), S6 Appendix]). Sample size and the frequency with which cattle herds were detected are represented by symbol size and colour, respectively. The graph was generated with *ggplot* in R and the trendline was fitted with the *glm* option using the *geom_smooth* command, specifying a binomial distribution and logit link function for the dependent variable.
