## Supplementary material for "Blood host preferences and competitive inter-species dynamics within an African malaria vector species complex inferred from signs of animal activity around aquatic larval habitats": S3_Appendix

**S3 Appendix: Detailed larval survey protocol**

*Survey design, including spatial and temporal constraints placed on sampling effort*

The upper limit of 2km from the camp location was decided upon to prevent geographic overlapping between neighbouring camp locations and to minimise spatial autocorrelation effects. Surveys were intensive and conducted in a challenging climate, where habitats were often in open, unshaded areas, so a time limit of four hours was decided upon to mitigate against investigator fatigue and ensure optimal data collection for the full duration of an entire survey circuit. This upper limit of survey duration was based on a prior experience of the overall project and an initial pilot of the field procedures, both of which indicated that longer surveys would prove unsustainable, with potential deleterious consequences for both data quality and the surveyor's wellbeing, especially given the extreme heat in the afternoon. Starting at the camp, surveys were initiated at the nearest waterbodies known to the VGS, and any other potential aquatic habitat that were seen along the way were also surveyed. Although this purposive sampling was not a randomised approach, it proved practically feasible and maximised the number of habitats sampled around each camp in an efficient manner over the limited time frame of one morning. If the 2km radius was reached before the time limit was up, a different route back to the camp was taken and any potential habitats that were identified on the way were surveyed.

*Characterising Habitats*

The habitat type was identified by assigning a numeric code from 1 to 12 that corresponded to twelve broad categories of water bodies (Table S3.5). Some examples of the various habitat types identified during surveys can be seen below in figure S3.1. Further information including a more detailed description of the potential habitat and the presence or absence of *short vegetation*, *tall vegetation* and *floating vegetation* were recorded by ticking the appropriate column in the row (Table S3.5). Visual estimates for *water depth* and *perimeter* were taken and assigned to their corresponding categories indicated on the form (Table S3.5).

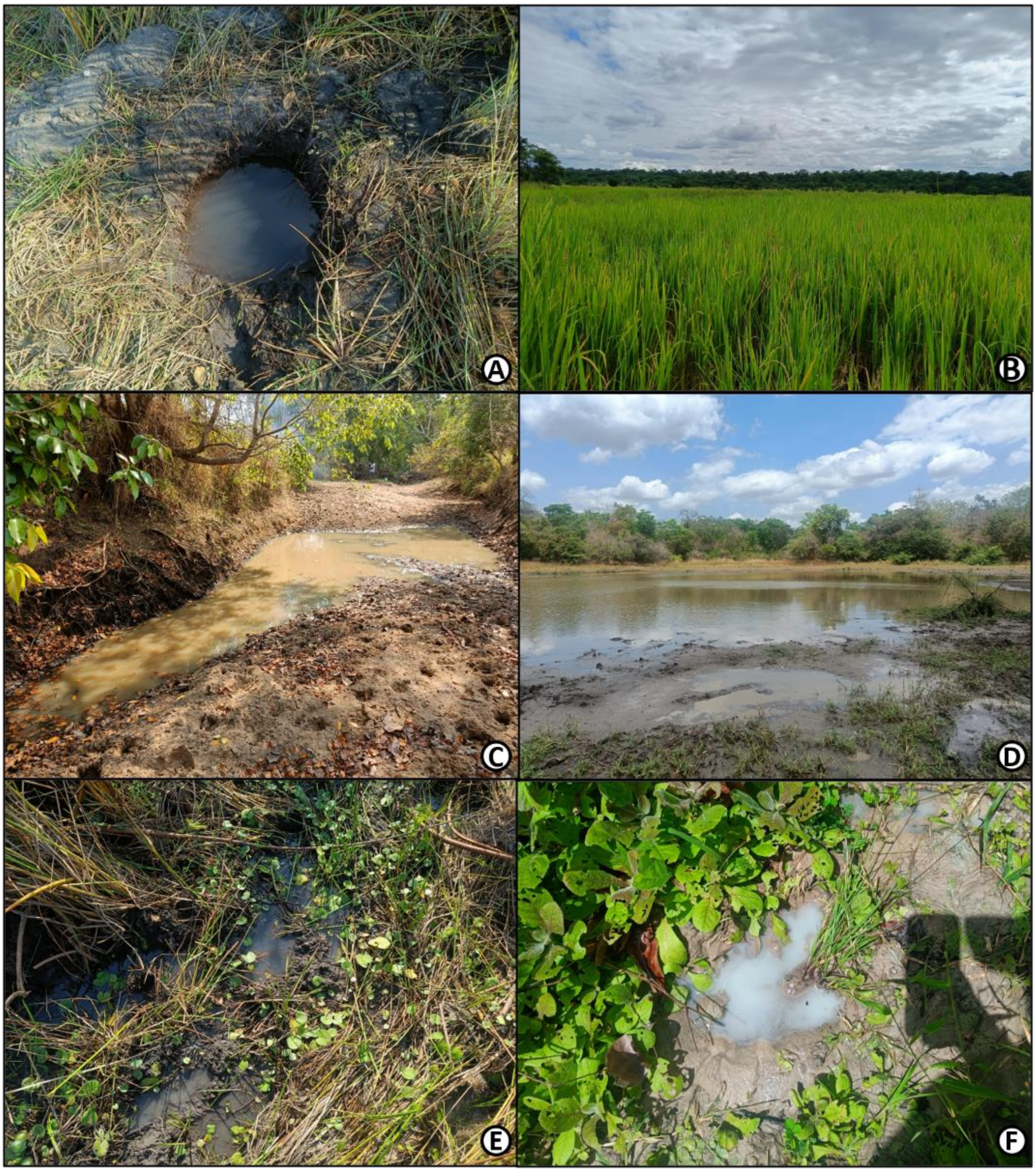


Figure S3.1. Examples of common larval habitat typers that were found during the study, A; human-dug well, B; rice paddy, C; pool along the bottom of a streambed, D; waterhole, E; swampy area, F; animal print (hippo).

*Dipping*

A sample of water, informally referred to in the field as a *dip*, was taken by briefly submerging a standard white 350ml dipper just below the water surface into the habitat at 45-degree angle so that the suction created by water displacement draws both water and larvae into the dipper cup (Figure S3.2), even where obstructing vegetation would frustrate sampling with conventional limnological sampling tools like sweep nets. The dipper was then removed immediately, to prevent captured larvae from escaping.

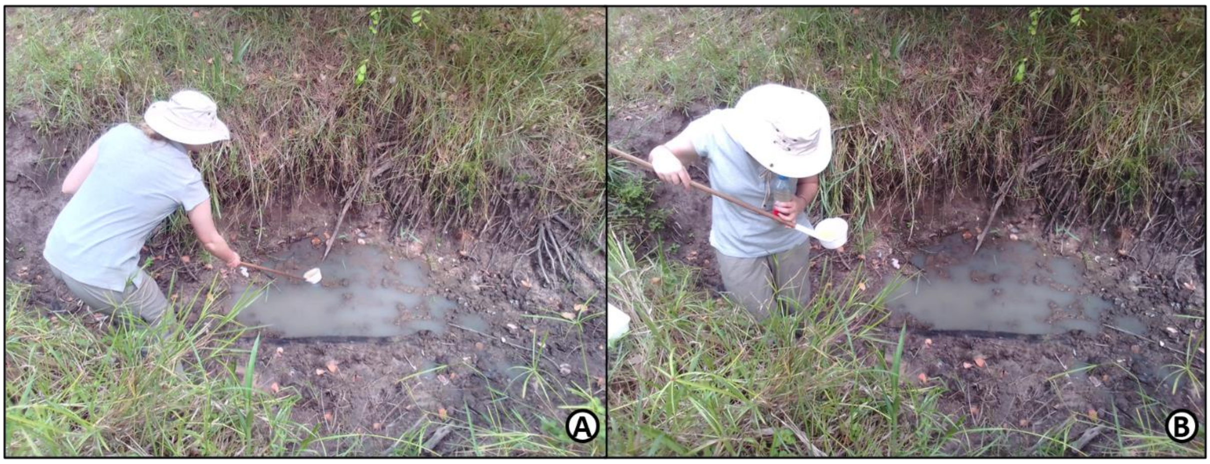


Figure S3.2. Example of the standard dipping technique to sample larvae (1) (A) and inspecting the dipper for anopheline and culicine larvae (B).

All dips were taken along the waterbody perimeter and were methodically numbered and spaced according to estimates of the habitat perimeter and criteria defined as follows. If a potential habitat was <2m in perimeter (Table S3.5), one or two dips were taken depending on size and feasibility. For human, livestock and wild animal prints or any other habitat that was too small to be effectively dipped, a standard 28ml turkey baster was used to suction water and larvae from the habitat into the dipping cup. These habitats were classified as having one dip. For medium sized waterbodies categorised between 2m and 20m, as well as two larger waterbody perimeter categories greater than 20m, and 200m, respectively (Table S3.5), the spacing between each dip was estimated by the number of paces taken by the individual carrying out the survey to ensure dips were taken at regular intervals around the full perimeter.

Aquatic habitats that were categorised as being approximately 2m to 20m in perimeter, were dipped for a total maximum of five times. Therefore, if the habitat was about 10m in perimeter or less, dips were taken once every two paces walked around the perimeter. If the habitat was estimated to be between 10m and 20m, dips were taken once every four paces. For larger waterbodies between 20m and 200m, and those that were greater than 200m, a maximum of 10 dips were taken once every 20 paces and every 50 paces, respectively. These criteria were developed to maximise sampling efficiency, consistency, and productivity after a pilot survey. Even if *An. gambiae* complex larvae (Figure S3.3) were identified immediately, dips were continually taken around the full perimeter or until the maximum number of dips was reached, because as many of these larvae as possible were collected, kept alive, and sent back to Msakamba to be reared for insecticide susceptibility testing (2).

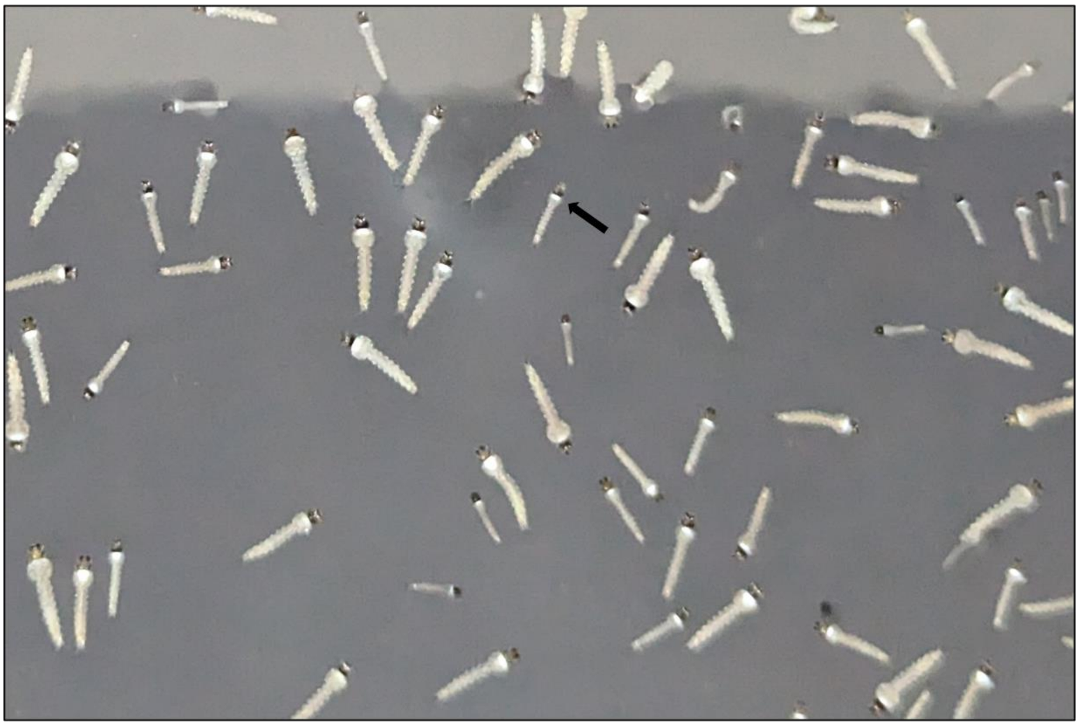

Figure S3.3. An image of *An. gambiae* complex larvae under the microscope as identified by a white collar immediately behind the head, which is a distinguishing morphologic trait of this species complex (3).

*Collection of larvae from aquatic habitats.*

For each dip containing anopheline larvae identified as *An. gambiae* complex (Figure S3.3), individual specimens were taken from the dipper using a clean, disposable, 3ml plastic pipette and placed inside a 500ml capacity water bottle, filled with approximately 350ml of water that was freshly filtered *in situ* at the camp prior to starting the survey. The purpose of filling the bottle approximately three quarters full was to avoid sloshing that could cause mortality or reduce fitness while providing sufficient oxygen. Approximately 1mm wide air holes were punctured in the bottle caps to further supply fresh oxygen. To make the process of transporting larvae back to Msakamba logistically manageable, larvae from several habitats that were similar in appearance and close together were pooled into one bottle, reducing the number of bottles that had to be carried on foot. Habitats that appeared to have exceptionally high numbers of larvae, were collected as a separate *batch* sample so that one bottle contained larvae from one highly productive habitat only. Each sample was labelled with the form serial number and form row (used as a primary key for sample tracing) and a unique sample label code (secondary key), along with the sample type (1; an individual larva, 2; pooled sample, i.e., larvae from multiple habitats, 3; batch i.e., from the same habitat). Once the survey was complete and the team returned to the camp, the water in each bottle was exchanged for newly filtered water and larvae were fed using finely ground TetraMin® fish food. Replacing water from their natural habitat with filtered drinking water was intended to minimise mortality from natural ammonia accumulation (4, 5). Bottles were placed in the shade to avoid overheating from the sun until the following morning, when each sample was provided with fresh water and fish food again and transported back to Msakamba for sample processing in either a customised backpack with the adult mosquitoes (6), or suspended on the end of string handles to prevent splashing which was used later during the study (Figure S3.4).

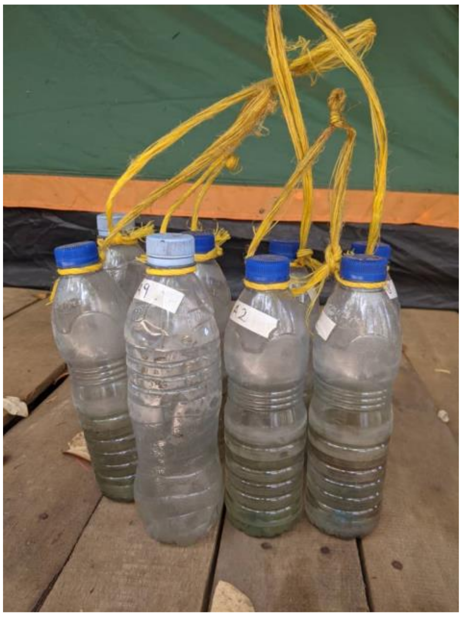

Figure S3.4. Hanging bottle design used later in the study to transport mosquito larvae by foot across long distances (2). This design prevented splashing and improved survival rates compared to using the backpack designed for transporting adults.

All field-identified *An. gambiae* complex larvae from January to November 2022 were reared to adults (known as the F0 generation), which were then fed and propagated through one or two further generations (F1 and F2, respectively), before they were used for insecticide susceptibility testing (2). Adult *An. gambiae* complex mosquitoes that were caught in the wild, but reared in captivity could only be identified by polymerase-chain reaction (PCR) analysis (7) in the Ifakara Health Institute laboratory in Ifakara, after these F0 adults had either already yielded eggs or died. These were used for preliminary analysis only (S2 Figure) and were not included in the final analyses for several reasons. Following the observation that the *An. gambiae* population at many camps consisted of a mixture of *An. arabiensis* and *An. quadriannulatus*, it was decided that PCR tests (7) on the F0 adult mosquitoes cannot give robust representations of sibling species composition. This is because the specimens were handled multiple times during the collection, transportation and rearing phases that could easily lead to survival bias towards the fittest of those two competing species under such shared, artificial environmental conditions. Another limitation was that all anopheline species that were present in an aquatic habitat during the surveys were collected in the same sample and were retained and transported back to the Msakamba insectary in the same bottles, irrespective of whether they had been identified as *An. gambiae* complex or not. Once the samples were returned to the field insectary for the rearing phase, samples of anopheline larvae were then visually re-identified in the field insectary in Msakamba using the same methodological approach and anything not identified as *An. gambiae* complex was removed from the sample during the rearing phase. This procedural flow may have exacerbated potential survival biases arising from overcrowding and predation (8, 9).

It was therefore decided to add a supplementary procedure for collecting more larvae and preserving them *in situ* to address these questions about how species composition varied with location, ecosystem integrity and potential host availability. The purpose of standardising and limiting the number of dips in the primary larval collection protocol was to strike a compromise between the amount of time spent at each waterbody and the number of different habitats that could be surveyed for assessing how occupancy varied across the ecological gradient of the study site. As it was essential to maintain the existing survey protocol without changing it, so that data collection to address the original questions about occupancy and insecticide resistance phenotypes were consistent and comparable, it was decided to supplement those collections of live larvae with more purposive and intensive collections from the same aquatic habitats to obtain larger, more carefully disaggregated samples and preserve them *in situ* for robust PCR analysis (7).

When a habitat was positively identified for the presence of *An. gambiae* complex, the first individual conducting the occupancy survey continued to collected larvae as per the original protocol that involved the live transportation to the field insectary as described above using the hanging bottle design (Figure S3.4). However, during this final round, separate samples of larvae were collected in parallel by a second trained VGS following immediately behind the first surveyor. This second surveyor collected larvae from aquatic habitats that were positively identified during the survey using the same dipping technique but far more exhaustively. Field identified *An. gambiae* complex only were immediately preserved into a 50ml Falcon® tube filled with ethanol using a clean, disposable plastic pipette. Directly preserving larvae right at the aquatic habitat at which they were collected from, eliminated any potential bias or human error that could distort the sibling species composition during the transport and rearing processes and therefore, ensured robust formal species identification. This approach also increased the sample sizes to be used for subsequent analyses as the only focus of the second VGS was to identify and preserve as many *An. gambiae* complex larvae as possible from each aquatic habitat found to contain them. Also, given the specific interest in sibling species composition, and likelihood of strong covariance of individual dip samples with habitats, combined with substantial variance among habitats, it was decided to keep separate, specific samples for each individual habitat, so that anticipated within habitat covariance and between-habitat variance could be accounted for in the statistical analysis as nested random effects.

Unlike the previous surveys, over rounds one to three in 2022, wherein field-identified *An. gambiae* complex larvae from multiple habitats were often pooled together, the larvae from different habitats were all stored as separate batch samples so that each tube contained preserved larvae from a singular aquatic site. A target of ten batch samples per camp was set to enable robust quantification and statistical evaluation of population composition heterogeneity around each camp location. Each habitat-specific batch sample was labelled with the camp number, the serial number and form-row number on the larval surveillance form and given a *breeding site identification number (BSID)* from 1 to 10, so that each batch sample could be traceable to a particular aquatic habitat from a particular camp. These batch samples of preserved field identified *An. gambiae* complex larvae were returned to Msakamba and were transported to the laboratory in Ifakara for PCR analysis (7).

These more carefully separated and clearly distinguished samples were also used to assess how heterogeneities in sibling species composition within the *An. gambiae* complex may vary with the availability of different mammalian species as potential blood sources for the adult mosquitoes that oviposited in those habitats, just as demonstrated for mixed populations of *An. arabiensis* and *An. gambiae s.s.* in fully domesticated environments elsewhere (10, 11). To perform these associations during data analysis, radial surveys of humans, livestock, wildlife activity and land use (12, S8 Appendix) were conducted around each camp in parallel with the larval surveys.

**Table S3.5.** The data entry form used to characterise an aquatic habitat and identify the presence or absence of aquatic stage mosquitoes.

**Survey Round (RND):** _______________ **ED1-SEN:** _______________________

**Form Type (FT):** EDSO1; Larval habitat survey for presence of aquatic mosquitoes and environmental attributes **Date (DT):**____/____/______ **Form Serial Number (SEN):**___________________

**Survey or Experiment Identifier (SEI):** _______ **Cluster/Camp (CR):**_________ **Camp Name or Description (CND):**_______________________________________________________________

**GPS Identifier (GID):**_________ **Camp GPS Way Point Number (CGWPN):**___________ **Camp GPS Coordinates:** Southing (CGCDS);___________________ Easting (CGCDE)_____________________

**Camp Comments (CC):** ____________________________________________________________________________________________________________

**Habitat type (HT) codes:**

1: Small puddles, tracks and depressions
2: Swampy areas, seepages and springs
3: Flowing stream bed
4: Flooded valley
5: Pond or water hole
6: Artificial drain or ditch
7: Wells and other human-made holes
8: Water storage & other artificial containers
9: Rice paddy
10: Ridge and furrow agriculture (Matuta)
11: Other agriculture
12: Others (describe)

| Form Row (FR) | Habitat GPS Way Point Number (HGWPN) | Habitat Type (HT) | Habitat Description (HD) | Habitat perimeter (HPER) | | | | Number of Dips (ND) | Emerging or Floating Vegetation (EFV) | | | | Water Depth (WD) | | | Aquatic Stage (AS) | | | | | | | | | Sample Label Code (SLC) | Sample Type (ST) (Ind=1, Batch=2, Pool=3) | Habitat Comments (HCM) |
|---|---|---|---|---|---|---|---|---|---|---|---|---|---|---|---|---|---|---|---|---|---|---|---|---|---|---|---|
| | | | | | | | | | | | | | | | | Anopheles | | | | Culex | | | Pupae | | | | |
| | | | | < 2m(HPER=1) | < 20m (HPER=2) | 20-200m (HPER=3 | > 200m (HPER=4) | | Absent (EFVA) | Short (EFVS) | Tall (EFVT) | Floating (EFVF) | <0.1m (WD=1) | 0.1-0.5m (WD=2) | >0.5m (WD=3) | Absent (ASAA) | Early (ASAE) | Late (ASAL) | Gambiae present (ASAGP | Absent (ASCA) | Early (ASCE) | Late (ASCL) | Absent (ASPA) | Present (ASPP) | | | |
| A | B | C | D | E | F | G | H | I | J | K | L | M | N | O | P | Q | R | S | T | U | V | W | X | Y | Z | AA | AB |
| 1 | | | | | | | | | | | | | | | | | | | | | | | | | | | |
| 2 | | | | | | | | | | | | | | | | | | | | | | | | | | | |
| 3 | | | | | | | | | | | | | | | | | | | | | | | | | | | |
| 4 | | | | | | | | | | | | | | | | | | | | | | | | | | | |
| 5 | | | | | | | | | | | | | | | | | | | | | | | | | | | |
| 6 | | | | | | | | | | | | | | | | | | | | | | | | | | | |
| 7 | | | | | | | | | | | | | | | | | | | | | | | | | | | |
| 8 | | | | | | | | | | | | | | | | | | | | | | | | | | | |
| 9 | | | | | | | | | | | | | | | | | | | | | | | | | | | |
| 10 | | | | | | | | | | | | | | | | | | | | | | | | | | | |
| 11 | | | | | | | | | | | | | | | | | | | | | | | | | | | |
| 12 | | | | | | | | | | | | | | | | | | | | | | | | | | | |
| 13 | | | | | | | | | | | | | | | | | | | | | | | | | | | |
| 14 | | | | | | | | | | | | | | | | | | | | | | | | | | | |
| 15 | | | | | | | | | | | | | | | | | | | | | | | | | | | |
| 16 | | | | | | | | | | | | | | | | | | | | | | | | | | | |
| 17 | | | | | | | | | | | | | | | | | | | | | | | | | | | |
| 18 | | | | | | | | | | | | | | | | | | | | | | | | | | | |
| 19 | | | | | | | | | | | | | | | | | | | | | | | | | | | |
| 20 | | | | | | | | | | | | | | | | | | | | | | | | | | | |
