## Supplementary material for "Blood host preferences and competitive inter-species dynamics within an African malaria vector species complex inferred from signs of animal activity around aquatic larval habitats": S4_Appendix

**S4 Appendix: Validation of field identification methodology for larvae from the *Anopheles gambiae* complex**

***Methods***

*Anopheles* larvae encountered during the larval occupancy surveys were positively identified in the field as members of the *An. gambiae* complex if a white collar immediately behind the head was visually observed. Therefore, to validate the field methodology used and to enable face-value interpretation of the occupancy rates recorded in the field and analysed by regression using the generalised linear mixed model (GLMM) described in S10 Appendix, positive PCR amplification rates from field identified larvae were fitted to a GLMM using the *glmm* package in R.

Habitat-specific batch samples of collected larvae from the fourth round either amplified as members of the *An. gambiae* complex or did not amplify at all, hence the proportion of field identified larvae that positively amplified as *An. gambiae* complex was treated as a binomial outcome and weighted by the total number of mosquitoes from that were tested by PCR (1). The BSID nested within camp number was treated as a random effect to account for any covariance within habitats and camp locations. Distance, SNEII, and historical landcover type were all assessed as univariate fixed effects to account for any spatial or environmental effects on the amplification rates of *An. gambiae* complex that might be associated with them, but none proved significant. Consequently, a GLMM with an intercept only and no other fixed effects was used to calculate an overall mean amplification rate for *An. gambiae* complex (G) with 95% confidence intervals (CIs) across all camps, using the following formulae, $G = \frac{e^{\beta_0}}{1+e^{\beta_0}}$ with 95% CIs = $\frac{e^{\beta_0 \pm 1.96\,(\sigma_0)}}{1+e^{\beta_0 + \sigma_0}}$, respectively, where $\beta_0$ is the intercept and $\sigma_0$ is the standard error of the mean for the intercept. To illustrate how consistently high these results were regardless of where samples were obtained from, rates of successful amplification of *An. gambiae* complex at each camp were displayed as a scatter plot with the points distributed across a horizontal axis that represents the distance of the camp from the nearest NNP boundary or the nearest human settlement inside the park (km) on a square root scale. The predicted means and confidence intervals for this plot, were calculated based on a second GLMM that included distance as a fixed effect as well as the intercept. Using the outputs from this GLMM the predicted mean (G) and a combined standard error of the means (σ) for each camp were calculated in Microsoft Excel® as $G = \frac{e^{\beta_0+\beta_1 X_{1,i}}}{1+e^{\beta_0+\beta_1 X_{1,i}}}$, and $\sigma = \sqrt{{\sigma_0}^2 + \sigma_1 {x_{1,i,}}^2}$, respectively, where $\beta_0$ is the intercept, $\beta_1$ is the effect size attributed to distance, $x_{1,i}$ represents the values of distance at camp $i$, and $\sigma_0$ and $\sigma_1$ are the standard errors of the means for $\beta_0$ and $\beta_1$, respectively. The 95% CI for the predicted means at each camp were then calculated as $G = \frac{e^{\beta_0+\beta_1 X_{1,i} \pm 1.96 * \sigma}}{1+e^{\beta_0+\beta_1 X_{1,i} \pm 1.96 * \sigma}}$. The predicted means and confidence intervals were then plotted using the *geom_line* and *geom_ribbon* functions, respectively.

***Results***

The DNA of 2,468 field-identified *An. gambiae* complex larvae taken from 286 aquatic habitats across 39 camps were tested for species identity by PCR (1). The number of aquatic habitats at each camp that larvae were collected from ranged from 1 to 10, with an overall average of 7 habitats per camp. The *a priori* target of 10 aquatic habitats per camp was not reached at some camps when there was a limited availability of occupied habitats that contained what appeared to be *An. gambiae* complex larvae. Camp number 29, *Kambi ya Simba*, was surveyed for occupancy but there were no available batch samples to be identified to species level by PCR (1)

The overall mean amplification rate of 96.6% [94.0%, 96.8%] (Figure S4), was very high, and an amplification of 5% failure rate would be normally considered excellent, even for insectary reared positive controls. Neither distance (OR = 1.09, P = 0.585), nor ecosystem integrity (OR = 0.99, P = 0.922), nor historical landcover type (OR = 1.01, P = 0.794) had any effect on amplification rates, indicating that any larva collected under these varying geographic, and environmental conditions can be confidently identified in the field as *An. gambiae* complex if a single white stripe is evident immediately behind the head. It also validates the results fitted by the GLMM which predicts *An. gambiae* complex occupancy rates across ecosystem integrity and specific habitat attributes (S1 Appendix, Table S1.2). Above all, it concludes that the field identification approach proved to be very reliable, with the exception of one outlier (Funga, camp 26 in Figure 1) located approximately 17km from the NNP boundary (Figure S4).

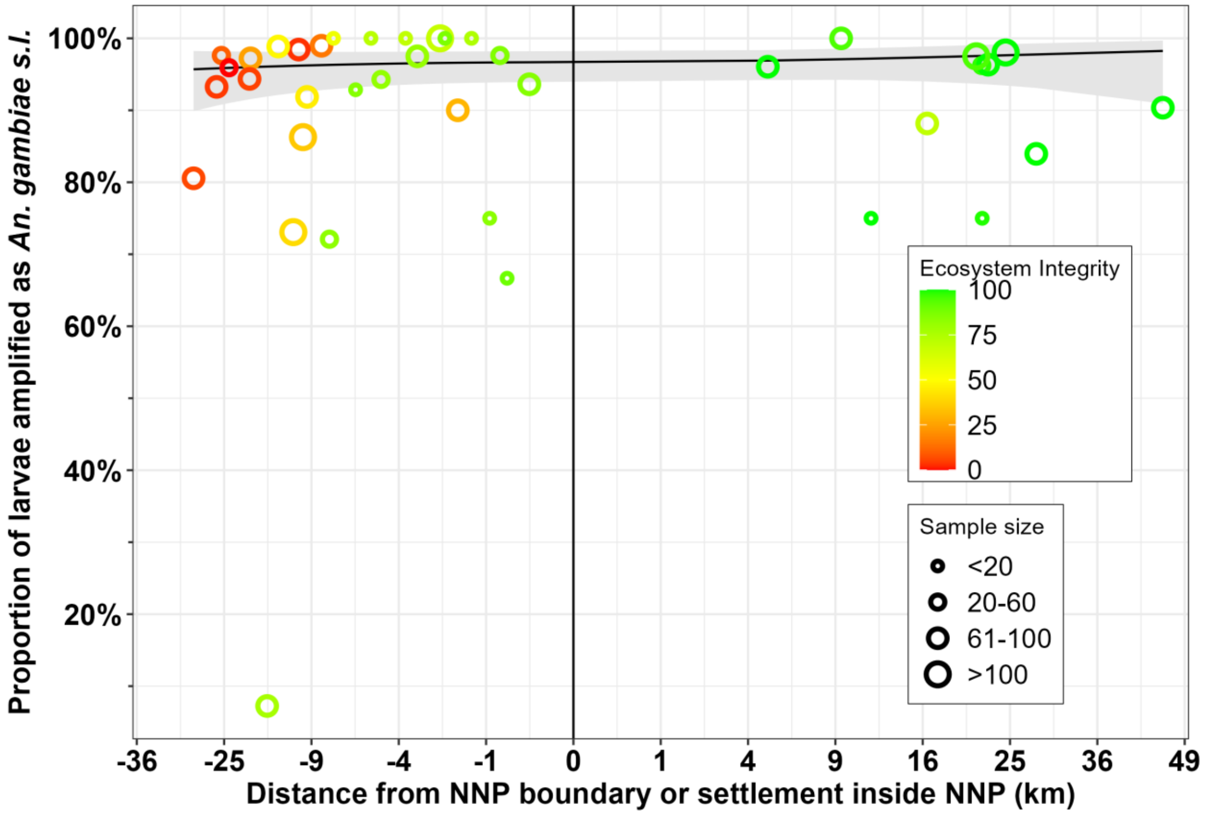


**Figure S4. A graphical illustration of how the proportion of field-identified larvae that successfully amplified as *An. gambiae* complex did not vary with SNEII or distance from the nearest point to the NNP boundary or the nearest human settlement within the park.** Each point represents the proportion of field-identified larvae confirmed to be *An. gambiae* complex by PCR (1) across all batch samples collected at each camp. The colour and size of each point represent the SNEII and the total sample size for each camp, respectively. Predicted means and confidence intervals were calculated from a GLMM fitted to the proportion of specimens that successfully amplified, using a binomial distribution and logit link for this binary dependent variable in which the non-significant effect of distance was treated as the only fixed effect with breeding site identification number nested within camps treated as random effects.
