## Supplementary material for "Blood host preferences and competitive inter-species dynamics within an African malaria vector species complex inferred from signs of animal activity around aquatic larval habitats": S5_Appendix

### S5 Appendix: Study area and sampling frame

#### *Study area*

This study was conducted in the Kilombero valley in the Morogoro region of southern Tanzania, which is recognised as East Africa's largest wetland at low altitude (1, 2). The valley is part of the Rufiji River Catchment Basin and is marked by the Udzungwa mountains in the north and the Mahenge mountains in the south (2). The wetlands of the Kilombero are a unique and biodiverse ecosystem, within 7,950 square kilometres have been designated as a Ramsar site since 2002 (2) (Figure S5.1). As well as being of high ecological and hydrological importance, the wetlands and surrounding area are of major economic significance, supporting a variety of agricultural, forestry and fishing livelihoods (1).

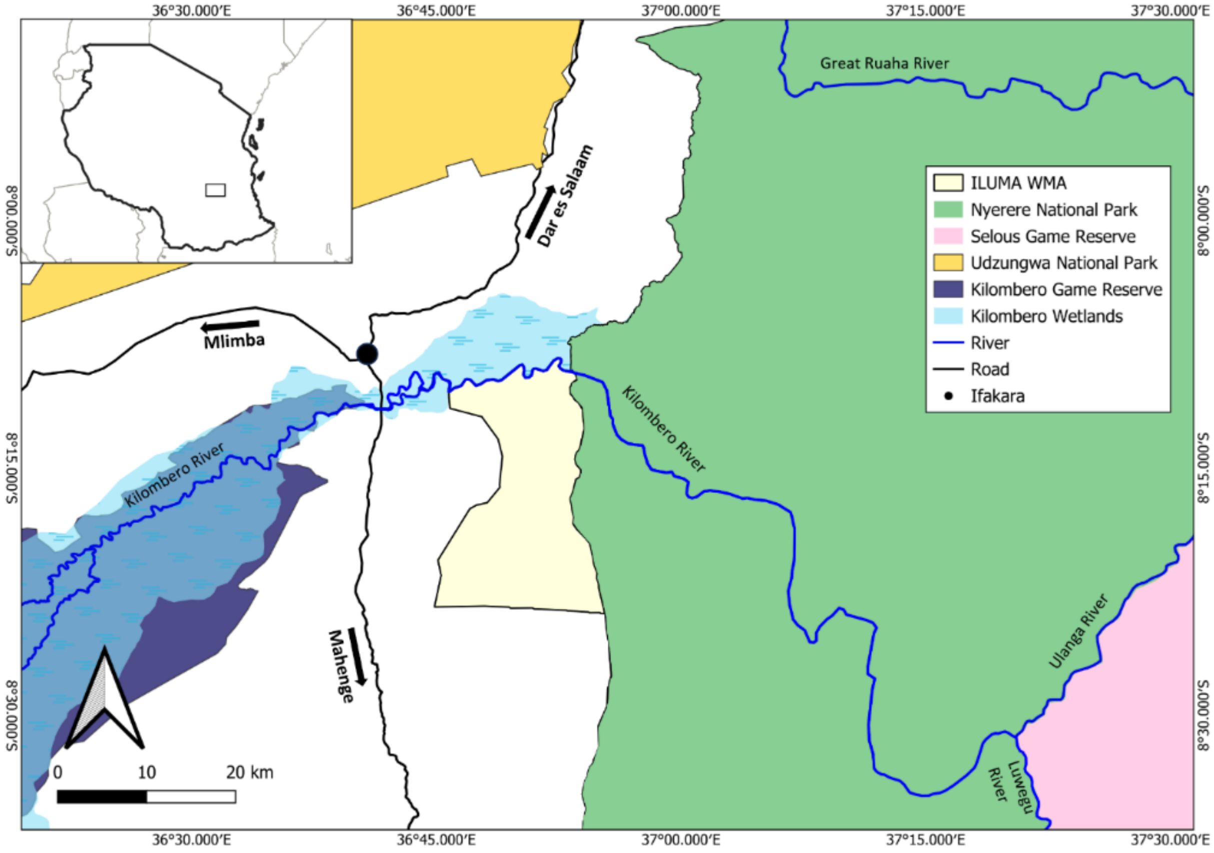


**Figure S5.1:** Map showing the Kilombero wetlands and the study area of the ILUMA Wildlife Management Area and Nyerere National Park.

The region has what are referred to as a short and a long rainy season, from approximately November to February and from March to April, respectively. The short rains are typically intermittent, sporadic and unreliable, whereas the long rains are usually more predictable and consistent. The dry season generally lasts from June to December, during which time the area experiences little to no rainfall. The malaria transmission systems and vector populations in the Kilombero valley have been exceptionally well-characterised (3-16), and malaria vector research continues to progress with the Ifakara Health Institute being based in Ifakara town, with recent studies from (17-22), to name a few. Today, the two main vectors include *An. funestus* Giles and *An. arabiensis* Patton (16, 18, 19, 23).

Located south-east of Ifakara town in the Kilombero and Ulanga districts, lies more than 500km$^2$ of community lands under the stewardships of 15 villages which is collectively managed through the Ifakara-Lupiro-Mangula Wildlife Management Area (ILUMA WMA) (Figure S5.1). It was formally established as a wildlife management area in 2015 for the purpose of practicing sustainable land use and promoting wildlife conservation in the buffer zone between a rigorously conserved state-protected area (formerly the Selous Game Reserve, now Nyerere National Park) and village communities, while also creating opportunities for local communities to generate income and advance rural development (24). Miombo woodland is the predominant natural land cover type across most of the ILUMA WMA, although a sizeable trait of dense groundwater forest stretches along the south-bank of the Kilombero River. On the north-bank, floodplain grassland extends to the rainforest covered Udzungwa mountains. These very distinct ecosystems support sizeable populations of wild herbivores and carnivores, including iconic species like elephant, African buffalo, lion, leopard and African wild dogs.

The western boundary of ILUMA WMA borders lands designated for agriculture and grazing extending as far as the main road connecting Ifakara and Mahenge (Figure S5.1). Due to its proximity to domesticated land and human activity, many areas within ILUMA WMA close to the western and southern boundaries are vulnerable to unauthorised exploitation by humans, and large sections of these areas have even been cleared for agriculture, grazing and settlement (25). Other types of unauthorised human disturbance occurring across the designated conservation area, include timber harvesting, charcoal burning, fishing, and game meat hunting. However, as one moves further into ILUMA WMA and away from human settlement, natural ecosystems become less exposed to human disturbance and are increasingly inhabited by abundant populations of wild mammals that offer mosquito populations diverse alternatives to humans and livestock as potential blood sources.

The eastern boundary of ILUMA WMA is marked by a road running in a north-south direction along its border with Nyerere National Park (NNP) (Figure S5.1). Established in 2019, NNP is Tanzania's largest and newest national park. NNP was formerly a part of the Selous Game Reserve and covers an area of more than 30,000 square kilometres of fully intact wilderness (26). Presently, most tourist activities and development occur in the northern section of the park, along the Rufiji River between Ikwiriri and Kisaki, that have been previously used for photographic tourism rather than big game hunting. At the time of the study, however, the western area of NNP that borders ILUMA WMA had not yet been extensively developed, so it experiences minimal human activity. The miombo woodlands of ILUMA WMA, extend eastward beyond the border road and well into the national park (Figure S5.1), providing an extensive habitat for the same species that are found in ILUMA WMA. Additionally, open grasslands, acacia scrub and undisturbed floodplains in NNP favour species that are not regularly seen in ILUMA WMA, such as impala, waterbuck and kudu.

This large study area was fundamental for addressing the objectives of this specific study and the broader goals of the overall project, as it represents a geographical gradient of natural ecosystem integrity ranging from fully domesticated land uses in the west to completely intact natural ecosystems to the east. Agricultural land and permanent villages to the west of ILUMA WMA offer typical larval habitats for mosquitoes of the *An. gambiae* complex, including early-stage rice paddies, ridge and furrow agriculture, livestock hoofprints, human-made tracks and artificial wells and drains (27-29). In the wild areas of eastern ILUMA WMA and in NNP, waterholes, prints of various wild animals and seasonal pools were considered as water bodies that could provide potential breeding sites. Additionally, receding groundwater levels during the dry season cause streams to stop flowing leaving small, stagnant isolated pools of groundwater within streambeds that can represent highly productive larval habitats (28, 30).

This comprehensive range of de facto land use practices and ecosystem characteristics were also considered to be indicative of the potential host species available to host-seeking *An. arabiensis* adults as potential sources of blood. Since cattle and humans are the known preferred blood source of *An. arabiensis* (31, 32), the abundance of this vector in and around villages may be readily rationalised because these two host species are plentiful in settlements and surrounding domesticated land. At the other end of the gradient, it has been suggested that wild game animals, particularly bovids, may act as suitable alternative hosts for wild *An. arabiensis* populations in wilderness areas (8, 33-35), such as the well conserved areas of ILUMA WMA and NNP. As WMAs act as buffer zones between national parks and permanently inhabited areas, wildlife, humans and cattle could all represent possible blood sources for *An. arabiensis* inside the ILUMA WMA. Furthermore, ILUMA WMA encompasses varying degrees of human disturbance and so the availabilities of cattle, human and wildlife species as potential blood sources was expected to vary considerably across fine geographical scales. Given the known blood-feeding behavioural plasticity amongst *An. arabiensis* populations and the anticipated variability in host species availability provided by the study area, it was therefore expected that these heterogeneities could be interpreted as a portfolio effect that stabilises vector populations (36).

***Sampling sites, study design and logistical field procedures.***

Due to transport constraints such as inaccessibility of many of the camps to vehicles during the wet season, the whole data collection process was completed on foot except for the camps located up to 131 km inside NNP, that were not originally planned *a priori*, but rather towards the end of the study as a post-hoc adaptive response to emerging results. These NNP camps were accessed via the Kilombero River using a motorboat in the wet season, and by car via established roads during the dry season. Additionally, transect surveys of land use and the activities of humans, livestock and wildlife were completed along the routes walked between camps within a survey circuit, and so these transits on what were referred to as *moving days*, needed to be completed on foot. Because of these challenges, the development of novel, logistically feasible procedures for collecting robust data and large numbers of live mosquito specimens, while also ensuring the safety of team members in a remote and challenging environment, were fundamental for the entire study framework.

A fenced camp with essential basic infrastructure like thatch roofs, tables, chairs, large tents, a kitchen and solar-powered electricity supply was established as the hub for all scientific and logistical processes in the field and named *Msakamba* (camp 1 in figure S5.2 and table S5.3). The central location of Msakamba (Figure 1) made it possible to reach any camp in ILUMA WMA within a day's walk, which ensured that live adult and larvae samples could usually be returned in good condition within 48 hours of collection. It also ensured that food supplies, as well as recharged batteries and power packs for mobile phones and field equipment could be regularly delivered to the mobile field team moving from camp to camp every two days. Village game scouts (VGS) recruited from stakeholder communities that are responsible for patrolling and protecting the ILUMA WMA, and for escorting any visitors to the conservation area, were also engaged as essential team members for this study. Msakamba was occupied and maintained on a permanent basis by a team of VGS and technicians, who were responsible for maintaining the field insectary, rearing field-caught mosquitoes and carrying out experimental assessments on their insecticide resistance phenotypes.

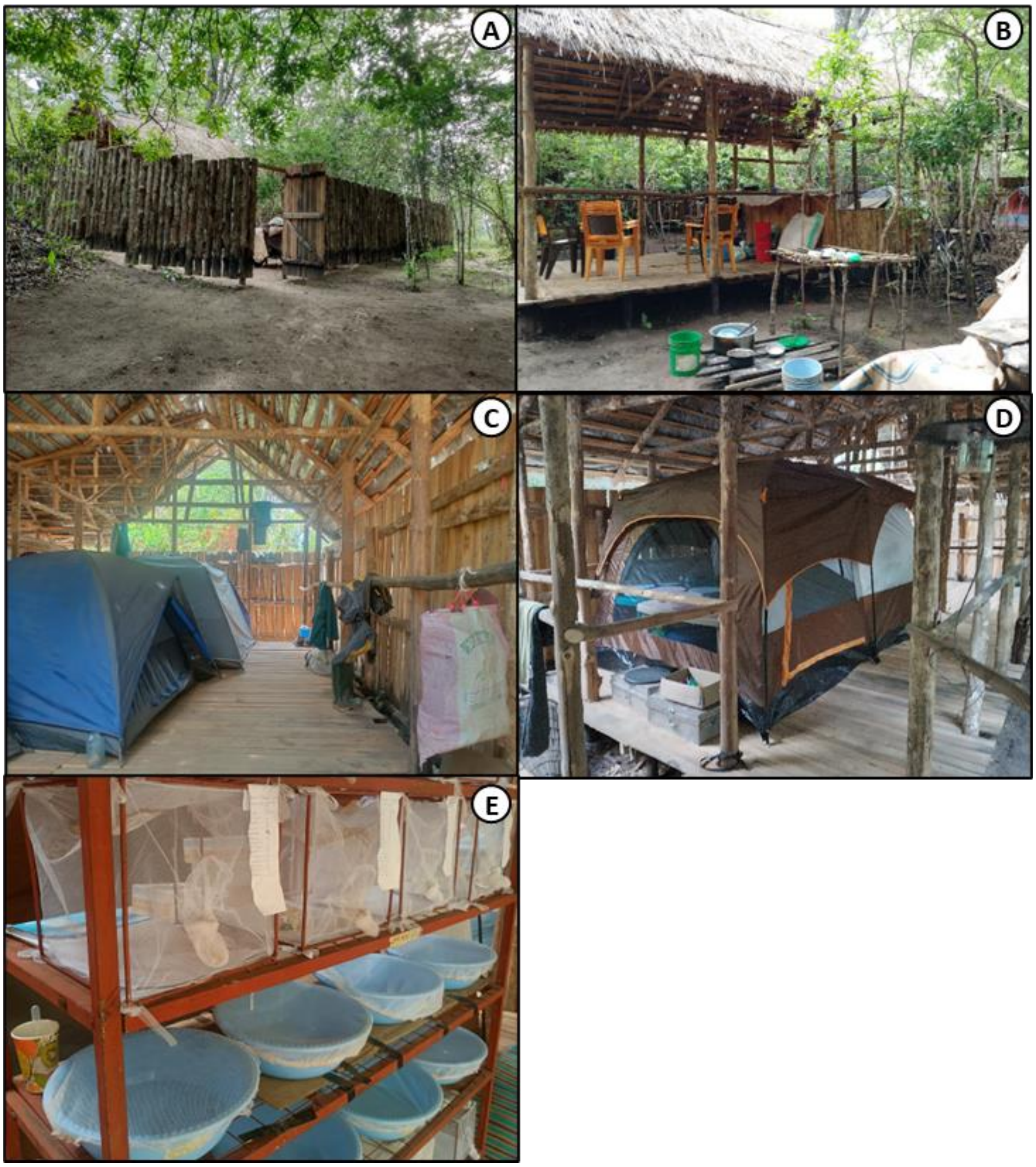


**Figure S5.2:** Illustrative photographs of the central camp established for the project at Msakamba. **A**: The surrounding fence for security, **B**: The kitchen and food storage area, **C**: Tents for sleeping in, **D**: A field insectary tent for housing mosquito adults and larvae, **E**; Mesh cages and plastic water basins for respectively rearing adult and larvae in the field insectary.

For the original protocol, after scouting potential locations in across ILUMA and considering the suggestions of the VGS based on their vast personal knowledge of the area, a total of 28 camps were selected inside the WMA and in or near the villages immediately to the west of the WMA border (Figure S5.1, Table S5.3). The broad geographic distribution of the camps (Figure 1) was planned to encompass as wide of a range of ecosystem states as possible, by including all parts of ILUMA WMA and the neighbouring domesticated land to the west (Figures S5.5 to S5.7). However, the exact position of a camp location was ultimately determined by the requirement of perennial surface water that *Anopheles* larvae could be collected from. Crucially, the availability of water for cooking, drinking (after filtering as seen in Figure S5.4) and bathing was essential for sustaining the field team in the absence of regular vehicle support.

**Table S5.3:** Number, name, location, coordinates, and ecological characteristics of each camp location, together with the quadrant circuit to which it was assigned and the number of times it was surveyed (25, 37).

| Number | Name | Circuit | Location | Coordinates | Number of times surveyed | Habitat Type |
|---|---|---|---|---|---|---|
| 1 | Msakamba | Southeast | Inside ILUMA WMA | -8.25981S, 36.85936E | 3 | Recovering miombo woodland along a small seasonal stream bed. |
| 2 | Msiba wa Deo | Southeast | Inside ILUMA WMA | -8.25821S, 36.88524E | 3 | Intact miombo woodland along a small seasonal stream bed. |
| 3 | Bwawa la Nyati | Southeast | Inside ILUMA WMA | -8.29598S, 36.88732E | 3 | Intact miombo woodland surrounding a large waterhole. |
| 4 | Bwawa la Nandete | Southeast | Inside ILUMA WMA | -8.3467S, 36.9024E | 3 | Degraded miombo woodland surrounding a large waterhole. |
| 5 | Korongo la Bundu | Southeast | Inside ILUMA WMA | -8.37768S, 36.90163E | 3 | Degraded miombo woodland along a small seasonal stream bed. |
| 6 | Bwawa la Namamba | Southeast | Inside ILUMA WMA | -8.34931S, 36.81653E | 3 | Degraded miombo woodland surrounding a large waterhole. |
| 7 | Bwawa la Chakacheni | Southeast | Inside ILUMA WMA | -8.31649S, 36.81507E | 3 | Degraded miombo woodland surrounding a large waterhole. |
| 8 | Bwawa la Njuju | Northeast | Inside ILUMA WMA | -8.24204S, 36.86268E | 3 | Intact miombo woodland surrounding a large waterhole. |
| 9 | Bwawa la Chamvi | Northeast | Inside ILUMA WMA | -8.22176S, 36.89241E | 2 | Intact miombo woodland surrounding a large waterhole. |
| 10 | Bwawa la Miembeni | Northeast | Inside ILUMA WMA | -8.22401S, 36.87518E | 1 | Mostly intact miombo woodland surrounding a waterhole. |
| 11 | Kisima cha Seba | Northeast | Inside ILUMA WMA | -8.20749S, 36.85738E | 2 | Intact miombo woodland adjacent to a small waterhole. |
| 12 | Bwawa la Maya | Northeast | Inside ILUMA WMA | -8.19492S, 36.88045E | 3 | Transition zone between miombo woodland and groundwater forest surrounding a large waterhole. |
| 13 | Bwawa la Mrope | Northeast | Inside ILUMA WMA | -8.16321S, 36.88341E | 2 | Dense groundwater forest surrounding a large waterhole. |
| 14 | Mikeregembe | Northeast | Inside ILUMA WMA | -8.15048S, 36.87539E | 3 | Authorised fishing camp on the banks of the Kilombero river, open but bordering groundwater forest. |
| 15 | Mdalangwila | Northeast | Inside ILUMA WMA | -8.16041S, 36.83887E | 3 | Authorised fishing camp on the banks of the Kilombero river, open |

| | | | | | | |
|---|---|---|---|---|---|---|
| | | | | | | but bordering groundwater forest. |
| 16 | Bwawa la Muamachi | Southwest | Village outside western boundary of ILUMA WMA | -8.38785S, 36.74629E | 3 | Village outside the conservation area surrounding a large waterhole. |
| 17 | Tuliza Moyo | Southwest | Village outside western boundary of ILUMA WMA | -8.35954S, 36.74821E | 3 | Village outside the conservation area along a small seasonal stream bed. |
| 18 | Mavimba Porini | Southwest | Village outside western boundary of ILUMA WMA | -8.33982S, 36.74628E | 3 | Village outside the conservation area along a large river and surrounding a large waterhole. |
| 19 | Bwawa la Selesussi | Southwest | Inside ILUMA WMA | -8.33064S, 36.76951E | 3 | Degraded miombo woodland surrounding a large waterhole. |
| 20 | Makingi | Southwest | Village outside western boundary of ILUMA WMA | -8.27593S, 36.74737E | 3 | Village outside the conservation area along a small seasonal stream bed. |
| 21 | Bwawa la Mpunga | Southwest | Inside ILUMA WMA | -8.27426S, 36.81946E | 3 | Highly degraded miombo woodland surrounding a large waterhole. |
| 22 | Kisaki | Northwest | Village outside western boundary of ILUMA WMA | -8.24595S, 36.7669E | 2 | Largest human settlement, at the base of a large hill adjacent to a spring and large waterhole. |
| 23 | Uwanja wa Ndege | Northwest | Village outside western boundary of ILUMA WMA | -8.23448S, 36.80218E | 2 | Human settlement along a small seasonal river. |
| 24 | Bwawa la Mkwajuni | Northwest | Inside ILUMA WMA | -8.21681S, 36.81141E | 2 | Intact miombo woodland surrounding a degraded large waterhole. |
| 25 | Bwawa la Mamba Luhogi | Northwest | Inside ILUMA WMA | -8.19352S, 36.78527E | 2 | Intact miombo woodland surrounding a highly degraded large waterhole. |
| 26 | Funga | Northwest | Inside ILUMA WMA | -8.16924S, 36.776E | 2 | Authorised fishing camp on the banks of the Kilombero river, open but bordering intact groundwater forest. |
| 27 | Bwawa la Mlenda | Northwest | Inside ILUMA WMA | -8.20037S, 36.82132E | 2 | Intact miombo woodland surrounding a large waterhole. |

| | | | | | | |
|---|---|---|---|---|---|---|
| 28 | Bwawa la Semka | Northwest | Inside ILUMA WMA | -8.20436S, 36.84023E | 2 | Largely intact groundwater forest surrounding a large waterhole. |
| 29 | Bwawa la Simba | Boma Ulanga | Nyerere NP (East of ILUMA WMA) | -8.31298S, 36.94381E | 1 | Transition zone of mixed miombo woodland and acacia savanna surrounding a waterhole. |
| 30 | Kiboko Zanzibar | Boma Ulanga | Nyerere NP (East of ILUMA WMA) | -8.26313S, 37.00293E | 1 | Open acacia savanna on the banks of the Kilombero river. |
| 31 | Zanzibar | Boma Ulanga | Nyerere NP (East of ILUMA WMA) | -8.2565S, 36.98337E | 1 | Transition zone of mixed miombo woodland and acacia savanna on the banks of the Kilombero river. |
| 32 | Bwawa la Moto | Boma Ulanga | Nyerere NP (East of ILUMA WMA) | -8.2741S, 36.93866E | 1 | Transition zone of mixed miombo woodland and acacia savanna surrounding a waterhole. |
| 33 | Kambi ya Mamba | Kilombero | Nyerere NP (East of ILUMA WMA) | -8.18872S 36.89856E | 1 | Acacia savanna on the banks of the Kilombero river. |
| 34 | Kambi ya Machuma | Kilombero | Nyerere NP (East of ILUMA WMA) | -8.30422S 37.11265E | 1 | Acacia savanna on the banks of the Kilombero river. |
| 35 | Serengeti Ndogo | Kilombero | Nyerere NP (East of ILUMA WMA) | -8.28739S 37.09637E | 1 | Acacia savanna on the banks of the Kilombero river. |
| 36 | Kambi ya Makutano | Kilombero | Nyerere NP (East of ILUMA WMA) | -8.40158S 37.17879E | 1 | Acacia savanna on the banks of the Kilombero river. |
| 37 | Kambi ya Mawe | Kilombero | Nyerere NP (East of ILUMA WMA) | -8.40511S 37.14682E | 1 | Acacia savanna on the banks of the Kilombero river. |
| 38 | Shughuli Kubwa | Msolwa | Nyerere NP (East of ILUMA WMA) | -8.51759S 37.3391E | 1 | Miombo woodland at the Kilombero, Ulanga and Luwegu river confluences |
| 39 | Bwawa la Chatu | Msolwa | Nyerere NP (East of ILUMA WMA) | -8.01597S 37.19808E | 1 | Transition zone of mixed miombo woodland and acacia savanna |
| 40 | Bwawa la Umeme | Msolwa | Nyerere NP (East of ILUMA WMA) | -7.9385S 37.77394E | 1 | Acacia savanna at the Great Ruaha, Rufiji and Ulanga river confluences. |

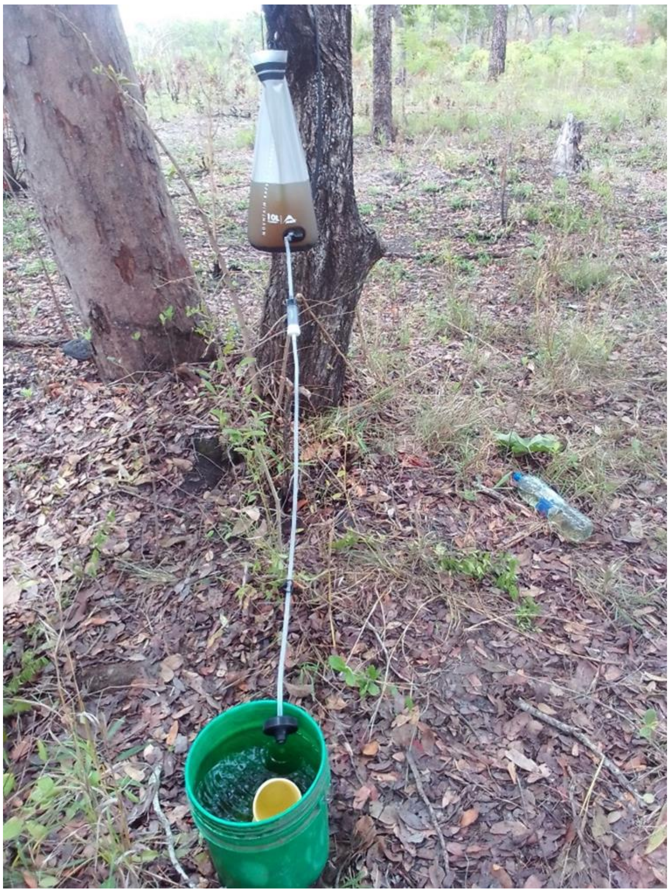

**Figure S5.4:** Filtering water from a nearby stream using the MSR Autoflow™ XL Gravity Filter 10L.

In most cases, the selection of camp locations with accessible surface water also ensured that firewood and at least some shade were available *in situ*. Accessibility by foot during the wet season was also a critical factor to consider, to ensure that each camp could be safely reached, and the transport of live mosquitoes could be completed even during periods of heavy rain and flooding. The presence of one or more glades or valleys with numerous perennial waterbodies, like waterholes, ponds and streambeds near the camp was also a requirement of the methodology used for collecting mosquito larvae (S3 Appendix) and adults (39).

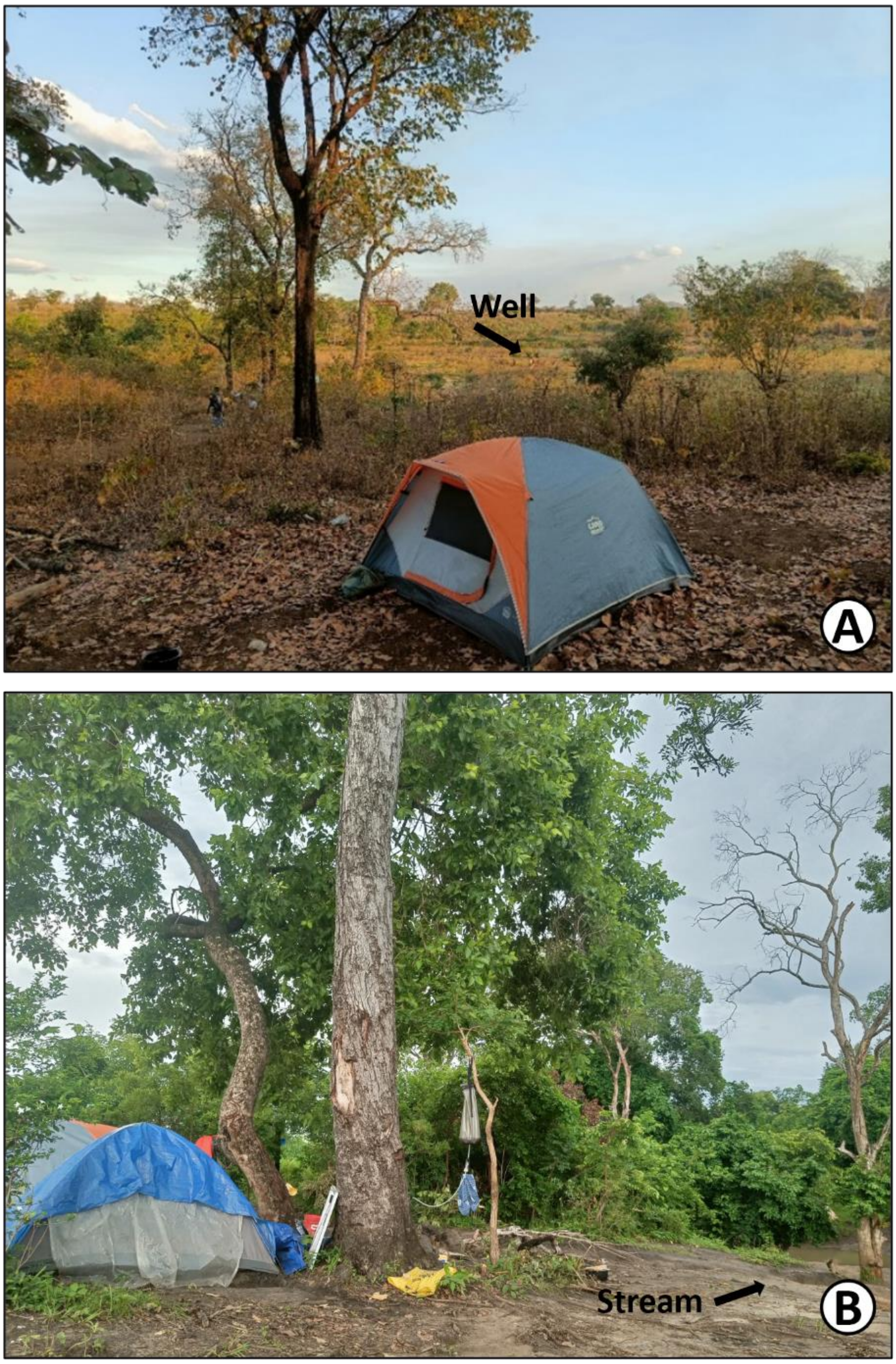


**Figure S5.5:** Camp number 18, *Mavimba Pori*, an example of miombo woodland that had been converted for agricultural land near the western border of ILUMA. The camp was set up close to a well during the dry season (**A**) and was located next to a flowing stream during the wet season (**B**), to fetch water for cooking and filtering drinking water.

The circuits were designed to manage the physical challenges of the study, and so camps were grouped into a circuit based on their geographic proximity to one another (Figure 1) and the practicality of visiting them all in a circular route that started and ended centrally at the Msakamba camp (Figure S5.2). Each circuit had a planned route that minimised walking distances between camps which ensured that the team had enough time to rest in the afternoon before proceeding with data collection in the evening. Camps were visited consecutively for two nights each and involved intensive daily adult mosquito collection procedures. Therefore, the circuit design for sequentially visiting camps in a rolling cross-sectional survey was crucial for sustaining optimal long-term data collection by limiting investigator fatigue and allowing them to take breaks of a few days in between periods of continuous field work that lasted about two weeks for a single circuit.

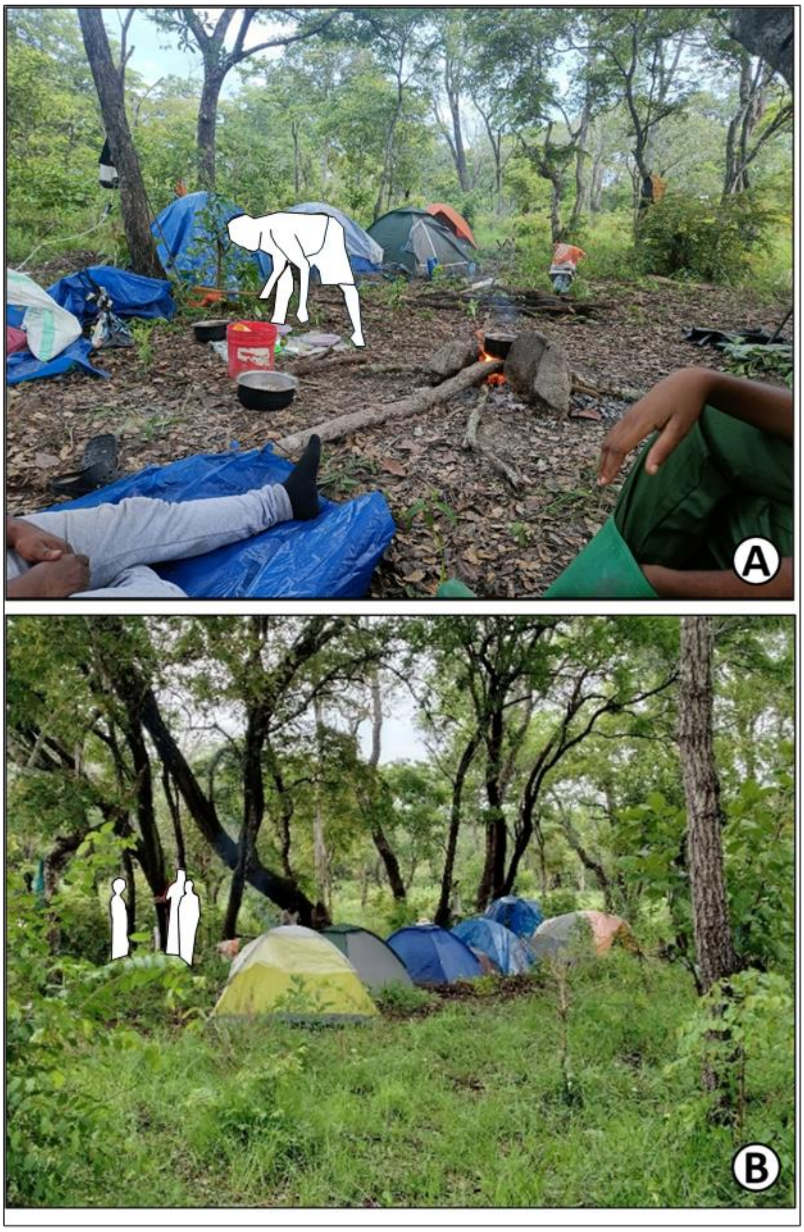


**Figure S5.6:** Camp number 11, *Kisima cha Seba*, located inside ILUMA WMA (A), and camp number 3, *Bwawa la Nyati* located inside ILUMA WMA and close to the NNP border. Both camps are examples of intact, mature miombo woodland.

The original protocol planned for a total of three rounds to be completed from January to March, March to May, and August to October 2022, representing the short rainy season, long rainy season, and the dry season, respectively. It was planned that each round of surveys in the longitudinal rolling cross-sectional study design visited all 28 camps detailed Table S5.3, but in each round a few camps were omitted for pragmatic reasons such as a lack of surface water during the dry season or inaccessibility due to severe flooding during intense rains. Also, in the first round, the north-west circuit was omitted because transport, handling and rearing procedures for the collected mosquito specimens failed, and so it was decided to adjust these procedures and start afresh with round two.

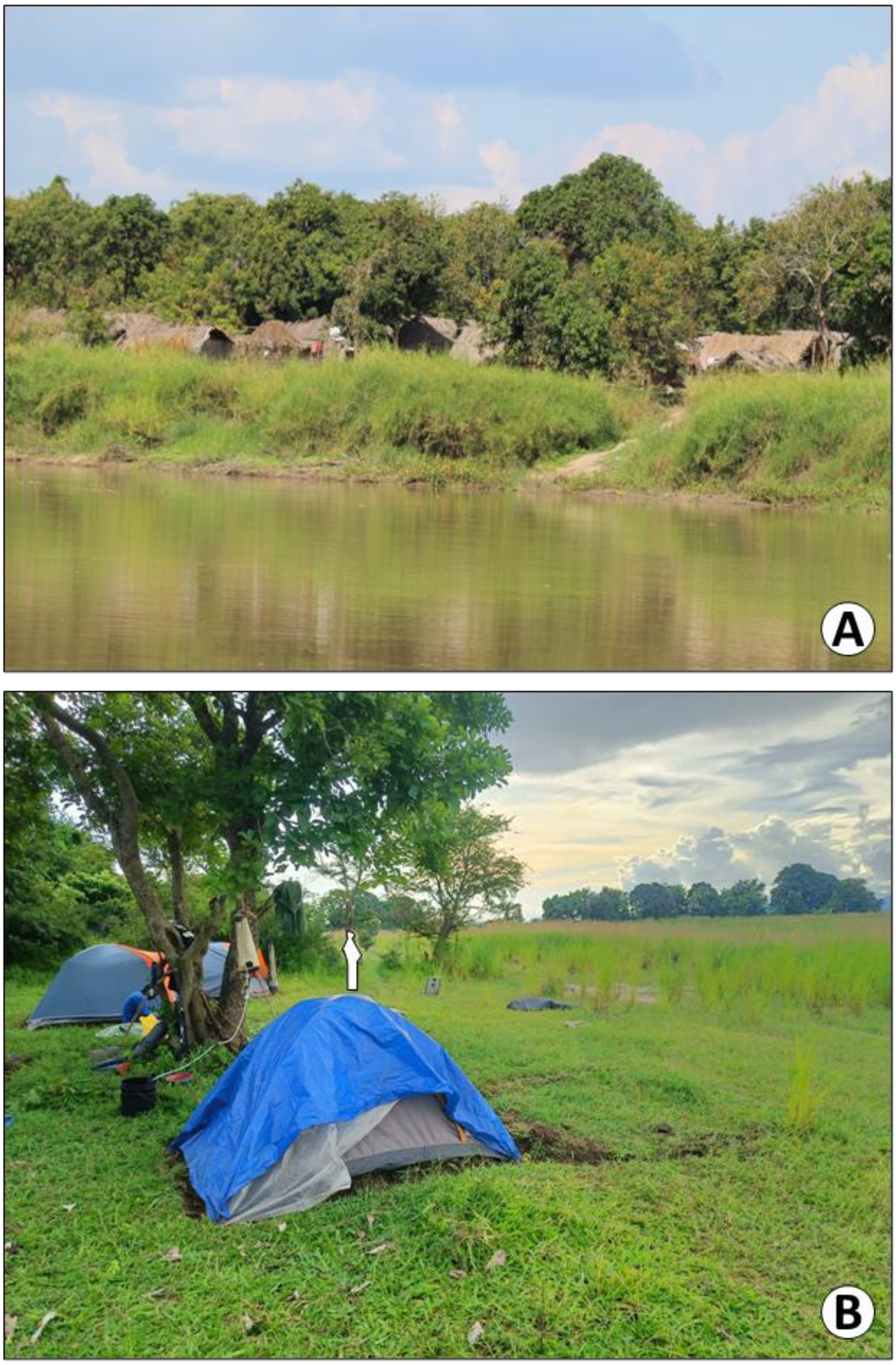

**Figure S5.7:** Examples of camp locations at the legal fishing camps inside the groundwater forest of ILUMA WMA on the south bank of the Kilombero, that practiced small-scale sustainable fishing. **A**; The biggest fishing camp inside ILUMA, Mikeregembe, where camp number 14 was located. **B**; Camp number 26, based next to the smaller fishing camp of Funga.

However, no camp inside the WMA lacked any signs of human disturbance, and only a few remained relatively conserved, so four new mobile camps inside NNP were added (numbers 29-32, Figure S5.8) to the study design at the end of round three in November 2022, forming a new circuit that was surveyed with vehicle support for logistical and safety reasons. These camps were located inside the boundary of NNP, immediately to the east of ILUMA WMA to capture the best conserved environments and were accessed by vehicle via the NNP ranger post at Boma Ulanga for which that circuit was named (Table S5.3, Figure 1). These were then repeated at the start of the fourth and final round of data collection which was completed from February to July 2023, representing the whole wet season and the beginning of the dry season for that calendar year. Considering the initial unexpected sibling species identification and their apparent association with cattle during initial data exploration (S2 Figure) and insecticide phenotype results obtained from the first four NNP camps (Kavishe DR, personal communication), it was decided to extend the sampling frame deeper into the park and adjust the field protocol to collect and immediately preserve additional collections of larvae that would not be used to rear adults in the Msakamba insectary (S3 Appendix).

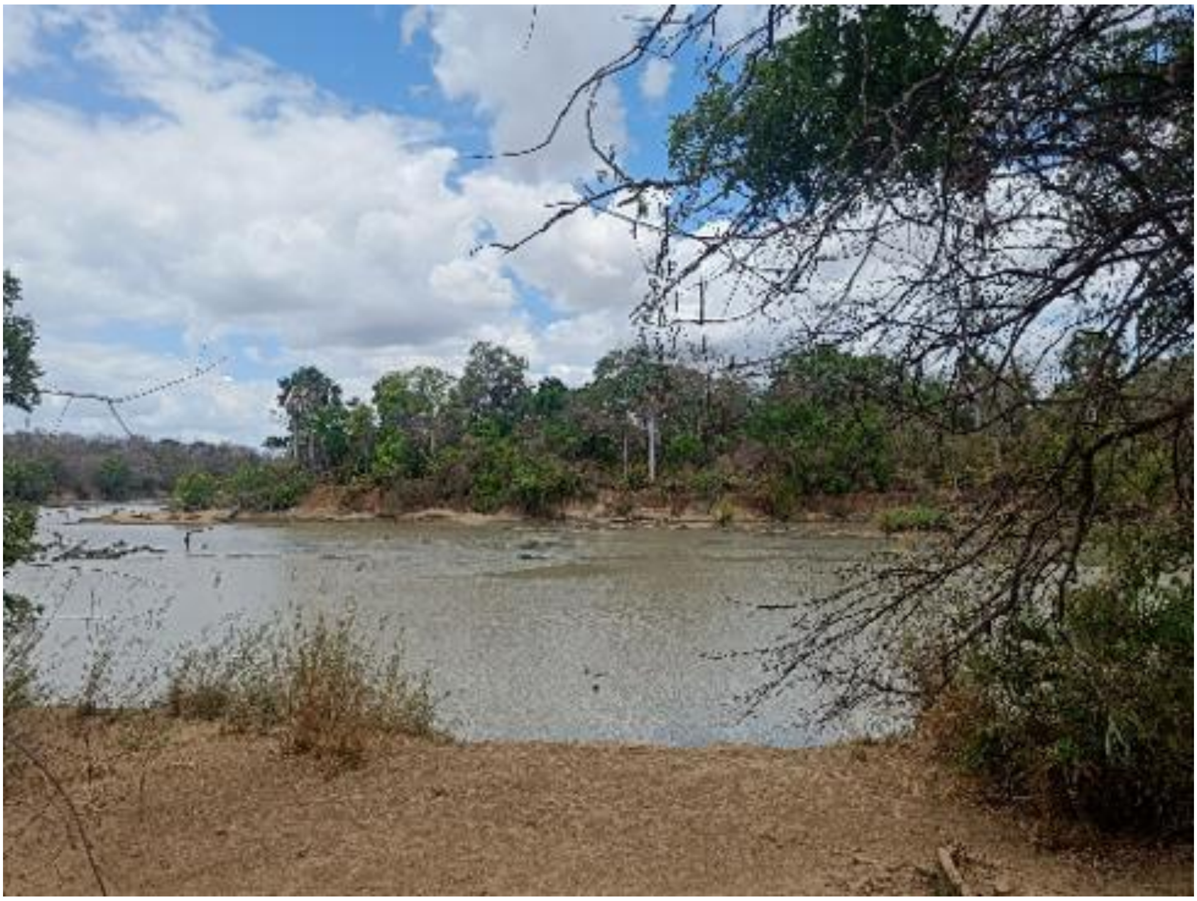

**Figure S5.8:** The view from camp number 31, *Zanzibar*, with a fully intact natural ecosystem inside NNP away from any signs of humans or livestock.

To obtain samples as far away from human beings and as deep into fully intact natural ecosystems as possible, another, eight camps inside NNP were added to the end of the fourth and final round. An additional five camps (*Kilombero circuit)* that were accessed by boat via the Kilombero river were visited (Figure 1), followed by camps 38-40 that were accessed by vehicle along main park roads via the Msolwa ranger post and were therefore named accordingly as the *Molswa circuit* (Figure 1). Camp number 38 marks the furthest possible aerial distance from any human habitation (Figure 1). The other two camps from the Msolwa circuit (39, 40) were in a north-east direction from ILUMA WMA, and were respectively located 21.9km and 41.5km from the nearest point of the NNP border. However, the latter was close to a human settlement built within the NNP, namely the construction site for the Julius Nyerere hydroelectric power plant, formerly known as Stiegler's Gorge (Figure 1). The number and name of all camp locations and the number of times they were surveyed is presented in table S5.3.

The team of three investigators responsible for collecting data and mosquito specimens were accompanied by eight VGS, whom in addition to providing food, water, shelter and security, were also integral to the implementation of the field protocol. Walking from one camp to the next was only possible under the guidance of at least two armed VGS members, whom also assisted with tracking for the surveys of humans, livestock and wild animals. Field procedures conducted inside NNP were also completed under the oversight and guidance of armed park rangers. Other VGS members assisted with carrying equipment and supplies, as well as the routine logistical and scientific activities such as setting up camp, cooking, filtering water for drinking, and collecting and maintaining mosquitoes before they were transported back to the central camp insectary at Msakamba. The field protocol was driven by the primary objective for the broader study that was to find insecticide susceptible mosquitoes, so all field activities, including the larval surveys, and the land use and mammalian activity surveys reported herein were centred around the daily collection of live adult mosquitoes.
