## Supplementary material for "Blood host preferences and competitive inter-species dynamics within an African malaria vector species complex inferred from signs of animal activity around aquatic larval habitats": S6_Appendix

### S6 appendix: Assessment of natural ecosystem integrity, historical natural land cover and distance from nearest national park boundary or human settlement

#### *Scoring locations in terms of a Subjective Natural Ecosystem Integrity Index (SNEII) based on consensus investigator impressions*

Following the initial year of data collection and prior to any analyses, a *subjective natural ecosystem integrity index* (SNEII) was devised (1), as a simple and intuitive means of assessing the ecological intactness of a location relative to its intact natural state and the level of degradation, if any, that had occurred there. This index was created by assigning each camp a score from 0% to 100%. An SNEII score of 0% represents a fully domesticated landscape, where human settlement has long been established and the ancillary activities that are associated with permanent settlements and unrestricted land use, such as land clearing, agriculture and extractive practices like charcoal burning are ubiquitous. At the other extreme end of the scale, a score of 100% represents a fully intact, natural ecosystem, with no evidence of human activity of any kind. Individual scores were carefully assigned based on a consensus among the three investigators of recollected impressions of the landcover and land use patterns. Field notes that were recorded and any photographs that were taken during the study were also considered in discussions to agree on consensus scores. An initial SNEII for camp numbers 1 to 32 was formulated in December 2022, after data collection had been completed for that calendar year. As 8 extra camps were added for the final round of data collection in 2023, the SNEII was subsequently edited to accommodate these additional 8 camps further inside NNP that were visited once. The scores were initially drafted and then finalised in Microsoft Excel® after further detailed conversations reached a conclusive consensus.

The primary and most important criterion considered by the three investigators when assigning a SNEII score was the perceived intensity of land degradation caused by the following: (a) the conversion of land for agriculture, (b) deforestation for the purpose of charcoal production, timber harvesting or human settlement (c) livestock herding. The intensity of degradation of the land surrounding a 2km radius of the camp was initially determined by estimating the proportion of remaining intact ecosystems using personal observations that were made while present in the camp and during the larval habitat surveys (S3 Appendix) and radial surveys of human, livestock, wildlife activity and land use (2) estimated proportion of intact land was then used as a reference starting point for assigning the score based on other considerations. For example, a camp with an estimated 10% of intact natural landcover was initially given an approximate score of 10 before that was considered for adjustment based on other factors as described below. Camps that evidently belonged on the extreme ends of the scale were assigned first before those that were less clear and required a more in-depth discussion. Most camps that required extra discussion were areas where intermittent patches of deforestation, small-scale agriculture and human settlement occurred in between expanses of intact woodland. These were all located inside the ILUMA WMA, in areas that were either regenerating from previous encroachment, or beginning to experience an influx of human activities.

The perceived presence of wildlife and humans as indicators of ecosystem integrity were considered to be of secondary importance when assigning scores but were nevertheless useful deliberations following the agreement of a consensus score with respect to the intensity of land degradation. Personal observations clearly indicated that species diversity generally increased in more intact natural ecosystems, but that some wild species were found more often than others in areas of degraded land. These criteria were useful when deciphering the scores of multiple camps within ILUMA WMA that had similar proportions of intact woodland and land degradation. It was also useful for areas of regenerating woodland that had been previously encroached upon and degraded by humans. Regenerated woodland with observed signs of high species diversity, influenced the investigators to assign a higher SNEII score, whereas woodlands in the early stages of regeneration that lacked obvious signs of wildlife diversity were instead adjusted downward. However, using wildlife as an indicator of ecosystem integrity often proved troublesome because it also required careful consideration of the fact that wild animal communities change naturally with land cover type, season, and even short-term weather conditions. The ability to distinguish land cover types was also a notable factor when assigning a value for this index as it was important to not mistake natural grasslands, scrublands, and valleys for deforestation.

From personal observations, it seemed to the investigators that the presence of humans was negatively associated with ecosystem integrity. However, it was not a consistently reliable criterion as some human activities including the practice of legal, sustainable fishing inside the WMA, had little perceptible impact on the surrounding natural environment. Although meat and fish poaching were not considered to have any direct effect on ecosystem integrity by the investigators, any evidence of these illegal activities prompted the investigators to assign a slightly lower final score depending on their perceived intensity. After the initial discussion was completed and the initial SNEII scores were drafted, these values were discussed again to ensure that the order and distribution of the final SNEII scores were assigned as well as possible regarding the above criteria. The overall outcomes of this subjective ecosystem scoring processes are detailed in table S6.1 and narrated as follows.

**Table S6.1:** The values and corresponding ranks for the subjective natural ecosystem integrity index (SNEII) based on a consensus of recollected impressions and the objective natural ecosystem integrity index (ONEII) based on principal component analysis (PCA) of natural resource use, direct sightings and signs of tracks, spoor and other signs for wildlife, livestock and humans, together with every recorded estimate of the proportion of land used for rice farming and other tillage crops.

| | | | Subjective Ecosystem Integrity Index | | Objective Ecosystem Integrity Index | |
|---|---|---|---|---|---|---|
| Camp | Camp Name | Location | Value [a] | Rank [b] | Value [c] | Rank [d] |
| 1 | Msakamba | Inside ILUMA | 70% | 16.5 | -0.301 | 17 |
| 2 | Bwawa la Msiba wa Deo | Inside ILUMA | 84% | 24.0 | -0.361 | 28 |
| 3 | Bwawa la Nyati | Inside ILUMA | 85% | 25.5 | -0.323 | 22 |
| 4 | Bwawa la Nandete | Inside ILUMA | 35% | 10.0 | -0.283 | 14 |
| 5 | Korongo la Bundu | Inside ILUMA | 30% | 9.0 | -0.279 | 13 |
| 6 | Bwawa la Namamba | Inside ILUMA | 40% | 11.0 | -0.266 | 12 |
| 7 | Bwawa la Chakacheni | Inside ILUMA | 15% | 7.0 | -0.255 | 10 |
| 8 | Bwawa la Njuju | Inside ILUMA | 81% | 21.0 | -0.325 | 24 |
| 9 | Bwawa la Chamvi | Inside ILUMA | 86% | 27.0 | -0.353 | 26 |
| 10 | Bwawa la Miembeni | Inside ILUMA | 68% | 15.0 | -0.397 | 30 |
| 11 | Kisima cha Seba | Inside ILUMA | 82% | 22.0 | -0.293 | 16 |
| 12 | Bwawa la Maya | Inside ILUMA | 87% | 28.0 | -0.359 | 27 |
| 13 | Bwawa la Mrope | Inside ILUMA | 90% | 29.0 | -0.325 | 23 |
| 14 | Mikeregembe | Inside ILUMA | 75% | 19.0 | -0.311 | 20 |
| 15 | Mdalangwila | Inside ILUMA | 72% | 18.0 | -0.310 | 19 |
| 16 | Bwawa la Mnyuamachi | Outside ILUMA | 6% | 5.0 | 1.185 | 5 |
| 17 | Tuliza Moyo | Outside ILUMA | 4% | 3.0 | 1.219 | 4 |
| 18 | Mavimba Pori | Outside ILUMA | 10% | 6.0 | 0.158 | 7 |

| | | | | | | |
|---|---|---|---|---|---|---|
| 19 | Bwawa la Selesusi | Inside ILUMA | 25% | 8.0 | -0.253 | 9 |
| 20 | Makingi | Outside ILUMA | 0% | 1.0 | 1.286 | 3 |
| 21 | Bwawa la Mpunga | Inside ILUMA | 55% | 14.0 | -0.263 | 11 |
| 22 | Kisaki | Outside ILUMA | 5% | 4.0 | 4.722 | 1 |
| 23 | Uwanja wa Ndege | Outside ILUMA | 3% | 2.0 | 2.750 | 2 |
| 24 | Bwawa la Mkwajuni | Inside ILUMA | 45% | 12.0 | -0.138 | 8 |
| 25 | Bwawa la Mamba Luhogi | Inside ILUMA | 48% | 13.0 | 0.391 | 6 |
| 26 | Funga | Inside ILUMA | 77% | 20.0 | -0.287 | 15 |
| 27 | Bwawa la Mlenda | Inside ILUMA | 83% | 23.0 | -0.306 | 18 |
| 28 | Bwawa la Semka | Inside ILUMA | 85% | 25.5 | -0.382 | 29 |
| 29 | Kambi ya Simba | Inside NNP | 97% | 32.0 | -0.478 | 36 |
| 30 | Bwawa la Kiboko Zanzibar | Inside NNP | 100% | 37.5 | -0.520 | 40 |
| 31 | Zanzibar | Inside NNP | 95% | 30.5 | -0.458 | 35 |
| 32 | Bwawa la Moto | Inside NNP | 100% | 37.5 | -0.513 | 39 |
| 33 | Kambi ya Mamba | Inside NNP | 98% | 33.0 | -0.415 | 32 |
| 34 | Kambi ya Machuma | Inside NNP | 100% | 37.5 | -0.500 | 37 |
| 35 | Serengeti Ndogo | Inside NNP | 95% | 30.5 | -0.451 | 34 |
| 36 | Kambi ya Makutano | Inside NNP | 100% | 37.5 | -0.446 | 33 |
| 37 | Kambi ya Mawe | Inside NNP | 100% | 37.5 | -0.504 | 38 |
| 38 | Shughuli kubwa | Inside NNP | 100% | 37.5 | -0.339 | 25 |
| 39 | Bwawa la Chatu | Inside NNP | 99% | 34.0 | -0.399 | 31 |
| 40 | Bwawa la Umeme | Inside NNP | 70% | 16.50 | -0.315 | 21 |

[a] The subjective natural ecosystem integrity score.
[b] The rank of each camp based on the subjective ecosystem integrity scores, where the lowest rank represents fully degraded, domesticated land and the highest rank represents an absolute intact natural ecosystem.
[c] PC1 values that were derived from a PCA accounting for all recorded detections of humans, livestock and wildlife, and land use activities that were standardised using z-scores.
[d] The rank of each camp based on standardised PC1 values, where the lowest rank represents the most degraded camp and the highest rank represents the best conserved camp.

All final scores of less than or equal to 10% were given to camps that were based at permanent human settlements outside ILUMA WMA (Table S6.1) and surrounded by domesticated land which was primarily used for rice farming and other tillage agriculture. Camps that were given SNEII scores of 15%, 25%, 30%, 35%, 45%, 48%, respectively, were all located inside the ILUMA boundary (Table S6.1) but were heavily encroached and included large areas of degraded land for agriculture and human settlement. A decrease in wildlife diversity was also seen around these camps based on recollected visual observations. Only two camps were assigned values between 50% and 70%, both of which were located inside ILUMA WMA (Table S6.1). *Bwawa la Mpunga* (camp number 21), was assigned a score of 55% and was based at a waterhole and surrounding swamp that had been exploited for rice farming. Despite this, the surrounding woodland was not as heavily degraded as other locations on the western boundary of ILUMA WMA. Similarly, *Bwawa la Miembeni* (camp number 10), was located in a large valley that was also exploited for rice farming. However, the surrounding woodland was fully intact and therefore received a higher score compared to *Bwawa la Mpunga* (Table S6.1). Although the location of the main camp *Msakamba* was surrounded by signs of land degradation and deforestation, it was assigned a score of 70% as progressive regeneration of the woodland and increased presence of wildlife was visually observed during the study period (Table S6.1).

All other camps with higher scores between 70% and 90% were in the groundwater forest in the north of ILUMA WMA or in the miombo woodlands that were located closer to the NNP border. Three small, legal fishing camps in the north of ILUMA WMA, on the southern bank of the Kilombero River (camp numbers 14,15 and 26), as seen in figure 1, were assigned high SNEII scores of 75%, 72%, and 77%, respectively, despite being human settlements with permanent structures. This is because small-scale sustainable fishing and very limited extractive activities are permitted in these fishing camps but are carefully monitored. Also, the vigilance of the residents, who often actively support the conservation activities of the ILUMA WMA, acts as a deterrent for unauthorised human activity in the areas surrounding their settlement (3). Correspondingly, the forests and wetlands that surround these authorised human settlements remain fully intact (S5 Appendix, figure S5.7). Two camps inside NNP were given a score of 95% as they had still the remnants of old tourist camps. Six camps in NNP had the highest possible score of 100%, in which habitats were absolutely intact and did not have a single observed sign of poaching activity.

The lowest score inside NNP was assigned to one camp located 42km inside the park boundary. A score of 70% was given to camp number 40, *Bwawa la Umeme* because the area around this camp was recently cleared in relation to the Julius Nyerere Hydropower Station located on the Rufiji River (Figure 1). Despite this evidence of human intervention, large proportions of the area around the camp were in good condition and personal observations indicated that wildlife was abundant and diverse, albeit less so than in any other NNP camp.

***Validation of the SNEII by comparison an Objective Natural Ecosystem Integrity Index (ONEII) based on statistical synthesis of formal quantitative surveys of human, livestock and wildlife activities, as well as land cover attributes***

To assess the validity of this intuitive approach to quantifying ecosystem integrity, this subjective index was tested for correlation with an alternative, *objective natural ecosystem integrity index* (ONEII), which used a far more laborious approach based on statistical syntheses of the data from the radial surveys of human, livestock and wildlife activity and land use (S8 Appendix). A principal component analysis (PCA) of every recorded detection of natural resource use, direct sightings and signs of tracks, spoor and other signs for wildlife, livestock and humans, together with every recorded estimate of the proportion of land used for rice farming and other tillage crops (S8 Appendix) was conducted using the *prcomp* function in open software R version 4.3.1. Given that most of the variance was attributed to the first principal component (PC1), estimates of this PC1 for each camp were derived and then normalised by z-scores as described in Duggan *et al.* (1, 3). As the PCA was based on recorded detections collected within a 2km radius of the camp and each camp was located 4-15km apart, the PC1 scores for each camp location was already assumed to be reasonably spatially independent. The direction of the scaled PC1 values based on all the recorded detections across all the completed radial surveys were interpreted intuitively with the most positive and most negative values for total detections clearly being associated with the most fully domesticated and fully intact, natural ecosystems, respectively (Table S6.1). The PC1 based indicators of ecosystem integrity were then ranked so that the lowest ranked camp represented a fully domesticated ecosystem with a permanent human settlement, while the highest rank camp represented a very well conserved ecosystem with an intact natural wildlife community in the complete absence of humans or livestock (Table S6.1).

A Spearman's rank correlation test using the *cor.test* function in R version 4.3.1, demonstrated a strong positive correlation between the ranked SNEII scores and the ranked standardised PC1 indicators of ecosystem integrity (Figure S6.2). Like the SNEII, the PCA-based ONEII also ranked the camps outside ILUMA WMA as the most degraded. However, for the ONEII, one encroached camp inside ILUMA WMA, *Bwawa la Mamba Luhogi* (camp number 25), was ranked lower than a camp outside the protection of ILUMA WMA (*Mavimba Pori,* camp number 18) (Table S6.1). This highlights the severity of encroachment inside ILUMA WMA and its impact on natural ecosystems. Rice farming and other tillage farming were all evident in low scoring camps outside ILUMA WMA and in the encroached areas inside ILUMA WMA, and was the predominant land use in the three lowest scoring camps, suggesting that agriculture is a key determinant of poor ecosystem integrity. The ONEII ranked the top ten highest ranked camps inside NNP.

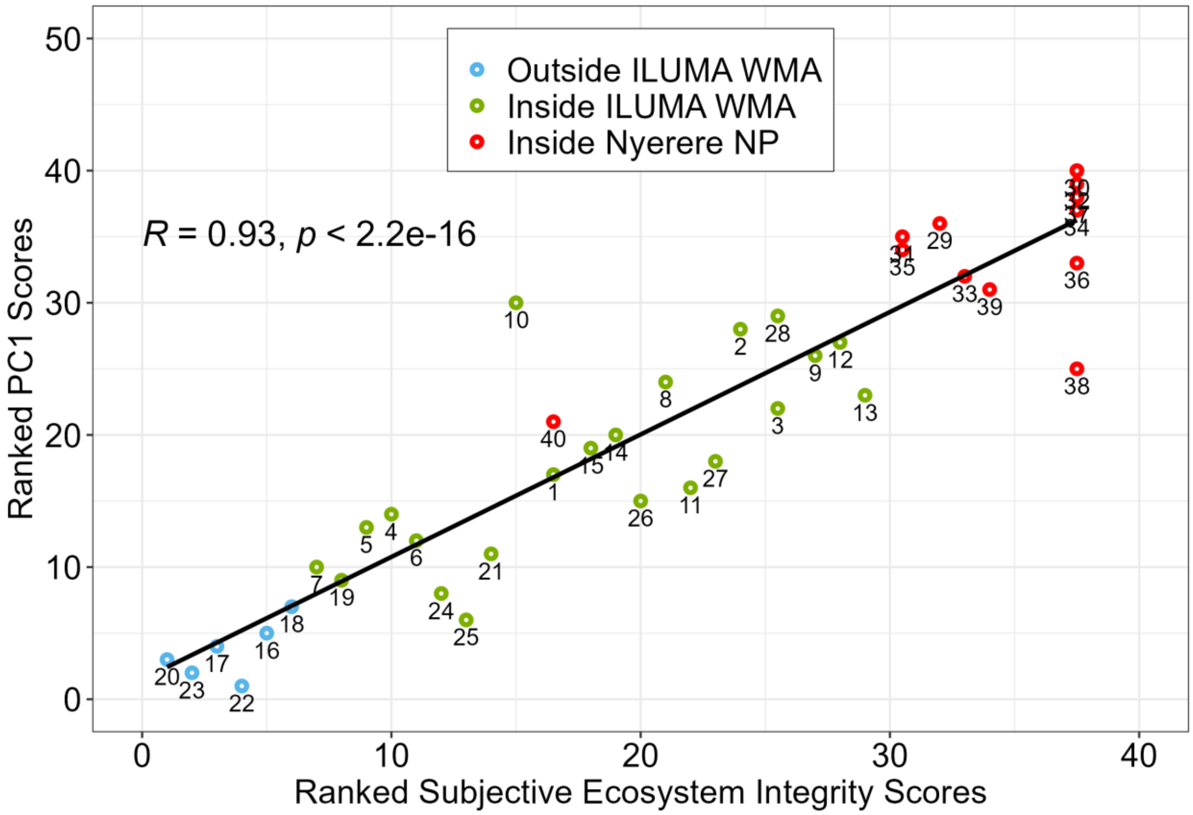


**Figure S6.2.** The ranked standardised PC1 values of the ONEII for each camp number plotted against the ranked SNEII scores for each camp. Note that while this graph is almost identical to that in Duggan *et al.* (1) it includes 8 extra camps, one of which (number 40), is a useful exception with unusually low ecosystem integrity for a location so far inside NNP which helps to prove the general rule of close correlation.

Apart from two moderate outliers, *Shuguli kubwa* (camp number 36) and *Bwawa la Miembeni* (camp number 10), that were respectively ranked lower and higher compared to SNEII ranks, the high correlation displayed in figure S6.2 indicates that this novel methodology of formulating an ecosystem integrity index based on repeated recollective impressions of the study sites was a highly effective approach, and even has advantages compared to the use of the ONEII. The more convenient and intuitive SNEII is measured on a scale from 0% to 100%, thus providing a simplified interpretation of the ecological state at each study site where larvae surveys were conducted. Furthermore, multivariate regression analysis demonstrated that the ONEII was far less sensitive than the SNEII to the observed occurrence and intensity of various human activities (1). Throughout this study, the SNEII was therefore consistently used as the sole synthetic indicator of natural ecosystem integrity in the analyses of larval occupancy (S10 Appendix) and the association between environmental parameters and *An. gambiae* complex sibling species composition (S10 Appendix)

***Assigning values for historical natural land cover and distance to nearest national park boundary nearest human settlement***

Before performing the statistical analyses on the occupancy data and sibling species composition data, two additional variables were added to the dataset to take further spatial and environmental factors of each camp location into account. First, a continuous variable *distance* was created to account for any potential variance attributable to the natural dispersal of mosquitoes (4), from suitable environments to unsuitable ones, which might directly impact occupancy rates or the species composition of populations (S9 Appendix). The values for each camp were calculated by measuring the number of kilometres (km) from the camp coordinates to the nearest point on the NNP park boundary line (Figure 1), using the geographic information system (GIS) open-source software, QGIS Version 3.30.2. Camps that were located outside the NNP boundary were assigned a negative value for the distance in km, while camps that were located inside the NNP boundary were assigned a positive value. Camp number 40, *Bwawa la Umeme,* which already proved to be an exception of the SNEII because of the observations of deforestation and human activity inside the national park, also required consideration with respect to the distance variable. Despite being one of the furthest camps, located approximately 40km from the nearest point on the NNP boundary (Figure 1), it was also located just 16km from the Julius Nyerere Hydropower plant (Figure 1), where there is also a large permanent human settlement resident to those working within the NNP, which may provide a more suitable environment for mosquito populations. Therefore, this camp was again considered to be an exception, and so to improve model fits during statistical analyses, the value in km for this camp only was adjusted to the distance from this known permanent human settlement inside the park rather than the distance to the nearest NNP boundary.

The categorical variable named *historical landcover* was also created to account for any potential environmental variability associated with different natural landcover types that may influence occupancy and species composition (further discussed S9 Appendix). There were three distinguished historical landcover types found across the study area; evergreen groundwater forest, moist miombo woodland savanna, and dry acacia savanna. The predominant landcover type was assigned to each camp and was based on investigator recollections, the historical knowledge of VGS regarding fully domesticated areas, and satellite imagery available through Google Earth®. The landcover type was easily distinguished using satellite imagery as the density of vegetation for each of these landcover types vary significantly. The perennial groundwater forest in the north of ILUMA WMA was much denser with a predominantly closed canopy cover that was dark green in colour compared to the deciduous miombo woodlands that had much more broken canopy cover. Camps located in dry acacia savanna were identified as having acacia trees and scrub with only patchy canopy cover, which can easily be distinguished by an expansive landscape of sparse vegetation and light-coloured soil. Camps that were located in the villages outside the western ILUMA WMA border were all assigned miombo woodland, as this was known to be their historical predominant landcover type according to the VGS who would have been resident there.

Once each camp was assigned a SNEII score, a value for the distance from the camp to the nearest NNP boundary or human settlement inside NNP, and the historical landcover type, the datasets were prepared for final analysis as outlined in S10 Appendix.
