## Supplementary material for "Blood host preferences and competitive inter-species dynamics within an African malaria vector species complex inferred from signs of animal activity around aquatic larval habitats": S7_Figure

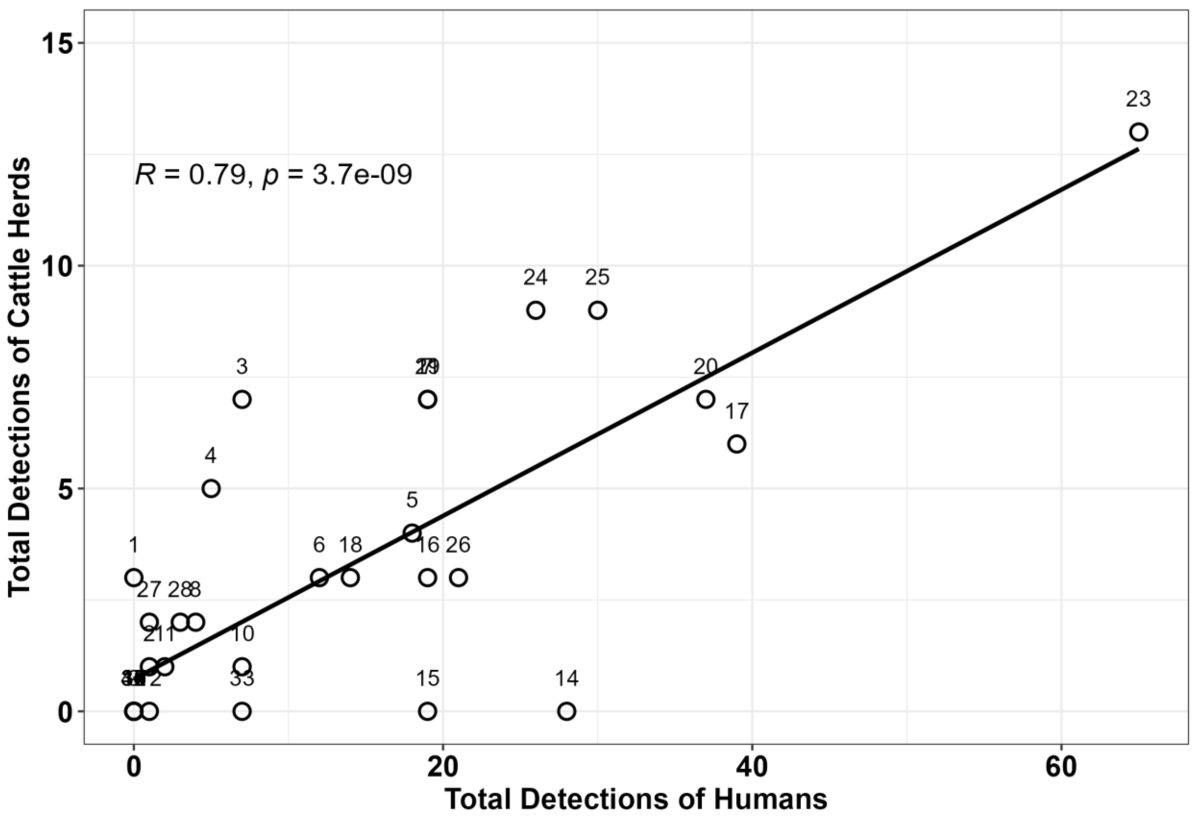


**S7 Figure.** The detection frequency of humans plotted against the detection frequency of cattle herds at each camp number demonstrating a strong linear correlation, as tested using a Pearson's linear correlation test.
