## Supplementary material for "Blood host preferences and competitive inter-species dynamics within an African malaria vector species complex inferred from signs of animal activity around aquatic larval habitats": S8_Appendix

### S8 Appendix: Procedures for surveying activities of humans, livestock and wildlife, as well as land use and vegetation cover, around mosquito larval habitats within a 2km radius of each camp location

All direct visual observations and indirect signs of activity by humans, livestock or wild mammals (Figures S8.1 to S8.4) encountered along the routes taken between and around the fringes of water bodies were recorded using the tools provided in Tables S8.5 to S8.8). This system for categorical classification of observed indirect signs of wild animals (Table S8.5), livestock and humans included approximate age classifications agreed with the VGS on the basis that they could distinguish them with reasonable reliability (Table S8.6). All data were recorded on a standardized form (Table S8.7) using a laminated hard copy of a detailed classification key (Table S8.8).

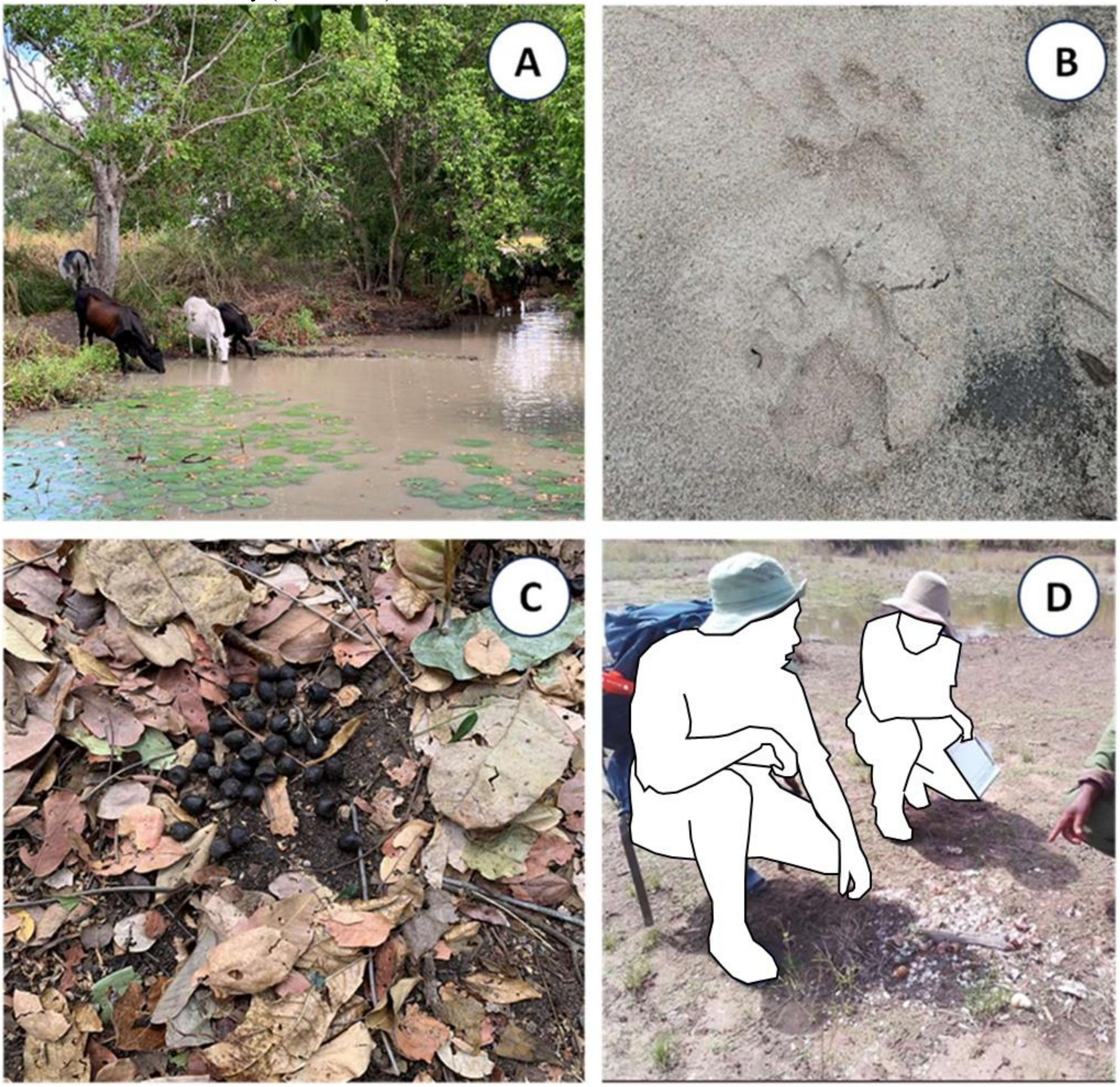

**Figure S8.1:** Photographs illustrating examples of different ways in which signs of livestock, humans or wild mammals were detected and recorded as either direct sighting, tracks, spoor or other signs. Panel **A** illustrates a typical direct sighting of cattle. Panel **B** illustrates an example of a rare observation, namely fresh lion prints on damp sand that resulted in perfectly clear footprints and an unambiguous identification of the animal and the age of its tracks. In contrast, panel **C** illustrates a typical example of one of the most frequently recorded detections throughout the entire survey-Hartebeest spoor, in this case approximately three days old. Panel **D** illustrates a particularly common example of “other signs”, specifically broken shells and a stick used as an anvil by water mongoose to crack open snails and shellfish.

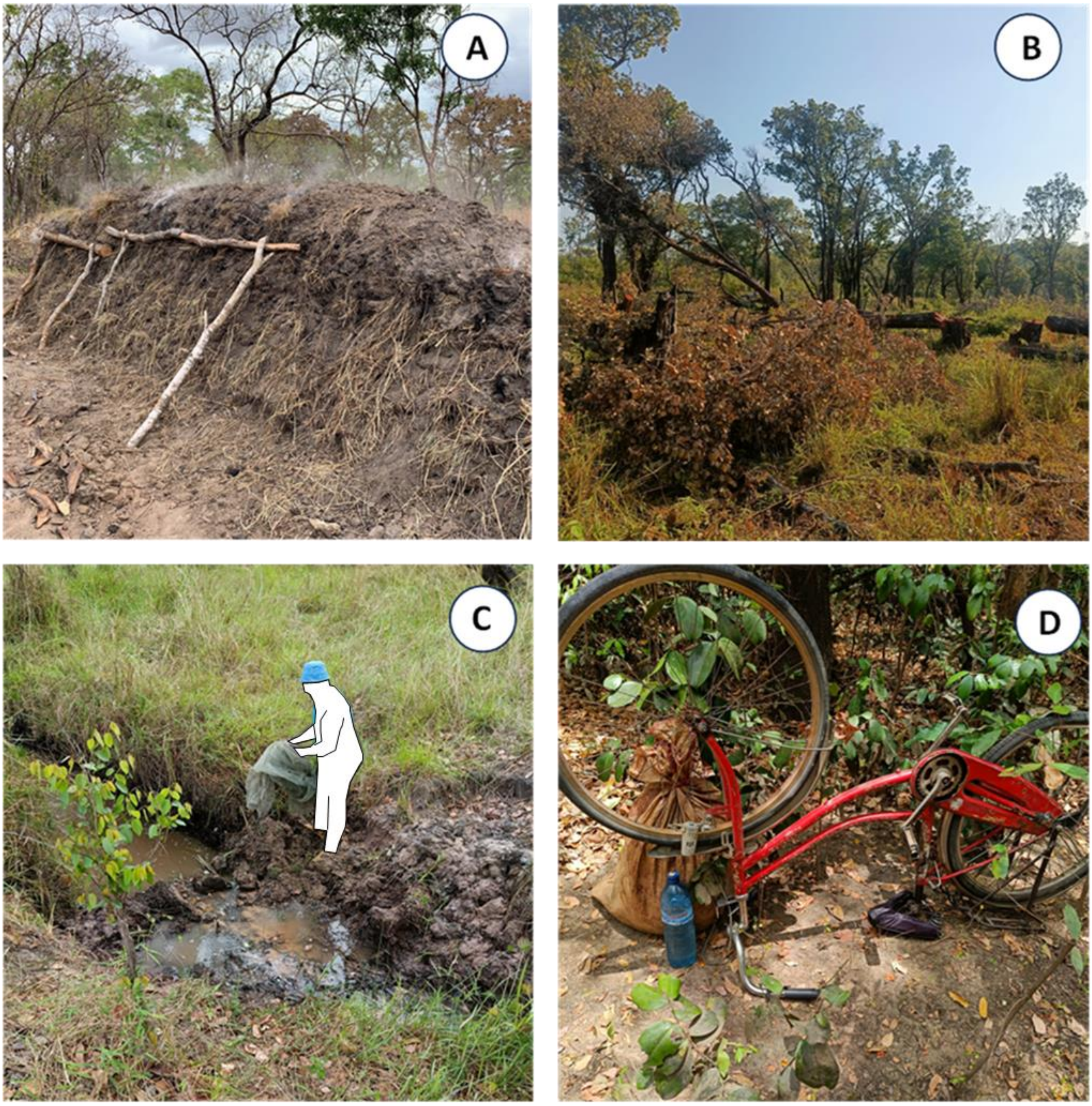

**Figure S8.2:** Photographs illustrating unauthorised activities recorded during data collection in the field. Panel **A** shows active charcoal burning. Timber from felled trees is placed in a pile, covered in soil to create a low oxygen environment, and then set on fire. Panel **B** displays an area that is undergoing active deforestation. The trees that have been felled would be used for charcoal production or for the construction of human settlements or occasionally timber to be sold commercially. Panel **C** shows evidence of unauthorised fishing, in which a dam made of soil and mud was constructed across a large stream to block the flow of water and prevent fish from moving past that point. Also exhibited is a bed net, which had been used to catch fish trapped within the dammed pond. Panel **D** exhibits items seized from a meat poacher during a routine patrol. Bicycles are an efficient mode of transport through the WMA for both people and goods. Sufficient supplies for up to a week and equipment used to capture wild animals were in the sack behind the bicycle.

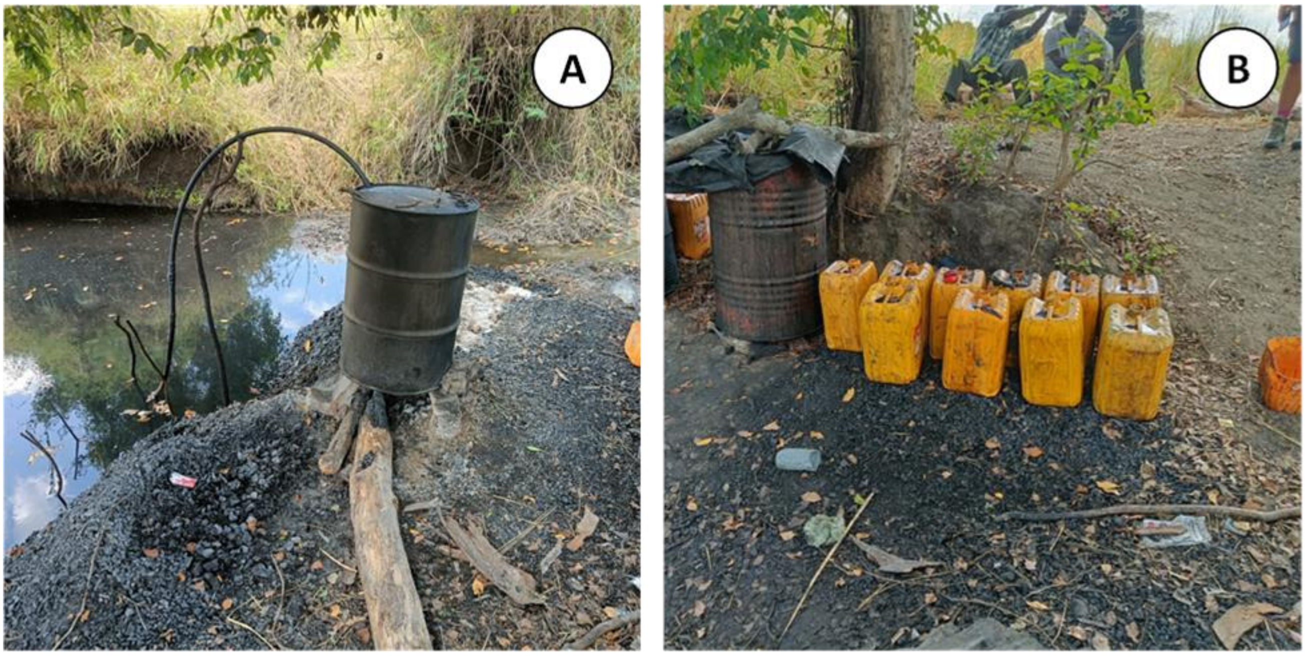

**Figure S8.3:** Panels **A** and **B** both illustrate an unauthorised human activity taking place in the ILUMA WMA that was not anticipated a priori, which is presented an example of an unusual activity. Here, a form of homemade alcohol known as *gongo* is being produced through distillation with grain cultivated illegally inside the conservation area. As with fish poaching, a river is blocked which causes knock-on issues for the aquatic ecosystem. Pollution of the watercourse is evident here.

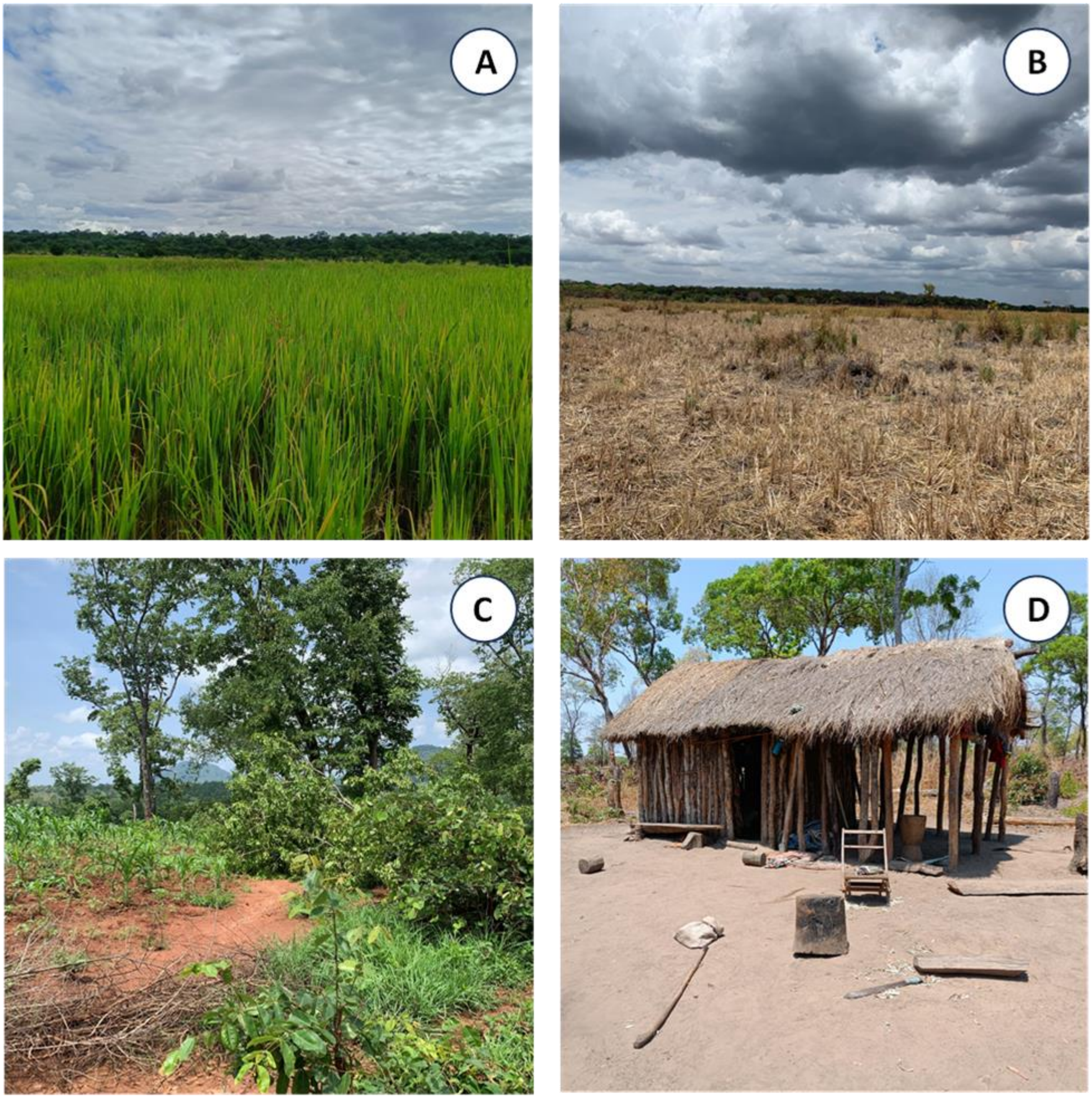

**Figure S8.4:** Images showing how the impression of land cover for rice farming and other tillage agriculture was recorded. Both panels **A** and **B** display rice agriculture in different stages. The rice in image **A** is still growing and is lush green. Image **B** depicts how this same area looked once the rice had been harvested and the dry season began. Panel **C** demonstrates the other forms of tillage farming detected in ILUMA. Here, maize is beginning to grow in deforested and cleared land. The mature trees in the background depict what the area looked prior to agricultural use. Panel **D** illustrates an unauthorised human settlement inside ILUMA structures like this usually accompany agricultural activity such as that showed in image **C**. Note here the level of land clearance in the area surrounding the settlement.

**Table S8.5:** Complete list of all wild herbivores, wild carnivores, wild primates and prosimians and wild rodents sampled in this study.

| **Wild Herbivores.** | **Wild Carnivores.** | **Wild Primates and Prosimians.** |
|---|---|---|
| Bohor Reedbuck (*Redunca redunca wardi*) | Lion (*Panthera leo*) | Lesser Bush Baby (*Galago senegalensis*) |
| Common Waterbuck (*Kobus ellipsiprymnus ellipsiprymnus*) | Leopard (*Panthera pardus*) | Greater Bush Baby (*Galago crassicaudatus*) |
| Puku (*Kobus vardoni*) | African Wild Cat (*Felis libyca*) | Yellow Baboon (*Papio cynocephalus cynocephalus*) |
| Bushbuck (*Tragelaphus scriptus*) | African Civet (*Civettictis civetta*) | Vervet Monkey (*Cercopithecus aethiops*) |
| Common Eland (*Taurotragus oryx*) | Genet (*Genetta spp.*) | Blue Monkey (*Cercopithecus mitis*) |
| Greater Kudu (*Tragelaphus strepsiceros*) | Spotted Hyena (*Crocuta Crocuta*) | |
| Hartebeest (*Alcelaphus buselaphus*) | African Wild Dog (*Lycaon pictus*) | **Wild Rodents.** |
| Wildebeest (*Coonochaetes taurinus*) | Side-Striped Jackal (*Canis adustus*) | Porcupine (*Hystrix cristata*) |
| Sable (*Hippotragus niger*) | Slender Mongoose (*Herpestes sanguineus*) | Giant Cane Rat (*Thryonomys swinderainus*) |
| Warthog (*Phacochoerus africanus*) | Banded Mongoose (*Mungos mungo*) | |
| Bushpig (*Potamochoerus porcus*) | Water Mongoose (*Atilax paludinosus*) | **Macroscelidea.** |
| Hippopotamus (*Hippopotamus amphibius*) | White-Tailed Mongoose (*Ichneumia albicauda*) | Sengi (*Petrodromus tetradactylus*) |
| Elephant (*Loxodonta Africana*) | Honey Badger (*Mellivora capensis*) | |
| African Buffalo (*S8ncerus caffer*) | African Clawless Otter (*Aonyx capensis*) | **Tubulidentata.** |
| Plains Zebra (*Equus quagga*) | | Aardvark (*Orycteropus afer*) |
| Suni (*Neotragus moschatus*) | | |
| Common Duiker (*S8lvicapra grimmia*) | | |

| |
|---|
| Natal Red Duiker (*Cephalophus natalensis*) |
| Impala (*Aepyceros melampus*) |
| Ugogo Dikdik (*Madoqua kirkii thomasi*) |
| Sharpe's Grysbok (*Raphicerus sharpei*) |

**Table S8.6:** Criteria defining terms used in data dictionary classification key.

| |
|---|
| **Direct sighting:** Live animal or human actually seen by one or more members of the field team. |
| **Tracks:** Prints (animal or human) seen by the data recorder on the ground on the immediate path being followed. |
| **Spoor:** Faecal excrement seen by the data recorder on the on the immediate path being followed. |
| **Other signs:** Any sign of animal activity that undoubtedly links to a specific species (elephant pulling bark from a tree or water mongoose cracking shells for example). |
| **Active:** Activity was directly observed or estimated to have occurred within the last 24 hours. |
| **Recent:** Activity estimated to have occurred between 2 and 5 days ago. |
| **Old:** Activity estimated to have occurred between 6 days and 6 months ago. |

**Table S8.7:** Data collection sheet used for all radial surveys of mammalian activity, land use and vegetation cover over the course of the study.

| **Form Type (FT):** Land Use Observations (LUO) | | | | **Date (DT):** / / | **GPS Identifier (GID):** | **Form Serial Number (SEN):** |
|---|---|---|---|---|---|---|
| **Transect Start Camp Number (TSCNO):** | | | | **Transect Start Camp Name or Description (TSCND):** | | |
| **Transect Segment Number (TSN):** | | | | | **Transect Start GPS Way Point Number (TSGWPN):** | |
| **Transect Segment Page Number (TSPN):** | | | | | **Transect End GPS Way Point Number (TEGWPN):** | |
| **Form Row (FR)** | **Entity or Activity Observed (EAO)** | **Observation Category (OC)** | **Additional observation attributes (AOA)** | **Observation count or class (OCC)** | **Observation Notes (ONO):** Specify details that add qualitative insight or enable post-hoc classification | |
| 1 | | | | | | |
| 2 | | | | | | |
| 3 | | | | | | |
| 4 | | | | | | |
| 5 | | | | | | |
| 6 | | | | | | |
| 7 | | | | | | |
| 8 | | | | | | |
| 9 | | | | | | |
| 10 | | | | | | |
| 11 | | | | | | |
| 12 | | | | | | |
| 13 | | | | | | |
| 14 | | | | | | |
| 15 | | | | | | |
| 16 | | | | | | |
| 17 | | | | | | |
| 18 | | | | | | |
| 19 | | | | | | |
| 20 | | | | | | |
| 21 | | | | | | |
| 22 | | | | | | |
| 23 | | | | | | |

**Table S8.8:** Data collection key devised during preliminary study in October and November 2021 and used throughout the course of the field study to inform accurate and consistent recording of various humans, livestock and animals, their activities and land cover attributes.

| **Entity or Activity Observed (EAO)** | **Observation Category (OC)** | **Additional Observation Attributes (AOA)** | **Observation Count or Class (OCC)** |
|---|---|---|---|
| **1X** "As you go" instances of specific human land and/or natural resources usages. | | | |
| **11**: Livestock Herding | | | **1**: Active, **2**: Recent, **3**: Old, **9**: Not determined or recorded |
| **12**: Charcoal Burning | | | **1**: Active, **2**: Recent, **3**: Old, **9**: Not determined or recorded |
| **13**: Timber Harvesting | | | **1**: Active, **2**: Recent, **3**: Old, **9**: Not determined or recorded |
| **14**: Fishing | | | **1**: Active, **2**: Recent, **3**: Old, **9**: Not determined or recorded |
| **15**: Hunting | | | **1**: Active, **2**: Recent, 3: Old, **9**: Not determined or recorded |
| **16**: Human Settlement | | **1,2,3…10** structures [n], 11-20 structures **[20],** 21-50 structures **[50],** >50 structures **[100]**, not determined or recorded **[9]** | **1**: Active, **2**: Recent, **3**: Old, **9**: Not determined or recorded |
| **17**: Water Body | | | **1**: Stagnant waterbody, **2**: Puddle(s) outside a seasonal streambed, **3:** Puddle(s) outside a seasonal streambed **4**: Flowing stream, **5**: Flooded valley |
| **18:** Human dug well | | | **1**: Active, **2**: Recent, **3**: Old, **9**: Not determined or recorded |
| **19:** Meat Poaching | | | **1:** Active**, 2:** Recent, **3:** Old**, 9:** Not determined or recorded |
| **2X**: "As you go" sightings, tracks and signs of humans and animals. | | | |
| **21**: Humans | 1: Direct sighting, 2: Tracks, 3: Spoor, **4**: Other signs (Explain in notes) | **1**: ILUMA personnel, **2**: NNP Rangers, **3**: Hunting company staff, **4**: Other authorized personnel from ILUMA and other authorized institutions, **5**: Community legal presence inside conservation area, **6**: Community legal presence outside the conservation area, **7**: Charcoal transport, **8**: Timber harvesting, **9**: Timber transport, **10**: Fish harvesting, **11**: Fish transport, **12**: Community illegal presence inside the conservation area, **13:** Charcoal burner. | Distance (m) from track for direct observations, or age for tracks and signs: **1**: Active, **2**: Recent, **3**: Old, **9**: Not determined or recorded |
| **22**: Livestock and work animals | **1**: Direct sighting, **2**: Tracks, **3**: Spoor, **4**: Other | **1**: Cattle, **2**: Goats, **3**: Sheep, **4**: Rooster, **5**: Chicken, **6**: Dog | Distance (m) from track for direct observations, or age for tracks and signs: **1**: Active, **2**: Recent, **3:** Old, **9**: Not determined or recorded |
| **23**: Wild Herbivores | **1**: Direct sighting, **2**: Tracks, **3**: Spoor, **4**: Other signs (Explain in notes) | 1X: Reduncini (**11**: Reedbuck, **12**: Waterbuck, **13**: Puku), 2X: Spiral Horned Antelopes (**21**: Bushbuck, **22**: Eland, **23**: Greater Kudu) 3X: Alcephalines (**31**: Hartebeest, **32**: Wildebeest), 4X: Hippotragini (**41**: Sable Antelope), 5X: Suidae (**51**: Warthog, **52**: Bushpig), 6X Plantigrade ungulates (**61**: Hippo, **62**: Elephant), 7X: Bovinae (**71**: African Buffalo), 8X: Equidae (**81**: Plains | Distance (m) from track for direct observations, or age for tracks and signs: **1**: Active, **2**: Recent, **3**: Old, **9**: Not determined or recorded |

| | | | |
|---|---|---|---|
| | | Zebra), 9X: Neotragini (**91**: Suni), 10X: Cephalophini (**101**: Common Duiker, **102**: Red Duiker), 11X: Aepycerotini (**111**: Common Impala), 12X: Madoquini (**121**: Dikdik), 13X: Raphicerini (**131**: Sharpe's Grysbok) | |
| **24**: Wild Carnivores | 1: Direct sighting, 2: Tracks, 3: Spoor, 4: Other signs (Explain in notes) | 1X: Pantherinae (**11**: Lion, **12**: Leopard), 2X: Felidae (**21**: African wild cat), 3X: Viverridae (**31**: African Civet, **32**: Cape Genet), 4X: Hyaenidae (**41**: Spotted Hyena), 5X: Canidae (**51**: African wild dog, **52**: Side - Striped Jackal), 6X: Herpestidae (**61**: Slender Mongoose, **62**: Banded Mongoose **63:** Water Mongoose, **64:** White Tailed Mongoose), 7X: Mustelids (**71**: Honey Badger), 8X Lutrinae ( **81**: African Clawless Otter) | Distance (m) from track for direct observations, or age for tracks and signs: **1**: Active, **2**: Recent, **3**: Old, **9**: Not determined or recorded |
| **25**: Wild Primates and Prosimians | **1**: Direct sighting, **2**: Tracks, **3**: Spoor, **4**: Other | **1**: Lesser Galago, **2**: Greater Galago, **3**: Baboon, **4**: Vervet, **5**: Blue Monkey | Distance (m) from track for direct observations, or age for tracks and signs: **1**: Active, **2**: Recent, **3**: Old, **9**: Not determined or recorded |
| **26**: Rodents | **1**: Direct sighting, **2**: Tracks, **3**: Spoor, **4**: Other | 1: Hystricidae (**11**: Porcupine, **12**: Giant Cane Rat) | Distance (m) from track for direct observations, or age for tracks and signs: **1**: Active, **2**: Recent, **3**: Old, **9**: Not determined or recorded |
| **27:** Macroscelidea | **1**: Direct sighting, **2**: Tracks, **3**: Spoor, **4**: Other | 1: Rhynchoninae (**11:** Sengei) | Distance (m) from track for direct observations, or age for tracks and signs: **1**: Active, **2**: Recent, **3**: Old, **9**: Not determined or recorded |
| **28:** Tubulidentata | **1**: Direct sighting, **2**: Tracks, **3**: Spoor, **4**: Other | 1: Orycteropodidae (**11:** Aardvark) | Distance (m) from track for direct observations, or age for tracks and signs: **1**: Active, **2**: Recent, **3**: Old, **9**: Not determined or recorded |
| **3X**: Summary impressions of land cover attributes averaged over entire segment. | | | |
| **31**: Rice farming | | | **1**: 0%, **2**: 1 - 25%, **3**: 26 - 50%, **4**: 51 - 75%, **5**: 76 - 100% |
| **32**: Other Tillage Crops | | | **1**: 0%, **2**: 1 - 25%, **3**: 26 - 50%, **4**: 51 - 75%, **5**: 76 - 100% |
| **33**: Grass and forb height | | | **1:** <1ft, **2:** 1 - 3ft, **3:** 3 -6ft, **4:** 6ft+ |
| **34:** Visibility of obstruction by Shrubs, Bushes and Trees | | | **1**: 0%, **2**: 1 - 25%, **3**: 26 - 50%, **4**: 51 - 75%, **5**: 76 - 100% |
| **35:** Ground Type | | | **1**: Hard & Dry, **2:** Sand**, 3:** Damp Soil, **4**: Mud, **5:** Water**, 6:** Rock**, 7:** Gravel & Pebbles |
