## Supplementary material for "Blood host preferences and competitive inter-species dynamics within an African malaria vector species complex inferred from signs of animal activity around aquatic larval habitats": S9_Appendix

**S9 Appendix: Conceptual framework for regression analysis of the association between *Anopheles gambiae* sibling species composition and indicators of activity by diverse mammalian species that could act as potential sources of blood for mosquitoes.**

The PCR results obtained from adult $F_0$ mosquitoes that were reared from wild-caught $F_0$ larvae demonstrated that in addition to *An. arabiensis*, wild populations near or inside NNP also included *An. quadriannulatus*. While performing initial GLMMs to assess the potential effects that may influence the relative abundance of each sibling species at each camp, it was evident that the environmental and spatial variables included (Subjective Natural Ecosystem Integrity Index (SNEII), distance to nearest park boundary or human settlement and historical landcover) covaried with the normalised indicators for the number of detections for each species, collected during the radial surveys of animal activity around each camp (1)(S8 Appendix). Although SNEII, distance and landcover were important to include in the occupancy analysis, to ensure that as many environmental drivers of larval occupancy as possible were accounted for, their potential for both direct, proximal causal effects upon *An. gambiae* complex species composition and more indirect, distal causal effects via direct effects upon potential host availability required careful consideration. This is because these fundamental ecological parameters also directly mediate the availability and distribution of the various mammals that may act as potential blood sources for mosquitoes. The conceptual analytical framework developed to guide this analysis (Figure S9), therefore illustrates how these three geographic variables may be reasonably expected to, directly and indirectly, influence proportional species composition of *An. gambiae* complex mosquitoes across the environmentally and ecologically diverse study area, serving as the foundational reasoning for how the generalized liner mixed model (GLMM) analyses were conducted.

Firstly, historical landcover and distance to the nearest NNP boundary or settlement inside the park (S6 Appendix) may have substantive direct abiotic effects on mosquito population composition (Figure S9). The purpose of adding landcover as an additional variable was to account for any potential variance attributable to the environmental factors, that are associated with each of the strikingly different landcover types across the study area, and may well influence occupancy and species composition. For example, it was observed that evergreen groundwater forest, miombo woodland and acacia savannah, were all associated with different soil types and different levels of canopy cover. These natural differences between landcover types may therefore impact environmental factors such as the level of shade or predominant surface water characteristics, all of which may influence the availability, attractiveness and productivity of aquatic habitats (2-4). Furthermore, climatic differences between the landcover types, in terms of temperature and rainfall, for example, may also affect species composition as one particular species may be more tolerant to certain conditions compared to the other (5, 6).

Also, mosquito dispersal is influenced by the geographic distribution of natural resources, including blood hosts and nectar, resting sites, mates, and aquatic habitats (7), so the distribution and gene flow of populations are directly affected. Hence, the addition of distance (S6 Appendix) to the list of potential independent variables considered for inclusion in the GLMMs, to account for the direct effects of dispersal (7) away from the preferred human and cattle hosts of *An. arabiensis* upon its relative abundance with respect to *An. quadriannulatus*. Additionally, the distance variable may also address any variance in population composition that may be attributable to long-distance windborne dispersal (7) of *An. quadriannulatus* away from its optimal ecological niche.

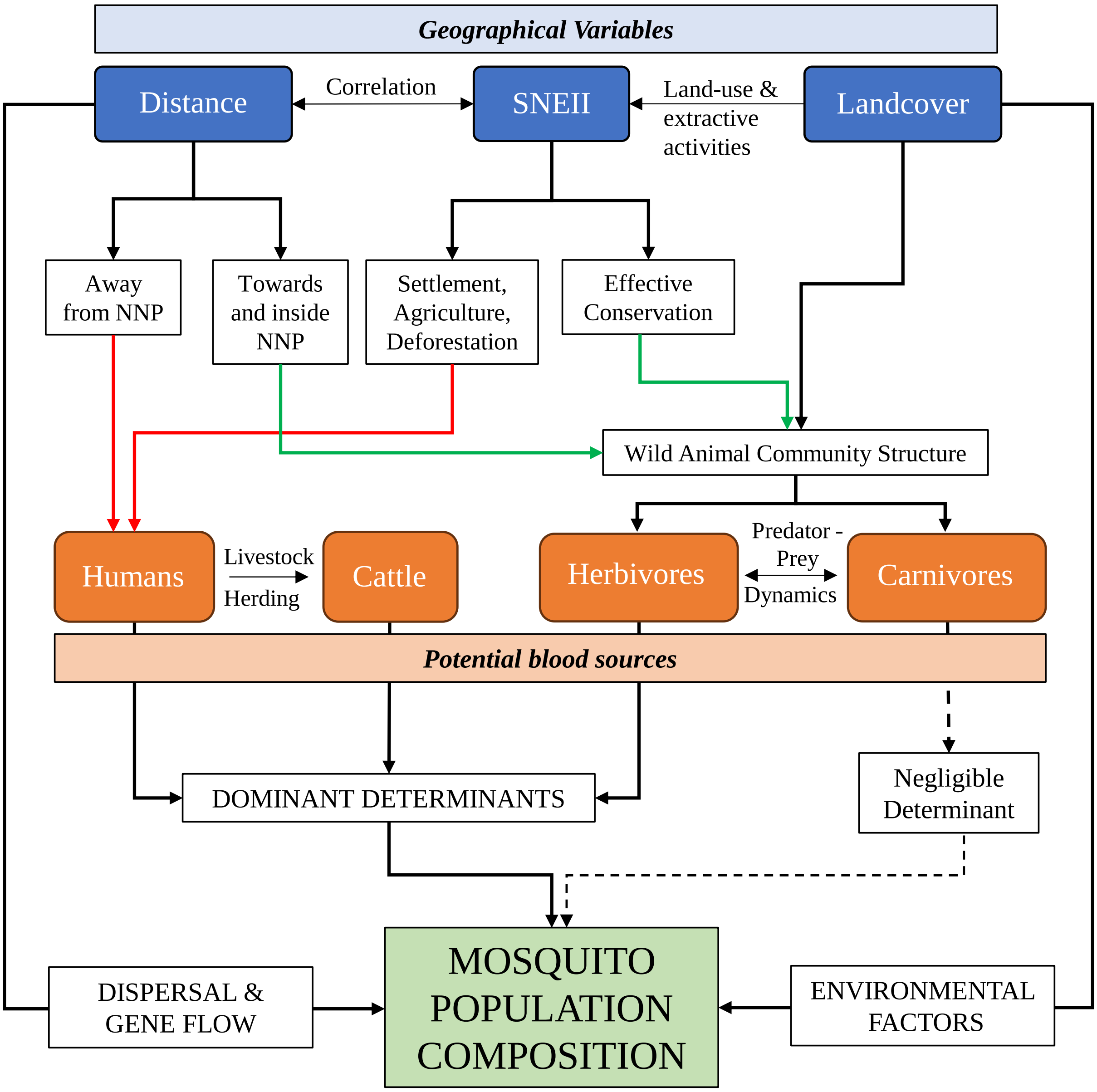


**Figure S9.** A flow diagram illustrating the direct effects of distance, SNEII and landcover on mosquito population composition, and the indirect effects that are mediated through the potential availability and distribution of blood sources.

However, although location, ecosystem integrity and landcover may all have direct effects on mosquito population composition (Figure S9), these spatial, geographic and environmental variables also fundamentally influence how animal communities are structured, which in turn drives the abundance and distribution of the numerous different mammalian species that may act as potential blood sources for different mosquitoes with different host preferences (8-10). With respect to landcover, some species would be more frequently associated with a certain landcover type as it provides a preferred habitat given the availability of natural resources that are required by that species. For example, it was visually observed during the study, and consistent with the literature (11) that impala was almost exclusively associated with dry acacia savanna, while buffalo were predominantly associated with moist miombo woodlands and red duiker were closely associated with dense groundwater forest.

Historical land cover can also influence the SNEII, as certain types of predominant vegetation are associated with different human activities that have distinctive effects on the intensity of land degradation, which was the primary criterion used by the investigators to assign the SNEII scores. For example, miombo woodland provides more suitable environmental conditions for agriculture than acacia savanna, and is therefore more likely to become badly deforested. Also, miombo trees were also regularly observed to be exploited for charcoal burning during the study, especially in the dry season. On the other hand, the groundwater forest provides ample opportunity for selective harvesting of hardwood trees for high value timber and were, therefore, often deforested less drastically than miombo woodlands, and were far less vulnerable than dry acacia savannah where trees of any kind are scarce. All these intricacies, in turn, impact the densities and distributions of humans, cattle, and different wild animals, thus creating a highly heterogeneous landscape with respect to the preferred blood resources each mosquito species depends on.

Furthermore, the distance into or away from the protected area of NNP has obvious direct effects on SNEII and both are, in turn, direct determinants of mammalian community composition, creating a complex web of causality and correlation (Figure S9). Indeed, this study area was specifically chosen due to the clear geographic gradient which runs from west to east, representing a complete transition from fully domesticated land use, to a buffer zone where humans, livestock and wildlife are all present, to wild conserved areas devoid of any humans or livestock. As the most important and primary criterion for the assigned SNEII values, land degradation due to human activity has negative consequences for wildlife communities and is also associated with the presence of humans and livestock (1, 12). Therefore, the furthest distances away from NNP and the lowest SNEII scores occurred at fully domesticated camps, where humans and livestock were clearly the most abundant host species available and so may influence the species composition to favour mosquitoes that prefer these hosts as blood sources. Similarly, as one moves eastwards away from these domesticated landscapes into ILUMA WMA and then NNP, where the natural ecosystems are increasingly intact, animal communities clearly become more abundant and diverse, providing a variety of alternative blood sources that mosquitoes could potentially exploit, thus affecting mosquito species composition.

It was therefore considered that the numbers of detections of humans, livestock and wildlife would be closely correlated with all three of these geographic variables, thus confounding conventional multivariate GLMM analyses of all these variables together, by precluding meaningful model identification and interpretation because both levels of the causal chain illustrated in figure S9 would be included at the same time. In view of this, a separate model was developed that used only these broadly geographic factors to address the fundamental environmental and spatial effects on *An. gambiae* complex species composition across the study area (S10 Appendix), before then investigating how variations in the mosquito population composition were associated with the availability of individual potential host species (S10 Appendix). If the multivariate host species models of humans, livestock and wild animals had also included the effects of distance, SNEII or landcover, it was considered likely that these covariates would have absorbed much of the variance that would otherwise be attributed to the preferred host species of mosquitoes. This could then readily result in the direct causal effect of a given host species being underestimated or even not identified.

Assuming that wild populations of *An. gambiae* complex survive in the absence of humans and livestock, and must presumably do so by acquiring blood meals from wild herbivores, the predator-prey interactions of wild animals in the study area were also considered when building the models of sibling species composition as a function of the availability of individual host species. Large carnivores commonly recorded in this study area included lion, hyena and leopard (Figure S8.8), and it was assumed that their abundance was associated with that of herbivores for a number of reasons. The population ecology of these carnivores, including their distribution and frequency with which they visit particular locations, are influenced by the availability and distribution of prey (13-18). However, these interactions are highly dynamic, and so the presence of carnivores can also deter herbivores that subsequently avoid certain ambush sites and locations where predators are perceived to be present (18-20). Because of this complex web of causality and covariance (Figure S9), and also because carnivores constitute a negligible fraction of overall mammalian blood mass when herbivores are abundant (21) (Figure S9), the numbers of detections of these large carnivores were also omitted from the multivariate host association model, even though they were all clearly associated with *An. gambiae* complex sibling species composition in univariate regression analysis (Table 2). Descriptions of how the GLMMs were fitted, interpreted, and displayed for *An. gambiae* complex sibling species composition based on (1) distance, SNEII, and landcover, and (2) the number of detections for each animal species, are described in S10 Appendix.
