## Supplementary material for "Blood host preferences and competitive inter-species dynamics within an African malaria vector species complex inferred from signs of animal activity around aquatic larval habitats": S10_Appendix

### S10 Appendix: Data entry, cleaning, preparation and analysis

#### *Data entry, cleaning and preparation.*

Three complete datasets including data from the larval occupancy surveys (S3 Appendix), the PCR results from collected field-identified *An. gambiae* complex larvae (S3 Appendix), and the corresponding data from the radial surveys involving humans, livestock, wildlife activity and land use (1)(S8 Appendix), were all required for the full and final statistical analysis. Recorded observations in the larval survey forms (Table in S3 Appendix) were entered into Microsoft Excel® directly after each survey was completed and cleaned using R Version 4.3.1 at the end of rounds 3 and 4 respectively, largely by examining frequency tables broken down by date for implausible values. As the data were directly entered in a horizontal format with observations for each habitat represented by a single line of data with multiple variables (S11 Data), no further restructuring was required for the larval occupancy analysis. These data were linked using Microsoft Access® by camp name and number to a table with the relevant spatial and geographical variables for each camp location, namely the Subjective Natural Ecosystem Integrity Index (SNEII), distance and landcover (S6 Appendix), to form the final dataset presented in supplementary file 11.

To validate the field identification methodology (S4 Appendix) and to calculate the proportional species composition of field-identified *An. gambiae* complex larvae, the PCR amplification results for each camp from the fourth round of surveys were aggregated by breeding site identification number (BSID) to give the total number of *An. arabiensis*, the total number of *An. quadriannulatus*, and the total number of unamplified larvae for each batch sample of field preserved larvae from specific individual aquatic habitats. The total number of larvae per habitat-specific batch sample at a given camp was calculated by summing all amplified and unamplified larvae for each BSID, while the proportion confirmed to be members of the *An. gambiae* complex was calculated by summing the total number of *An. arabiensis* and the total number of *An. quadriannulatus* identified and dividing by the total number of larvae tested. This dataset was then linked by camp name and number to SNEII, distance and historical landcover (S12 Data). To plot the results of the larval validation (Figure in S4 Appendix), and the effect of spatial and geographic parameters on proportional sibling species composition (Figure 4), the data set was aggregated by camp number so that each row contained the sum of *An. arabiensis,* the sum of *An. quadriannulatus,* the number of unamplified larvae, the total number of larvae per batch sample and the total number identified as *An. gambiae* complex.

Data from the 40 radial surveys of human, livestock, wildlife activity and land use across each camp location from the fourth round of surveys in 2023 were cleaned, restructured and linked to the existing radial survey data, using open software R Version 4.3.1, Microsoft Excel®, and Microsoft Access®, exactly as described elsewhere (1, 2), so that each of the many specific types of detections (e.g., old hartebeest spoor, direct sighting of a charcoal burner, recent cattle tracks, etc.) at each camp was represented by a separate variable. However, the complete dataset that entailed of recorded detections from 116 radial surveys across the whole study period from January 2022 to July 2023 was used for the PCA to derive the Objective Natural Ecosystem Integrity Index (ONEII; S6 Appendix [1, 3]). A subset was also created of all 40 radial surveys from round four only and new variables for the total observations of each species was derived by summing every observation count and class for each species. Total detections for each species were normalised using z-score means: $z = \frac{x-\mu}{\sigma}$, where z is the total number of detections of any kind for a given species at each camp, x is the number of detections for a species at each camp, $\mu$ is the mean number of detections for the same species across all camps, and $\sigma$ is the standard deviation of the mean number of detections for that species across all camps. This normalised indicator of the number of detections for each species at each camp was then linked with the PCR results that had been aggregated by BSID and already merged with the spatial and geographic parameters, forming the final dataset presented in S12 Data.

***Statistical analysis of occupancy of aquatic habitats by An. gambiae complex larvae***

The *mgcv* package and *gamm* function were used to fit generalized linear mixed models (GLMMs) to the occupancy dataset (S11 Data), examining the dependence of the proportion of aquatic habitats occupied by *An. gambiae* complex larvae upon various biotic and abiotic factors, using a systematic forward-step selection process. Camp number was added as a random effect to account for covariance of occupancy status across potential larval habitats within the area around individual surveyed camps, and random variance that may occur between camps and is not explained by any of the fixed effects examined and was consistent across survey rounds. Temporal autocorrelation was accounted for by including the number of weeks since the study commenced as a first order continuous autoregression term nested within the camp number random effect. This also helped mitigate the tendency of the rolling cross sectional study design to spuriously exaggerate the significance of various fixed effects examined because of the temporal and environmental covariance between surveys within a circuit which were completed in sequence at similar times and places. As *An. gambiae* complex larvae were recorded as either present or absent in a habitat, this binary dependent variable was fitted to a binomial distribution with a logit link function.

Univariate analysis was initially completed with all the temporal, environmental, abiotic, and biotic parameters (S12 Data) included in separate models as the sole independent variable with a fixed effect. These parameters included all the attributes that were recorded in the larvae survey form (Table in S3 Appendix), SNEII, distance to the nearest NNP boundary or settlement inside the park and historical landcover type. The effect of SNEII, the number of dips and distance to the NNP boundary or settlement inside the park were assessed as continuous fixed effects, whereas all other variables including, individual implementing the survey, season, landcover, habitat type, perimeter, water depth and vegetation type were treated as categorical factors. The reference group for a categorical variable was assigned to the level that was most frequently recorded, except for the covariate, round, in which round 4 was treated as the reference group because it represented all surveys carried out by a different individual who clearly detected larvae with greater sensitivity than the investigator who preceded him over the first three rounds.

For the habitat type variable, waterholes and rice paddies were pooled together as a single category as they had statistically indistinguishable occupancy rates and were often similar in appearance. Streams were combined with springs and swampy areas based on similar criteria. All other habitat types that were not significantly different or were too scarce for which there were too few observations to assess as a separate category, were pooled into the selected reference group of pools, puddles, tracks and depressions, and categorised as *other habitat types*. The new reference group included all the other possible anthropogenic habitat types; artificial ditches or drains, human-dug wells, artificial containers, ridge and furrow agriculture, and other agriculture. These man-made habitats also had similar attributes to the habitats as the original reference group. The category flooded valleys was redundant due to small sample size (n=7) and therefore, was also added to the new reference category. This reduced the number of habitat types by from 12 categories (Table in S3 Appendix) to now three categories for GLMM analysis. The habitat perimeter variable was also recategorized by pooling the two larger groups (*21-200m* and *>200m*) together to form the category *>20m*, which reduced the original four categories (Table in S3Appendix) to three. The Akaike information criterion (AIC) score was used to assess the goodness of fit of each model and the *anova* function was used to test for statistically significant differences between the alternative model fits.

The final multivariate model was then assembled by adding independent variables in order of their significance in the univariate analysis results, with variables being removed again based on the principle of parsimony and lack of evidence for improvements in the goodness of fit. For example, if a variable was statistically significant in the univariate analysis but lost significance or didn't improve the goodness of fit when added to a model, it was dropped from the model and excluded from further consideration. The final model only included the attributes that remained statistically significant at the end of the model building process.

The results of the GLMMs were expressed as odds ratios (OR) calculated as $e^{\beta_i}$ , where $\beta_i$ is the fixed effect in question. For categorical variables, the OR of $\beta_i$ was compared against the reference group for that variable. The OR for distance was calculated as $e^{10\beta_i}$, to represent the change in odds of occupancy for every ten kilometres further towards or beyond the NNP boundary. The OR of SNEII was expressed as $e^{100\beta_i}$ to represent the change in occupancy when moving from a fully domesticated ecosystem to fully intact, natural ecosystem. The 95% confidence intervals for all these ORs were calculated as $e^{\beta_i \pm 1.96*\sigma}$, where $\beta_i$ is the fixed effect in question and σ is the standard error of the mean of the given fixed effect.

Variations in the proportion of habitats occupied by *An. gambiae* complex larvae in relation to variation in SNEII, were represented graphically as a scatterplot using the *ggplot2* and *scales* packages in R. The data were aggregated by date, so that each data point on the scatterplot would represent the proportion of occupied habitats for one whole larval survey on a given day. The predicted means and confidence intervals for the linear trends observed were calculated based on a GLMM identical to the final best-fitting multivariate model except that all fixed effects other than SNEII were treated as random effects. Using the outputs from this GLMM, the predicted means and confidence intervals for plotting were then calculated based on the same formulae used for plotting in S8 Text, where $\beta_0$ is the intercept, $\beta_1$ is the effect size attributed to SNEII, and $x_{1,i}$ represents the values of SNEII at camp $i$, and $\sigma_0$ and $\sigma_1$ are the standard errors of the means for $\beta_0$ and $\beta_1$, respectively.

***Statistical analysis of effects of distance, ecosystem integrity and landcover on An. gambiae complex species composition***

Following revision of the field procedures to ensure samples of larvae caught at all camp locations were preserved *in situ* (S3 Appendix) and using the final linked dataset (S12 data), GLMMs were also fitted with logit link function to a binomial distribution, treating the dependent variable as the proportion of *An. arabiensis* rather than *An. quadriannulatus,* weighted according to the total number of amplified *An. gambiae* complex specimens identified. The BSID nested within camp number was treated as a random effect to account for any covariance within habitats and camp locations. Univariate analysis of distance, SNEII and historical landcover type were initially completed and the best model fit was derived by the same algorithm based on the principles of parsimony and goodness of fit as described above.

The proportion of *An. arabiensis* at the mean camp according to the fixed effects in the best fit multivariate model was calculated as $\frac{e^{\beta_0}}{1+e^{\beta_0}}$ , where $\beta_0$ is the intercept of the model. The ORs and 95% CIs for the univariate and multivariate analyses were expressed for distance and landcover as described in the section above. The proportion of *An. arabiensis* rather than *An. quadriannulatus* were aggregated by camp to generate a scatter plot where the vertical axis represents the proportional species composition, and the horizontal axis represents the distance variable. The predicted means (A) and standard error of the means (σ), with 95% CIs for the proportion of the PCR-identified members of the *An. gambiae* complex that were *An. arabiensis,* were plotted using the formulae $A = \frac{e^{\beta_0+\beta_1 X_{1,i}+\beta_2 X_{2,i}+\cdots\beta_n X_{n,i}}}{1+e^{\beta_0+\beta_1 X_{1,i}+\beta_2 X_{2,i}+\cdots\beta_n X_{n,i}}}$, and $\sigma = \sqrt{{\sigma_0}^2 + \sigma_1 {x_{1,i,}}^2 + \sigma_2 {x_{2,i,}}^2 + \cdots \sigma_n {x_{n,i,}}^2}$, respectively, where $\beta_0$ is the intercept, $\beta_{1,}$ $\beta_2$ ..…$\beta_n$, are the effect sizes attributed to the fixed effects included in the best fit multivariate model, and $x_{1,i}, x_{2,i}, \ldots x_{n,i}$ represents the values of each respective fixed effect at camp $i$, and $\sigma_0$, $\sigma_1$ ... $\sigma_n$ are the standard errors of the means for $\beta_0$, $\beta_0$ ... $\beta_n$, respectively. The 95% CI for the predicted means at each camp were then calculated as $G = \frac{e^{\beta_0+\beta_1 X_{1,i}+\beta_2 X_{2,i}+\ldots\beta_n X_{n,i}\pm 1.96*\sigma}}{1+e^{\beta_0+\beta_1 X_{1,i}+\beta_2 X_{2,i}+\ldots\beta_n X_{n,i}\pm 1.96*\sigma}}$. The predicted means and confidence intervals were then plotted using the *geom_line* and *geom_ribbon* functions, respectively.

***Statistical analysis of associations between An. gambiae complex species composition and potential host availability***

A second set of GLMMs of PCR-identified specimens that were identified as *An. arabiensis* rather than *An. quadriannulatus,* were fitted to investigate the effects that humans, cattle, and various wild mammals that might act as potential blood sources, might have on population composition of this mosquito species complex. Using the dataset provided in supplementary file 11, BSID nested within camp number were again treated as random effects and the spatial and geographic parameters from the best fit GLMM as described in the section above were omitted based on a logical conceptual framework (S9 Appendix). A Pearson's correlation test was completed to assess the covariance between detected activity levels of humans and cattle (S7 Figure), motivating the calculation of a new variable by summing the total numbers of detections of humans and the number of total detections of cattle herds. The new variable was then normalised as z-score means exactly as it had been done for the total number of detections for a given mammalian species at each camp. Univariate outputs of the total detections of humans, the total detections of cattle, and the combined new variable were compared in terms of goodness of fit, and the latter was used for multivariate analysis because it yielded the lowest AIC scores. The normalised total detections for all other species were individually assessed as the sole fixed effects in univariate models, and the same forward-step selection process as described for the occupancy model was completed to identify the best-fit multivariate model.

Univariate and multivariate results of the species of interest were calculated exactly as previously described. Note, however, that they need to be interpreted differently because the OR reflect the change in the odds of a specimen being *An. arabiensis* rather than *An. quadriannulatus,* per increase of one standard deviation above the mean number of detections of the animal species in question. The predicted means and confidence intervals for the relative abundance of *An. arabiensis* at each camp that were based on the normalised total detections of the animal species from the best model-fit, were calculated using the same formulae as described in the section above and were graphically displayed on the same horizontal and vertical axes.
